# A cytoskeletal toggle switch controls amnion architecture and fate

**DOI:** 10.64898/2026.11.13.766348

**Authors:** Chloé Roffay, Oliver M. Drozdowski, Sarah Jay, Cindy Horenburg, Maria Novatchkova, Elias Janscht, Alberto Moreno-Cencerrado, Nicolas Rivron, Édouard Hannezo, Diana Pinheiro

## Abstract

Binary cellular decisions are generally attributed to gene regulatory networks, with cytoskeletal-driven shape changes often treated as an output. Yet, whether the cytoskeleton can itself implement switch-like logic to drive abrupt morphological transitions remains unknown. In the primate amniotic sac, the extraembryonic amnion derives directly from the columnar epiblast and abruptly flattens into a squamous epithelium to form a highly asymmetric cavity. Here, we identify a cytoskeletal switch driving this developmental bifurcation in human gastrula models. Mutual inhibition between actomyosin and keratin networks generates two mechanical attractors: amnion cells flip into a keratin-rich, squamous shape, while the epiblast remains myosin-rich and columnar. Ectopic activation of YAP1-TEAD signaling triggers the myosin-keratin switch without inducing amnion specification, establishing that the switch is not simply a consequence of fate changes. Conversely, preventing the switch disrupts amnion specification, showing that this shape bifurcation gates lineage commitment. Single-cell transcriptomics across cytoskeletal perturbations revealed a transcriptional signature for the toggle switch, which is recapitulated *in vivo* in the human and marmoset amnion. Together, our findings show that the cytoskeleton acts not only as an effector of tissue shape but as a decision-making module, implementing the same toggle switch logic as gene regulatory networks to drive morphological bifurcations.

## Introduction

A primary goal of development is to generate highly specialized tissue architectures that enable biological function. In epithelia, cell shape is closely tied to function: squamous cells permit rapid diffusion across lung alveoli, cuboidal cells support secretion and absorption in glandular ducts, and columnar cells sustain active transport in the gut while building a robust barrier^1^. Although extensive work revealed a central role for cytoskeletal-generated forces in sculpting tissues into form^1–4^, how these molecular inputs robustly generate, and maintain, a specific set of shapes remains unclear. Indeed, in several systems, cells were shown to adopt a small number of discrete shapes, with sharp transitions pushing them between distinct morphologies^5–10^. How discrete cellular states arise has been addressed largely in a different context, namely cell fate specification, following the metaphor of Waddington’s epigenetic landscape^11–14^. Gene regulatory networks (GRNs) are well-understood to enable binary fate decisions, via recurring network motifs. For instance, mutual repression between transcription factors is a simple design principle allowing cells to commit to only one of two possible states^11–15^. Within this framework, cell shape is often treated as a downstream product of fate specification. Although cytoskeletal forces are well-established regulators of cell state^4,16^, whether the cytoskeleton can itself encode a discrete switch between tissue architectures remains unknown.

Having evolved independently in insects and amniotes as a convergent solution to protect and support the embryo, extraembryonic tissues are a prime example of sharp morphogenetic transitions, often flattening into expansive epithelial sheets during early development^17,18^. The amnion is a thin extraembryonic membrane that surrounds the embryo throughout gestation and functions as the dominant load-bearing layer of the foetal membranes^17–23^. In primates, the amnion derives directly from the pluripotent epiblast^20–22,24,25^, yet the two tissues must adopt highly divergent squamous and columnar shapes — bifurcating from a shared origin to give the amniotic sac its asymmetric form^19,21,22,26–31^. This divergence is one of the earliest fate decisions, and shape changes, in the post-implantation human embryo, occurring even before the primitive streak is morphologically apparent^32^. Because the primate amnion differs from mouse in developmental origin, timing of formation and the GRNs controlling its identity^33–38^, how the sharp morphogenetic bifurcation between epiblast and amnion is controlled and coordinated with fate specification has remained largely unknown.

To tackle this question, we took advantage of a widely used *in vitro* model recapitulating the signalling events and transcriptional signatures of the early human gastrula: 2D gastruloid discs^39–43^. Upon stimulation with BMP4, these micropatterned human pluripotent stem cell (hPSC) colonies self-organize into concentric rings of the germ layers (ectoderm and mesendoderm) surrounded by extraembryonic amnion-like cells, expressing GATA3, TFAP2A, CDX2 and ISL1 (Extended data figure 1a)^43–46^. Combining biophysical modelling, direct force measurements and functional perturbations, we show that double negative feedback between the actomyosin and keratin networks forms a cytoskeletal toggle switch that generates two mechanical attractors. Amnion cells flip into a high-keratin/low-tension state, and thus adopt a squamous shape, while the columnar-shaped epiblast remains in a high-myosin/high-tension state. Ectopic activation of YAP1 is sufficient to trigger the cytoskeletal toggle switch even in the absence of fate commitment. In turn, blocking YAP1-TEAD signalling, or the myosin-keratin switch disrupted both amnion architecture and fate, supporting that shape gates lineage commitment in the amnion. Finally, single-cell transcriptomics across cytoskeletal perturbations identified a transcriptional signature for the cytoskeletal toggle switch, which is recapitulated in human and marmoset embryos *in vivo*, and validated in non-human primate stem cell models. Together, these findings identify a mechanical counterpart to the classic ‘toggle switch’ logic that drives sharp morphogenetic transitions during early primate development.

## Results

### Sharp columnar-to-squamous transition in the amnion is cell-autonomous

To test whether gastruloid discs recapitulate the morphometric signatures of the early human gastrula *in vivo*, we established an automated image analysis pipeline for segmentation-based extraction of cell shape and tissue architecture. First, gastruloid discs were immunostained for markers of pluripotent epiblast/ectoderm (SOX2), primitive streak/nascent mesoderm (BRA), and amnion (CDX2), together with the tight junction protein ZO-1. High-resolution images were then acquired, automatically projected, and cell outlines segmented to extract apical cell area and height across the gastruloid disc radii (Extended data figure 1b-d, see Methods for detailed description), thus providing the first quantitative morphological atlases in human gastrula models.

Pluripotent discs, stimulated with BMP4 ligands for only 1 hour (h), displayed a homogeneously columnar organization, with comparable apical area and height along the disc radii (Figure 1a-d, Extended data figure 2a-c). By 24 h, apical area began to increase in specifying amnion cells, marked by expression of CDX2, GATA3 and TFAP2A at the gastruloid edge (Figure 1a-d, Extended data figure 2a-c). This apical expansion was accompanied by a local decrease in gastruloid height (Figure 1c,d, Extended data figure 2c), with increasingly pronounced amnion flattening until 48 h (Figure 1a-d, Extended data figure 2c-f). In contrast, the embryonic core, composed of BRA- and SOX2-positive cells, maintained a more columnar organization throughout the differentiation time course, with cells becoming increasingly smaller apically and taller (Figure 1a-d, Extended data figure 2a-f). Importantly, these distinct morphological signatures arise due to BMP4-induced self-organization, since unstimulated micropatterned colonies remained uniformly columnar even after 48 h (Extended data figure 2g-j). Overall, this quantitative analysis supports that, despite their simplified 2D geometry, gastruloid discs recapitulate the distinct architectures of the extraembryonic amnion and epiblast observed in the early human gastrula^19,21,22,26–31,47^.

**Figure 1:**
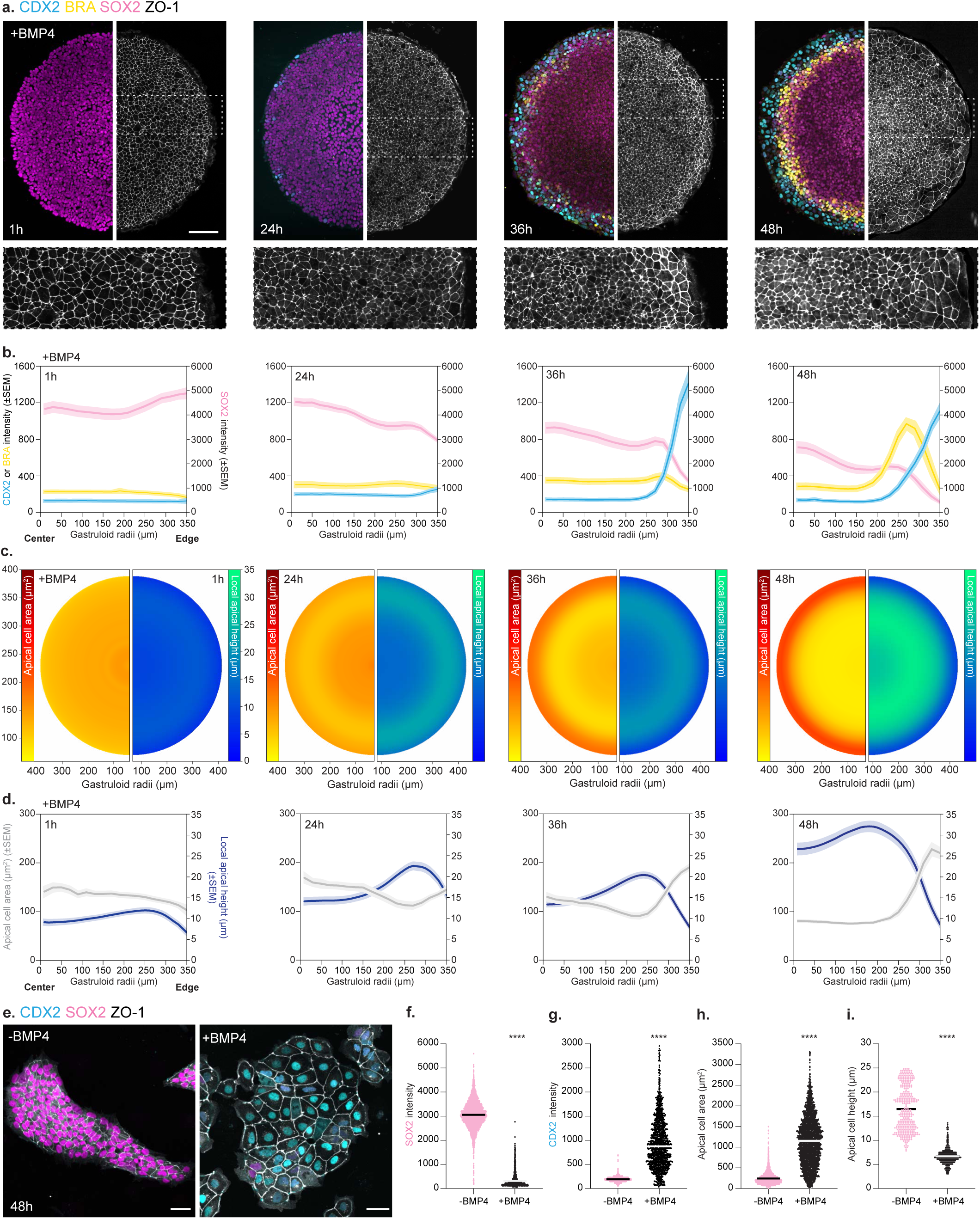
Columnar-to-squamous transition in the amnion is cell-autonomous. **a**. Immunostaining of BMP4-stimulated gastruloid discs, marked for CDX2 (amnion: cyan), BRA (mesoderm: yellow), SOX2 (pluripotent and ectoderm: magenta), and ZO-1 (grey), at 1, 24, 36 and 48h of BMP4 stimulation. Maximum projection is shown for cell fate markers, and the custom workflow was used for the apical surface projection, based on ZO-1 local intensity. Zoom-in images are shown below. **b**. Average intensity of CDX2, BRA and SOX2 as a function of gastruloid radii after 1 (n=45, N=3), 24 (n=30, N=2), 36 (n=48, N=2) and 48h (n=44, N=3) of BMP4 stimulation. **c**. Average atlases of apical cell area (left) and apical height (right) of gastruloid discs after 1 (n=80, N=4), 24 (n=82, N=4), 36 (n=102, N=4) or 48h (n=51, N=4 for apical area; n=62, N=4 for local apical height) of BMP4 stimulation. **d**. Average apical cell area (left axis) and local apical height (right axis) as a function of gastruloid radii after 1, 24, 36 and 48h of BMP4 stimulation (n and N as in C). **e**. Immunostaining of unconfined colonies that were either left untreated (-BMP4, left) or stimulated with BMP4 ligands (+BMP4, right) for 48h, marked for CDX2 (cyan), SOX2 (magenta), and ZO-1 (grey). The corresponding side views are shown in Figure 2j. (**f,g**) Average intensity of SOX2 (f) or CDX2 (g) for unconfined colonies that were either left untreated (n=4541 cells, N=2) or stimulated with BMP4 ligands (n=1279 cells, N=2) for 48h. **h**. Average apical cell area in unconfined colonies that were either left untreated (n=5168 cells, N=2) or stimulated with BMP4 ligands for 48h (n=2402 cells, N=2). **i**. Average apical-basal height in unconfined colonies that were either left untreated (n=360, N=2) or stimulated with BMP4 ligands for 48h (n=464, N=2). Scale bars: 100 μm (a), 50 μm (e). Error bars are shown as mean ± SEM (b,d). Mann-Whitney test. ****P<0.0001 (f-i).

Next, we asked whether amnion architecture is triggered by intrinsic forces or instead set by pulling forces arising in the neighbouring epiblast. To isolate the amnion from the mechanical influence of adjacent embryonic tissues, we stimulated small, unconfined hPSC colonies with BMP4 ligands for 48 h, which were previously shown to uniformly differentiate into amnion (Figure 1e-g)^44,45,48–50^. These unconfined CDX2-positive colonies adopted a fully squamous architecture, with cells displaying large apical areas and small apical-basal height (Figure 1e,h,i). In contrast, unstimulated colonies seeded in a similar manner remained columnar even after 48 h (Figure 1e,h,i). These findings support that amnion cells autonomously control their transition into a squamous, flattened morphology. Such active morphogenesis contrasts with previous work in mouse and quail embryos, in which external forces—such as hydrostatic pressure^51–53^ or cell motility^54^ — were shown to play a key role in driving the squamous architecture of extraembryonic tissues.

### The columnar-to-squamous transition in the amnion is mediated by active wetting

To investigate how amnion-intrinsic forces remodel its architecture, we then turned to biophysical modelling. In 3D bubbly vertex models, an established framework for understanding epithelial architecture^55^, a squamous transition arises from differential tension along the apical-basal axis, namely via i) increased lateral tension or, ii) decreased apical or basal tensions (Extended data figure 3a-h)^56–61^. To examine changes in apical-basal tension experimentally, we reconstructed 3D cell shape in gastruloid discs and automatically extracted the interfacial apical angles^62^ during self-organization. Surprisingly, we found that the interfacial apical angles, and thus the ratio of apical-to-lateral tensions, in the amnion did not markedly change over time (Figure 2a,b), ruling out differential apical or lateral tensions as drivers of the squamous transition. Consistent with this, modelling all scenarios of differential tension along the apical-basal axis yielded only modest cell flattening (Extended data figure 3c-h, see Theory Note for details)^56,57^, far short of the fully flattened architecture of amnion cells observed experimentally (Figure 1c,d, Extended data figure 2f). In search of an alternative biophysical mechanism, we found that modelling a global drop of all interfacial tensions could drive a complete columnar-to-squamous transition in the amnion domain (Figure 2c-e, Extended data figure 3i,j, see Theory Note). This arises in the model, because tension becomes low compared to basal adhesion forces, a phenomenon analogous to the physics of wetting^63,64^ (Extended data figure 3i,j, see Theory Note for details).

**Figure 2:**
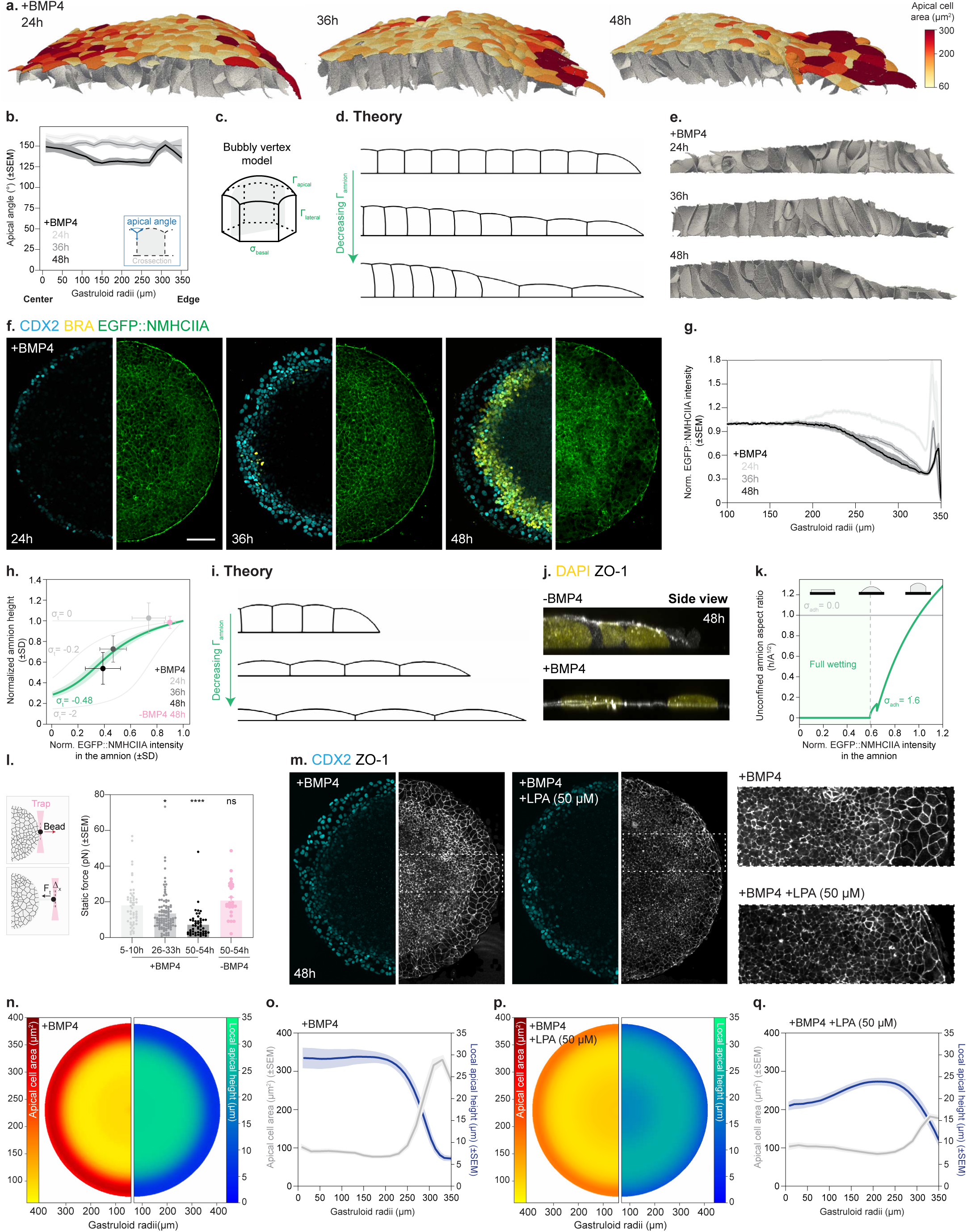
Active wetting mediates the columnar-to-squamous transition in the amnion. **a**. 3D reconstructions of self-organizing gastruloid discs (see Methods for details). Apical view is color-coded for apical cell area and the corresponding legend is shown on the right. **b**. Interfacial apical angles as a function of gastruloid radii, 24 (n=3, N=2), 36 (n=3, N=2) and 48h (n=3, N=2) after BMP4 stimulation. Schematic representation of the method to extract interfacial apical angles (see Methods for details). **c**. Schematic representation of the bubbly 3D vertex model indicating apical, lateral and basal tensions, as well as basal adhesion. The two-dimensional cross-section is highlighted in grey and shown for different parameter values in (d,i). **d**. Bubbly vertex model simulations (cross-sectional 2D model) considering increasingly uniformly lower actomyosin tension at the gastruloid edge. **e**. Cross-section of the 3D reconstructed gastruloid discs during self-organization, showing the lateral interface for direct comparison with model predictions in (d) (see Methods for details). **f**. EGFP::NMHCIIA (green)-expressing gastruloid discs, immunostained for CDX2 (cyan) and BRA (yellow), after 24, 36 and 48h of BMP4 stimulation. Corresponding LifeAct::mRFP expression for these gastruloid discs are shown in Extended data figure 4c. **g**. Normalized intensity of EGFP::NMHCIIA as a function of gastruloid radii after 24 (n=16; N=2), 36 (n=16; N=2) and 48h (n=24; N=2) of BMP4 stimulation. **h**. Height of amnion cells as a function of the fold-change in EGFP::NMHCIIA intensity, predicted theoretically (see Theory Note for details) versus the experimentally determined values for self-organizing gastruloid discs or unstimulated micropatterned colonies at 48h. Green line corresponds to maximum-a-posteriori Bayesian fit of confinement stress, with the associated 95% confidence interval (see Theory Note for details). **i**. Cross-sectional bubbly vertex model simulations of unconfined colonies with uniformly decreasing myosin tension (see Theory Note for details). **j**. Side view of unconfined colonies, immunostained ZO-1 (grey) and nuclei (DAPI, yellow), that were either left untreated (-BMP4) or stimulated with BMP4 ligands (+BMP4) for 48h (n and N as in Figure 1f,g). **k**. Phase diagram of unconfined colony shapes as a function of the fold-change in EGFP::NMHCIIA intensity. This predicts a full wetting transition below a critical tension threshold. **l**. Static force, measured in picoNewtons (pN), at the edge of self-organizing gastruloid discs (5-10h: n=52, N=3; 26-33h: n=101, N=4; 50-54h: n=51, N=4) or in unstimulated colonies (-BMP4 50-54h: n=25, N=2). A schematic of the tether-pulling assay is shown on the left (Extended data figure 4I for a representative curve). **m**. Immunostaining of gastruloid discs treated with BMP4 ligands alone or together with LPA (50 µM) for 48h, marked for CDX2 (cyan) and ZO-1 (grey). (**n,p**). Average atlases of apical cell area (left) or local apical height (right) in gastruloid discs stimulated with BMP4 ligands alone (n, n=22; N=2) or together with LPA (50 µM) (p, n=21; N=2) for 48h. (**o,q**). Average apical cell area (left axis) and local apical height (right axis) as a function of gastruloid radii upon stimulation with BMP4 ligands alone (o) or together with LPA (50 µM) (q) for 48h (n and N as n,p). Maximum projection is shown for cell fate markers, and a custom workflow was used for the apical surface projection of EGFP::NMHCIIA and ZO-1. Scale bars: 100 μm (f,m). Error bars are shown as mean ± SEM for (b,g,l,o,q) and as mean ± SD (h). Kruskal-Wallis test. *P=0.0486. ****P<0.0001. ns: not significant (l).

To investigate this wetting transition in the amnion, we first examined the distribution of key force generators — non-muscle myosin IIA (EGFP::NMHCIIA) and F-Actin (LifeAct::mRFP1)^3,65,66^, using newly established fluorescent reporters (Extended data figure 4a,b). In line with previous work^67,68^, we observed that an actomyosin cable surrounds both BMP4-stimulated and unstimulated colonies (Figure 2f,g, Extended data figure 4c-e). Within the amnion domain, however, both EGFP::NMHCIIA and LifeAct::mRFP1 intensity declined progressively over time (Figure 2f,g, Extended data figure 4c). Importantly, this decrease was observed at both cell-cell and cell-ECM contacts (Extended data figure 4f). Since myosin was shown to be a strong predictor of cortical tension^69–71^, we inputted the experimentally measured EGFP::NMHCIIA fold-change into the analytical model and asked whether this was sufficient to predict amnion shape. We found that this model quantitatively recapitulated the extent of amnion flattening observed experimentally, with confinement stress as the single fitting parameter (Figure 2h, see Theory Note for details). Moreover, based on the levels of basal adhesion inferred in gastruloid discs, modelling the global tension drop in small, unconfined amnion colonies predicted even more marked flattening (Figure 2i; see Theory Note for details), matching our experimental measurements (Figure 1e,h-i, Figure 2j,k). As an orthogonal approach to examine the tensile state of the amnion, we took advantage of the Flipper-TR probe to estimate plasma membrane tension^72–75^, a mechanical parameter known to critically depend on cortical tension^76,77^. Fluorescence lifetime imaging revealed that, soon after BMP4 stimulation (1 h), probe lifetime, and thus tension, were higher at the gastruloid edge than at the centre (Extended data figure 4g,h), in line with a recent report^46^. Importantly, however, this trend was fully reversed by 48 h of BMP4 stimulation (Extended data figure 4g,h), suggesting that tension, just like myosin levels (Figure 2f,g, Extended data figure 4c,f), significantly drop during amnion specification.

To measure the mechanical properties of the amnion, we performed tether-pulling assays using optical tweezers. In this assay, the force exerted on the bead by the membrane tether is a direct readout of membrane tension (Figure 2l, schematic on the left)^78–80^. Consistent with the Flipper-TR measurements (Extended data figure 4g,h) and the wetting model (Figure 2d,e, Extended data figure 3i,j, see Theory Note), we found that tension gradually decreased at the edge of gastruloid discs (Figure 2l, Extended data figure 4i). In contrast, membrane tension remained high in unstimulated micropatterned colonies even after 48 h (Figure 2l), further supporting that the squamous transition in the amnion coincides with a global relaxation in cell surface mechanics. To test these correlations functionally, we then incubated gastruloid discs with lysophosphatidic acid (LPA), a well-established activator of actomyosin tension^81^. Consistent with the model’s prediction (Extended data figure 4l), LPA-treated gastruloid discs, which displayed higher myosin levels at the edge by 48 h (Extended data figure 4j,k), showed impaired amnion flattening (Figure 2m-q).

Finally, we sought to investigate whether the wetting mechanism we uncovered in gastruloid discs could explain amnion morphology *in vivo*. We therefore adapted our model to the 3D geometry of the embryo, comparing active wetting with an alternative model in which increased luminal pressure in the amniotic cavity drives amnion morphogenesis (Extended data figure 4m, see Theory Note for details). Strikingly, even for the same degree of amnion flattening, the two models predicted qualitatively different shapes for the amniotic sac (Extended data figure 4n,o). The pressure model resulted in fully spherical amniotic sacs (Extended data figure 4o), consistent with previous modelling in other systems^53,82^. In contrast, modelling active wetting in the amnion correctly predicted the low curvature observed in the amnion *in vivo* (Extended data figure 4n) and recapitulated the typical slit-shape of the amniotic cavity in CS5-CS6 human embryos^19,21,26–28,47,83^ and *in vitro* 3D models for the human amniotic sac^84,85^.

### Amnion shape bifurcation is implemented through the rewiring of cytoskeleton composition

Next, we asked how the global drop in actomyosin tension in the amnion is molecularly controlled. For this, we first reanalysed an available single-cell RNA sequencing (scRNA-seq) dataset of self-organizing gastruloid discs^39,40^ for UNIPROT categories enriched specifically in the amnion by 44 h of BMP4 stimulation. Keratins, members of the intermediate filament family well-known for their high mechanical resistance to deformation^86–92^ and often expressed in extraembryonic tissues^35,39,92–95^, were amongst the top enriched in the amnion (Extended data figure 5a-c; see Methods for details). While the precise dynamics differed for each keratin (Extended data figure 5f-i), we detected an overall increase in keratin filaments, particularly for the KRT8/18 pair, at the gastruloid edge after 24 h of BMP4 stimulation (Extended data figure 5d-i, Figure 3a,b). By 48 h, amnion cells displayed a dense filamentous keratin network (Figure 3a,b, Extended data figure 5f-i). Interestingly, keratins were recently shown to be required for the formation of the amnion, allantois and blood islands in mouse^95^, yet the underlying mechanistic basis for this requirement has remained unknown.

**Figure 3:**
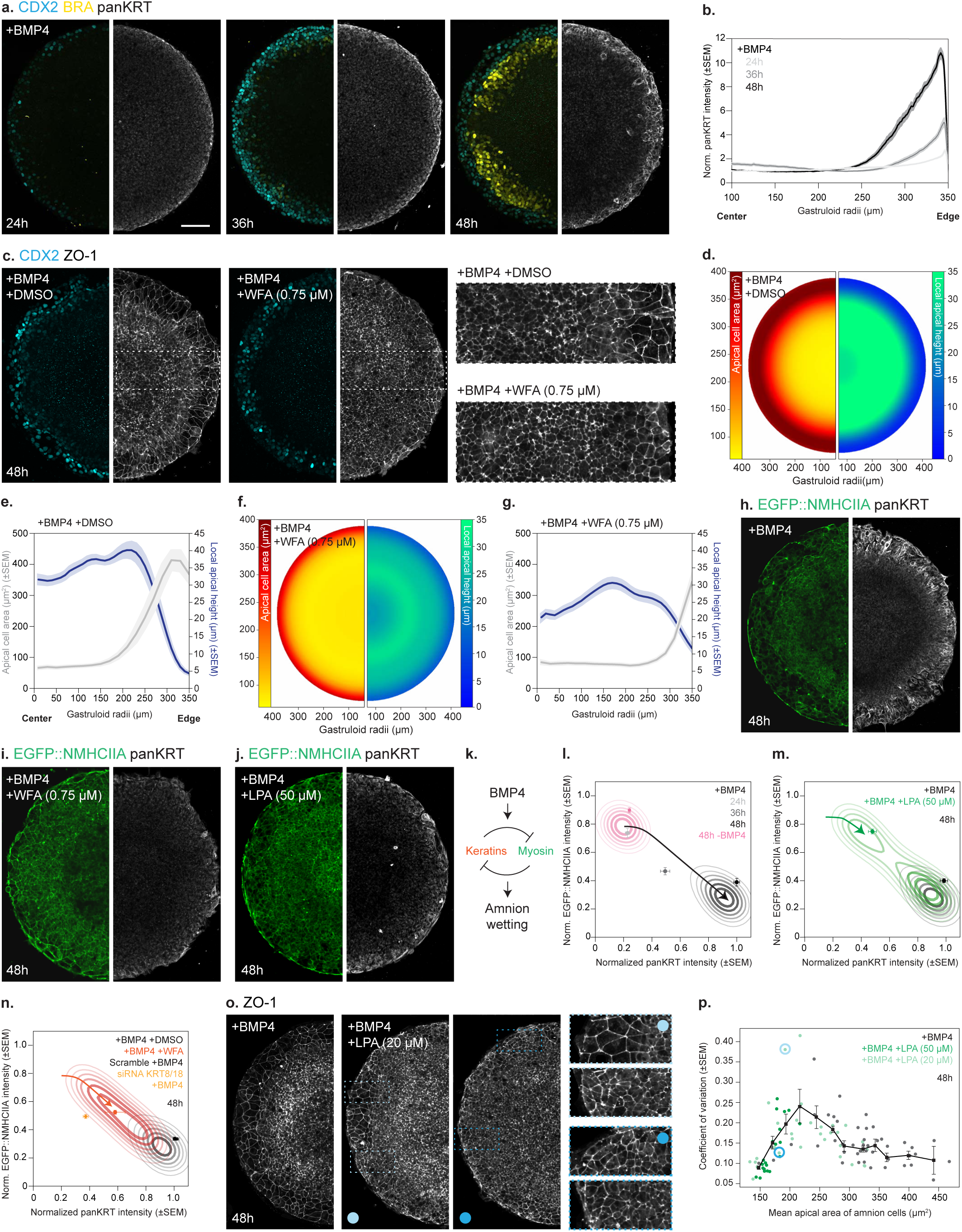
Rewiring cytoskeleton composition, from actomyosin to keratin networks, drives amnion wetting. **a.** Immunostaining of BMP4-stimulated gastruloid discs, marked for CDX2 (cyan), BRA (yellow) and pan-keratin (panKRT, grey) after 24, 36 and 48h of BMP4 stimulation. **b**. Normalized intensity of pan-keratin as a function of gastruloid radii after 24 (n=25, N=3), 36 (n=24, N=3) and 48h (n=24, N=3) of BMP4 stimulation. **c**. Immunostaining of DMSO or WFA-treated gastruloid discs, marked for CDX2 (cyan) and ZO-1 (grey), after 48h of BMP4 stimulation. DMSO and WFA were added from 24 to 48h. Zoom-ins are shown on the right. (**d,f**). Average atlases of apical cell area (left) and local apical height (right) for DMSO (d, n=20, N=3; n=32, N=3) or WFA-treated (f, n=21, N=3; n=33, N=3) gastruloid discs after 48h of BMP4 stimulation. (**e,g**) Average apical cell area (left axis) and apical height (right axis) as a function of gastruloid radii for DMSO (e) or WFA-treated (g) gastruloid discs incubated with BMP4 ligands for 48h (n and N as in d,f). (**h-j**). Immunostaining of pan-keratin (grey) in control (h), WFA (i) or LPA (j)-treated gastruloid discs expressing EGFP::NMHCIIA (green) after 48h of BMP4 stimulation (see quantifications in Extended data figure 4k, Extended data figure 6b,l,o). **k**. Schematic representation of the proposed myosin-keratin toggle switch, where BMP4 signalling favours the keratin-high state, while myosin levels determine the extent of amnion wetting (see Theory Note for details). (**l-n**) Comparison between the landscape derived from the toggle switch model versus the data collected across experimental conditions. Contours represent the landscape derived from the model in different conditions. **l.** Predicted trajectory (black line) between the unstimulated (pink) and the BMP4-stimulated attractor (black) (n and N as indicated in Figure 2g, Figure 3b, Extended data figure 4e and Extended data figure 5e). **m.** Predicted trajectory in LPA-treated gastruloid discs, where the high actomyosin state is stabilized (green line; final attractors in green). Grey contours highlight the BMP attractor as in (l) (n and N as indicated in Extended data figure 4k, Extended data figure 6o). **n.** Predicted trajectory in gastruloid discs where keratin filaments were disrupted, following WFA incubation or in KRT8/18 siRNA-targeted gastruloid discs (orange line; final attractors are shown in orange). Grey contours highlight the BMP attractor as in (l) (n and N as indicated in Extended data figure 6b,f,l,n). **o.** Immunostaining of gastruloid discs treated with BMP4 ligands alone or together with LPA (20 µM) for 48h, marked for ZO-1 (grey). Zoom-ins are shown on the right. **p.** Coefficient of variation of the apical area of amnion cells as a function of the average, calculated for each individual gastruloid across control (n=54, N=3), LPA (20 µM; n=31, N=3) or LPA (50 µM; n=22 N=2). Black line indicates a peak of variability at intermediate values of apical cell area across all conditions. The circles correspond to the coefficient of variation for the two gastruloid discs shown in (o). Maximum projection is shown for cell fate markers and pan-Keratin, while the custom workflow was used for the apical surface projection of EGFP::NMHCIIA and ZO-1. Scale bars: 100 μm (a,c,h-j,o). Error bars are shown as mean ± SEM (b,e,g,l-n,p).

Since keratin upregulation coincided with the global drop in myosin levels (Figure 2f,g), we reasoned that amnion wetting may be implemented through the rewiring of cytoskeleton composition from actomyosin to keratin-based filaments. To directly test this, we incubated self-organizing gastruloid discs with Withaferin-A (WFA), which was previously shown to trigger disassembly of intermediate filaments^96–98^. Gastruloid discs treated with WFA between 24 and 48 h displayed both a marked reduction of keratin filaments (Extended data figure 6a,b) and a defective squamous transition in the amnion (Figure 3c-g). To test the role of KRT8/18 filaments directly, we interfered with their expression using a mix of small interfering RNAs (siRNAs; Supplementary Table 2). This resulted in strong, even if mosaic, downregulation of KRT8/18 proteins (Extended data figure 6c,d) as well as a decrease in total keratin levels (Extended data figure 6e,f). As observed upon WFA treatment (Figure 3c-g), KRT8/18 downregulation compromised the squamous transition in the amnion, resulting in more homogeneous profiles of apical area and height across the disc radii (Extended data figure 6g-k).

To determine whether changes in keratin and actomyosin organization are parallel or coupled events, we then examined actomyosin organization upon WFA or KRT8/18 downregulation and, conversely, keratin organization in LPA-treated gastruloid discs. Strikingly, in WFA or KRT8/18 siRNA-treated gastruloid discs, we detected higher myosin levels at the gastruloid edge compared to controls (Figure 3h,i, Extended data figure 6l-n). Conversely, in LPA-treated gastruloid discs, keratin levels and organization at the edge were significantly reduced (Figure 3j, Extended data figure 6o), consistent with double negative feedback between the actomyosin and keratin cytoskeleton. This interplay corresponds to the simplest regulatory implementation of a toggle switch, a canonical network motif often observed in GRNs to ensure sharp fate transitions^99–101^. Modelling this regulatory logic yielded a landscape with two mechanical attractors and further suggested that initial transcriptional upregulation of the keratin cytoskeleton upon BMP4 stimulation is sufficient to destabilize the myosin-high (keratin-low) attractor in favour of the myosin-low (keratin-high) attractor (Figure 3k,l, Extended data figure 7a-l, see Theory Note for details). The transition between two attractors agreed well with the dynamics of amnion formation in myosin-keratin phase space (Figure 3l), as well as with the perturbation experiments (Figure 3m,n, Extended data figure 7m,n, see Theory Note).

A toggle-switch motif is expected to produce maximal variability in the system’s final state under perturbations of intermediate strength, where the two mechanical attractors can coexist (see Theory Note for details). To test this prediction experimentally, we treated gastruloid discs with lower doses of LPA (20 versus 50 µM). Strikingly, this resulted in highly heterogeneous amnion shapes, both across gastruloids and even within a single disc, with co-existence of regions of columnar or squamous cells (Figure 3o,p). Simulating the toggle-switch motif at varying myosin activation levels (Extended data figure 7o) predicted a maximal coefficient of variation per gastruloid for intermediate apical area, nicely recapitulating the experimental data across experimental conditions (Figure 3p; see Methods for details). Together, these findings support that double negative feedback between myosin and keratins produces two mechanical attractors: a low tensile/high-keratin, squamous shape in the amnion, while the epiblast remains high tension/high-myosin and columnar.

### YAP1 activity mediates the cytoskeletal switch in the amnion

Next, we examined the mechanisms mediating this cytoskeletal toggle switch, focusing on well-established mechanosensors able to regulate cell geometry and cytoskeletal organization. In line with recent studies^46,102–104^, we found that the mechanosensor YAP1 is enriched in the nucleus at the gastruloid edge (Figure 4a,b). This enrichment is specific to amnion cells, as unstimulated colonies showed lower and more edge-restricted YAP1 nuclear accumulation even after 48 h (Figure 4a,b). In line with this, we also found strong accumulation of nuclear YAP1 in small, unconfined colonies that uniformly differentiated into amnion (Extended data figure 8a,b, Figure 1e-g).

**Figure 4:**
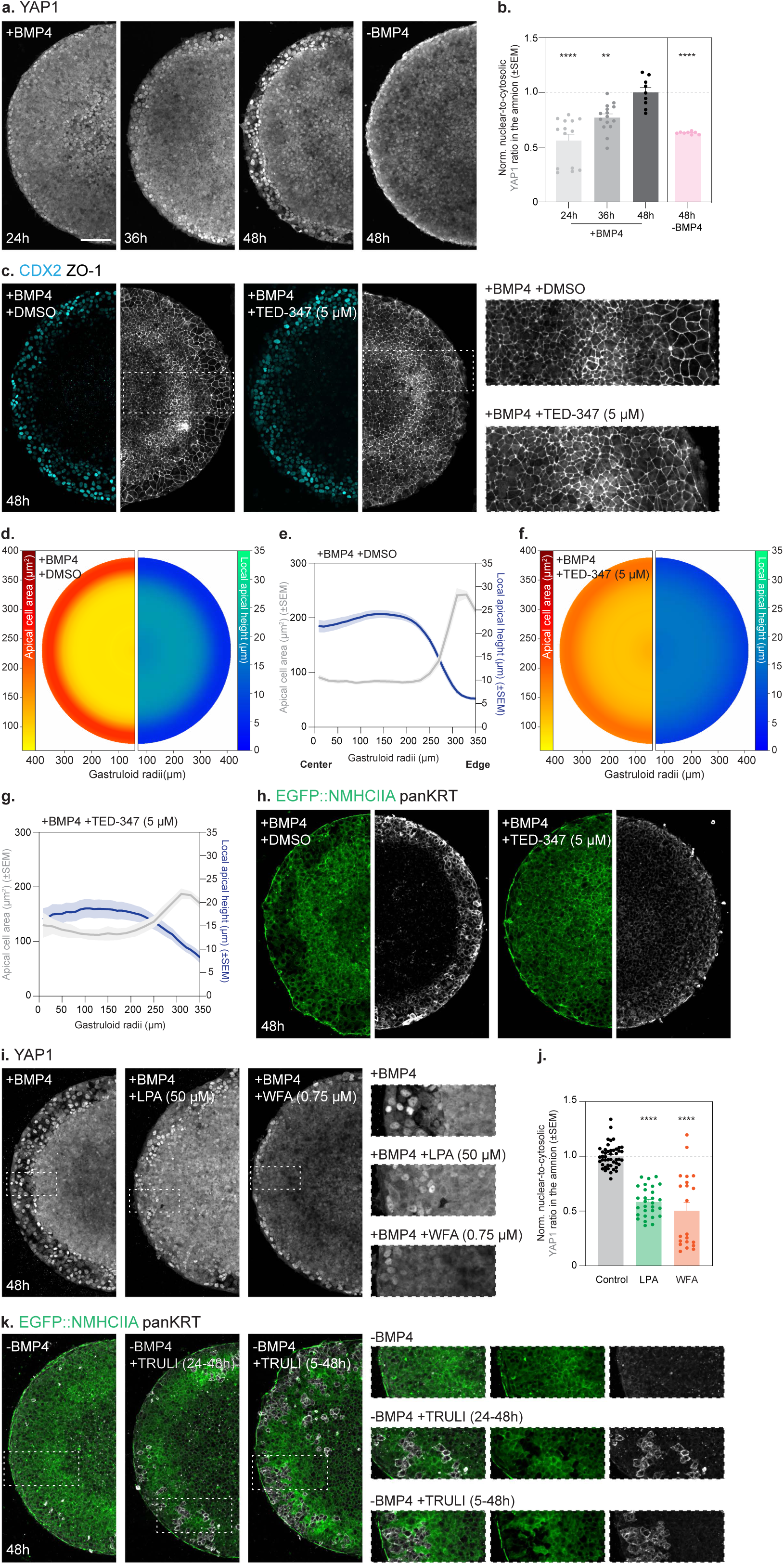
YAP1 mediates the cytoskeletal switch in the amnion. **a**. Immunostaining of YAP1 (grey) in BMP4-stimulated gastruloid discs at 24, 36 and 48h or unstimulated micropatterned colonies at 48h. Zoom-ins are shown on the right for 48h for discs that were either left untreated (-BMP4) or stimulated with BMP4 (+BMP4) for 48h. **b**. Normalized YAP1 nuclear-to-cytoplasmic ratio (see Methods) in the amnion domain after 24 (n=13, N=2), 36 (n=14, N=2) and 48h (n=9, N=2) of BMP4 stimulation or micropatterned colonies left unstimulated for 48h (n=10, N=1). **c**. Immunostaining of CDX2 (cyan) and ZO-1 (grey) in DMSO or TED-347-treated gastruloid discs after 48h of BMP4 stimulation. DMSO and TED-347 (5 µM) were added between 24 and 48h. Corresponding zoom-ins are shown on the right. (**d,f**) Average atlases of apical cell area (left) and local apical height (right) for DMSO (d, n=25, N=3) or TED-347-treated (f, n=19, N=3) gastruloid discs after 48h of BMP4 stimulation. (**e,g**) Average apical cell area (left axis) and apical height (right axis) as a function of gastruloid radii for DMSO (e) or TED-347-treated (g) gastruloid discs incubated with BMP4 ligands for 48h (n and N as in D,F). **h**. Immunostaining of pan-keratin (grey) in DMSO or TED-347-treated gastruloid discs expressing EGFP::NMHCIIA after 48h of BMP4 stimulation (see quantifications in Extended data figure 8e,f). **i**. Immunostaining of YAP1 (grey) in control, LPA (50 µM) or WFA (0.75 µM)-treated gastruloid discs after 48h of BMP4 stimulation. Zoom-ins are shown on the right. **j**. Normalized nuclear-to-cytoplasmic ratio of YAP1 (see Methods) in the amnion domain for control (n=46, N=4), LPA (50 µM; n=29, N=2) or WFA (n=21, N=2) treated gastruloid discs after 48h of BMP4 stimulation. **k.** Immunostaining of pan-keratin (grey) in micropatterned colonies that were either left untreated (-BMP4) or treated with TRULI (-BMP4 +TRULI) after 48h. TRULI (20 µM) was added between 24 and 48h or between 5 and 48h, as indicated in the figure. Zoom-ins are shown on the right. Maximum projection is shown for cell fate markers, pan-Keratin and YAP1, while the custom workflow was used for the apical surface projection of apical ZO-1 and EGFP::NMHCIIA. Scale bars: 100 μm (a,c,h,i,k). Error bars are shown as mean ± SEM (b,e,g,j). Kruskal-Wallis test. ****P<0.0001 (j).

To test whether YAP1 regulates amnion shape, we incubated gastruloid discs between 24 and 48 h with TED-347^105^, a potent inhibitor disrupting the interaction between YAP1 and its well-established partners, the TEAD transcription factors^106–108^. TED-347-treated gastruloid discs displayed both lower nuclear YAP1 at the gastruloid edge (Extended data figure 8c,d) and defective amnion flattening by 48 h (Figure 4c-g). In line with a role for YAP1 in regulating the cytoskeletal switch, the compromised amnion shape observed in TED-347-treated discs was accompanied by lower density of keratin filaments and higher actomyosin levels at the gastruloid edge (Figure 4h, Extended data figure 8e,f). Importantly, our findings indicate that YAP1 is not simply upstream of the cytoskeletal switch, since blocking amnion flattening, via LPA, WFA or KRT8/18 downregulation, also resulted in lower nuclear accumulation of YAP1 (Figure 4i,j, Extended data figure 8g,h). These experiments further support that, in the amnion, YAP1 nuclear accumulation responds to cell shape changes and not, as observed in other contexts^106–109^, to increased actomyosin tension, since actomyosin levels at the gastruloid edge in all these conditions were in fact higher than in control discs (Figure 3h-j, Extended data figure 4j,k, Extended data figure 6l-o). Overall, our findings support that YAP1 both senses BMP4-induced cell shape changes and feeds back onto them, by mediating double negative feedback between the myosin and keratin cytoskeleton (see Theory Note for incorporation of YAP1 within the toggle-switch model).

A key prediction of this model is that ectopic YAP1 activation is sufficient to trigger the cytoskeletal switch, even in the absence of BMP4 stimulation (see Theory Note for details). To functionally test this, we incubated micropatterned colonies with the LATS kinase inhibitor TRULI^110^ from 24 to 48 h, thus strongly enhancing nuclear accumulation of YAP1 (Extended data figure 8i,j). This ectopic nuclear accumulation of YAP1 was sufficient to trigger cytoskeletal rewiring, with co-existence of cells in the keratin-high (myosin-low) state and a large fraction of cells remaining myosin-high (keratin-low, Figure 4k). Extending TRULI exposure from 5 to 48 h further increased the proportion of cells undergoing the cytoskeletal switch (Figure 4k). Together, these findings identify a central role for YAP1 in mediating double negative feedback between the actomyosin and keratin networks and further support the proposed toggle switch logic, given the co-existence of two distinct cytoskeletal states within the same micropatterned colony (see Theory Note for details). Importantly, YAP1-dependent cytoskeletal rewiring was achieved without significant expression of amnion fate markers (Extended data figure 8k,l), supporting that the toggle switch arises at the level of the cytoskeleton and is not solely a consequence of fate changes.

### The cytoskeletal switch is necessary for amnion fate specification

Next, we asked whether the cytoskeletal switch, and thus amnion wetting, is necessary for amnion specification. Perturbing the cytoskeletal switch, either via LPA, WFA, downregulation of KRT8/18 or TED-347, resulted in a marked increase in SOX2 levels in the amnion domain (Figure 5a,b, Extended data figure 9a,b). This co-expression of amnion and pluripotency markers across orthogonal perturbations, suggested that shape changes in the amnion are needed for its fate specification. To determine how impairing the cytoskeletal switch alters the lineage trajectory of amnion cells, we next performed scRNA-seq of control (DMSO), LPA, WFA or TED-347-treated gastruloid discs at 48h of BMP4 stimulation (Figure 5c). Across perturbations, we found a consistent decrease in the expression of amnion markers, namely GATA3, HAND1, ISL1, GABRP, VTCN1 or WNT6 (Figure 5d, Extended data figure 9c). Since the expression of several of these genes was previously associated with amnion maturation^34,35,49^, these findings suggested that amnion wetting is required for its fate maturation. In line with this, we also found, particularly for WFA and TED-347-treated gastruloid discs, that a population of cells co-expressing amnion and epiblast markers was strongly enriched (Figure 5e, Extended data figure 9d-g). To further examine the differentiation trajectory of the amnion lineage in control (DMSO), LPA, WFA or TED-347-treated discs, we used Palantir^111^ and CellRank^112^ algorithms (Figure 5f, Extended data figure 9h). Perturbing the cytoskeletal switch consistently shifted the distribution of amnion cells towards earlier pseudotime (Figure 5g), coinciding with the expression kinetics of the early amnion marker GATA3 (Extended data figure 9i,j). Together, these findings support that amnion shape changes not merely follow cell fate specification but actively gate it.

**Figure 5:**
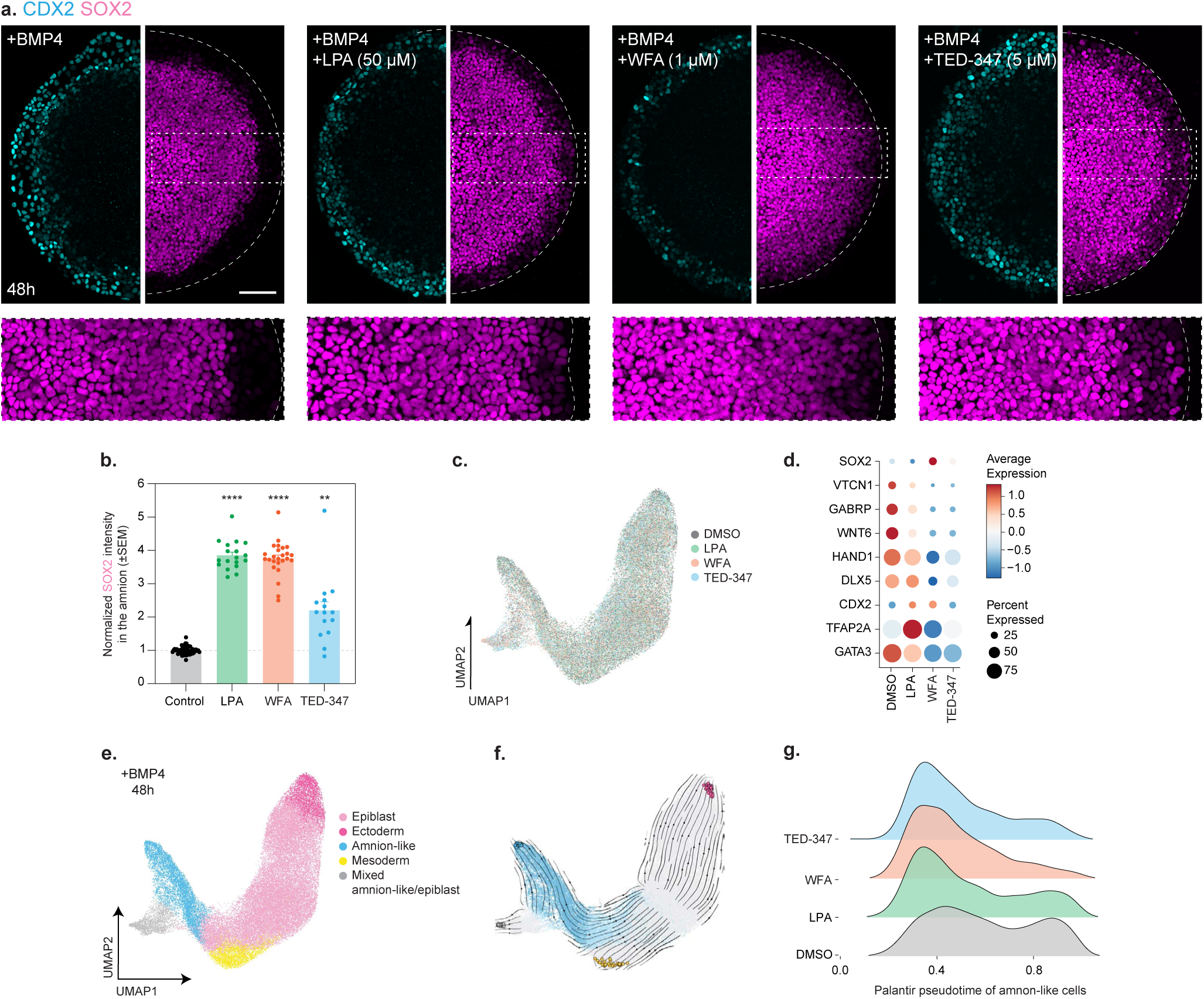
The cytoskeletal switch gates amnion specification. **a**. Immunostaining of control, LPA, WFA or TED-347-treated gastruloid discs, marked for CDX2 (cyan) or SOX2 (magenta) after 48h of BMP4 stimulation. Maximum projections are shown for cell fate markers, with the corresponding zoom-ins shown below. **b.** Normalized intensity of SOX2 for control (n=55, N=8), LPA (50 µM; n=18, N=2), WFA (n=25, N=3) or TED-347 (n=15, N=3)-treated gastruloid discs after 48h of BMP4 stimulation (see Methods for details). **c.** UMAP embedding of DMSO (n=11920 cells, N=2), LPA (n=12134 cells, N=2), WFA (n=11377 cells, N=2) and TED-347 (n=8946 cells, N=2)-treated gastruloid discs at 48h of BMP4 stimulation (n=44377 cells, N=2; see Methods for details), coloured by experimental condition. **d.** Dot plot showing selected marker genes in DMSO, LPA, WFA or TED-347-treated gastruloid discs at 48h of BMP4 stimulation for cells with an amnion-like signature, defined based on the gene signature enrichment scores (AUCell > 0.19, see Extended data figure 9c and Methods for details). Dot size shows the percentage of cells expressing the gene within the population and the colour intensity represents the mean expression. **e.** UMAP embedding of DMSO, LPA, WFA and TED-347-treated gastruloid discs at 48h of BMP4 stimulation, coloured by cell type derived via Leiden clustering and annotated by module scoring based on^39,40^ (n and N as in c, see Methods for details). **f.** CellRank-inferred cell fate dynamics. Streamlines show the CellRank transition matrix projected onto the UMAP embedding. Cells are coloured by their fate probability towards the amnion lineage, cells with a probability > 0.5 are highlighted in light blue and cells with a probability of 0.9 in darker blue. Remaining cells are shown in grey. Terminal states are colour-coded as in (e). **g.** Ridgeline plot of pseudotime distributions for cells committed to amnion fate (with a CellRank fate probability > 0.9) in gastruloid discs treated with DMSO, LPA, TED-347 or WFA at 48h of BMP4 stimulation (see Methods for details). LPA (50 µM) was added for 48h together with BMP4, while WFA (0.75 or 1 µM) and TED-347 (5 µM) were added only between 24 and 48h of BMP4 stimulation (a-g). Scale bars: 100 μm (a). (b) Kruskal-Wallis test. ****P<0.0001, **P=0.0019.

To understand how changes in cell shape are conveyed to the nucleus, we took advantage of a recently published single-nucleus RNA and ATAC sequencing dataset of self-organized gastruloid discs^41,42^ to infer enhancer-driven GRNs controlling amnion fate, using SCENIC+^113^ (Extended data figure 9k). Interestingly, this analysis once more converged on YAP1-TEAD signalling by showing that TEAD transcription factors are among the most active regulons in amnion-like cells (Extended data figure 9l, Supplementary Table 3). To validate these predictions experimentally, we turned back to our own scRNA-seq datasets and confirmed that the activity of both TEAD regulons is indeed strongly reduced in TED-347-treated discs (Extended data figure 9m). Importantly, perturbing the cytoskeletal switch directly, via LPA or WFA, had a similar effect (Extended data figure 9m), further linking amnion cell shape to YAP1-TEAD-dependent transcription. These findings place YAP1 as a central regulator of human gastrulation, not only in the embryonic^46,102,103,109^ but also in the extraembryonic compartment.

### Signatures for the cytoskeletal switch are conserved *in vivo*

To identify a transcriptional signature for the cytoskeletal switch, which could be used to probe the human amnion *in vivo*, we first confirmed, across scRNA-seq datasets of self-organizing gastruloid discs that both keratins and the inferred TEAD regulons are enriched in amnion-like cells (Figure 6a, Extended data figure 10a,b). This was accompanied by the upregulation of RhoGAPs, which are well-established negative regulators of actomyosin contractility^3,65,114^. Since RhoGEFs, positive regulators of contractility^3,65,114^, did not show the same enrichment (Figure 6a, Extended data figure 10a,b), these findings are consistent with our observations that the amnion is in a low tensile state. Importantly, the expression of keratins and RhoGAPs is also systematically reduced across orthogonal perturbations of the cytoskeletal switch (Figure 6b-d), thus experimentally validating its transcriptional signature. Remarkably, we found that this same transcriptional signature was conserved *in vivo*, with amnion cells in the CS7 human gastrula^115^ showing a strong upregulation of keratins, the inferred TEAD regulons, and RhoGAPs compared to epiblast cells (Figure 6e).

**Figure 6:**
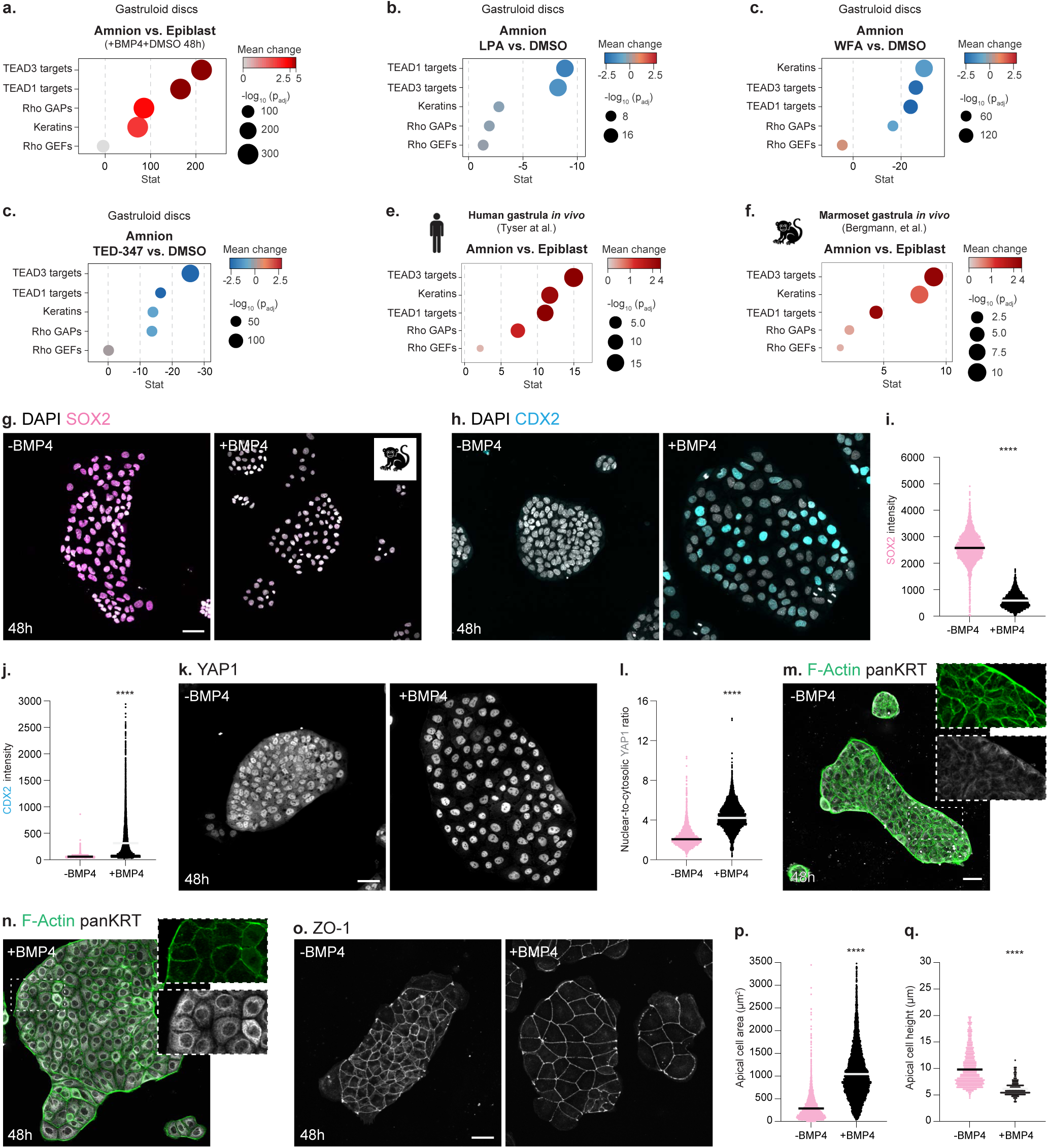
*In vivo* transcriptional signatures of the cytoskeletal switch in primate embryos, including humans. **a.** Difference in transcriptional activity for the cytoskeletal switch in amnion-like vs. epiblast cells in DMSO-treated gastruloid discs at 48h of BMP4 stimulation. Cell type classification is based on the gene signature enrichment scores (n=11920 cells, N=2, see Methods for details, see also Extended data figure 10a,b for the same analyses in other scRNA-seq datasets of gastruloid discs). (**b-d**) Difference in transcriptional activity for the cytoskeletal switch in LPA (n=12134 cells, N=2), WFA (n=11377 cells, N=2) and TED-347 (n=8946 cells, N=2)-treated gastruloid discs compared to control (DMSO, n=11920 cells, N=2) at 48h of BMP4 stimulation for cells with an amnion-like signature, defined based on the gene signature enrichment scores (AUCell > 0.19, see Extended data figure 9c and Methods for details). (**e,f)** Difference in transcriptional activity for the cytoskeletal switch in amnion vs. epiblast in the human (CS7^115^, e) or marmoset gastrula (CS6^31^, f) *in vivo*. (**g,h)** Immunostaining of SOX2 (g, magenta) or CDX2 (h, cyan) in unconfined marmoset colonies that were either left untreated (-BMP4) or stimulated with BMP4 ligands (+BMP4) for 48h. (**i,j)** Average intensity of SOX2 (i) or CDX2 (j) for unconfined marmoset colonies that were either left untreated (SOX2: n=3437 cells, N=3; CDX2: n=7804 cells, N=3) or stimulated with BMP4 ligands (SOX2: n=4842 cells, N=3; CDX2: n=9946, N=3) for 48h. **k**. Immunostaining of YAP1 (grey) in unconfined marmoset colonies that were either left untreated (-BMP4, left) or stimulated with BMP4 ligands (+BMP4, right) for 48h. **l**. Normalized nuclear-to-cytoplasmic ratio of YAP1 (see Methods), in unconfined marmoset colonies that were either left untreated (n=2164 cells, N=3) or stimulated with BMP4 ligands (n=6272 cells, N=3) for 48h. (**m,n**). Immunostaining of F-Actin (grey) and pan-keratin (grey) in unconfined marmoset colonies that were either left untreated (m, N=3) or stimulated with BMP4 ligands (n, N=3) for 48h. **o.** Immunostaining of ZO-1 (grey) in unconfined marmoset colonies that were either left untreated (-BMP4, left) or stimulated with BMP4 ligands (+BMP4, right) for 48h. (**p,q**) Average apical cell area (p) and apical-basal height (q) in unconfined marmoset colonies that were either left untreated (apical area: n=3738 cells, N=3; height: n=1040 cells, N=2) or stimulated with BMP4 ligands for 48h (apical area: n=8145 cells, N=3; height: n=399 cells, N=2). Maximum projections are shown for cell fate markers, ZO-1 and the cytoskeletal markers, with the corresponding zoom-ins shown below. Dot colour represents the mean change in expression, with dark red indicating higher activity, grey lower and blue negative correlation. Dot size indicates the significance of the difference, shown as -log10(adjusted p-value), with larger dots indicating greater significance (a-f). LPA (50 µM) was added for 48h together with BMP4, while WFA (1 µM) and TED-347 (5 µM) were added only between 24 and 48h of BMP4 stimulation (a,b). Scale bars: 50 μm (g,h,k,m,n,o). Mann-Whitney test. ****P<0.0001 (i,j,l,p,q).

Finally, since amnion fate programs and architecture are thought to be well conserved across primates^30,31,34^, we examined whether the cytoskeletal switch may operate in other species. Consistent with this, analysing a transcriptomic dataset for the CS6 marmoset gastrula^31^ revealed broad conservation of the YAP1-dependent cytoskeletal switch signature (Figure 6f). To test this transcriptional signature at protein level, we then stimulated small, unconfined colonies of marmoset PSCs with BMP4 *in vitro* (see Methods for details). After 48 h, these colonies were positive for typical amnion markers such as GATA3, CDX2 and HAND1 (Figure 6g-j, Extended data figure 10c,d). Strikingly, this was accompanied by increased nuclear accumulation of YAP1, a dense keratin network and lower F-Actin levels compared to colonies seeded in a similar fashion but left unstimulated for 48 h (Figure 6k-n). As expected from the wetting model, these BMP4-stimulated unconfined colonies displayed the characteristic squamous shape of amnion cells (Figure 6o-q, Extended data figure 10e). Together, these findings suggest that the cytoskeletal switch is a conserved feature of amnion formation in primates.

## Discussion

While recent studies shed light on the evolution and GRNs determining the identity of extraembryonic tissues during primate post-implantation development, their mechanical and morphogenetic programs have remained largely unknown^18,22,28,116^. In primates, including humans, amnion specification is accompanied by a drastic transition into a squamous, flattened architecture, giving the amniotic sac its distinctive asymmetric shape^19,21,22,26–31,47^. Yet, how the amnion achieves a sharp morphogenetic bifurcation from the adjacent epiblast, which remains columnar, was an open question. Here, we show that, in contrast to classical models in which shape change is driven by anisotropic tension along the apical-basal axis, amnion morphogenesis is achieved via a wetting transition, i.e. a switch into a low-tensile state. This contrasts with the prevailing view of extraembryonic tissues as passive mechanical support deformed by external forces^51–54^, and instead points to an active, cell-autonomous mode of amnion morphogenesis. Remarkably, modelling the wetting transition in a 3D geometry recapitulated the characteristic asymmetric shape of the amniotic sac *in vivo*^19,21,22,26–29,47^ and *in vitro* models of post-implantation amniotic sacs^84,85^.

Unlike wetting transitions based on tissue polarity or fluidization^117,118^, in the amnion, this transition is enforced through the rewiring of cytoskeletal composition, from actomyosin to keratin-based networks. In contrast to the prevailing view of keratins as passive mechanical resistance^86–90,92^, our study highlights that they can also drive autonomous deformation, via modulating actomyosin organization and cell mechanics. Beyond remodelling amnion architecture, keratin networks are also well placed to later allow the epithelium to withstand increasing foetal growth and fluid pressure within the amniotic cavity^18,22,28,35,116^, given their ability to strain-stiffen in response to applied force^86–89,91^.

Keratins were previously found to interact with actin filaments, for instance, during cell migration or early mouse development, where disrupting actin filaments compromises the stability of the keratin network and vice-versa^92,94,119–126^. Our findings reveal negative, rather than positive, feedback between these cytoskeletal networks — a mutual inhibition enforcing a sharp transition between two well-defined morphogenetic attractors: columnar versus squamous. This cytoskeletal switch is mediated by YAP1 but, unlike in mouse early development where YAP1 promotes keratin-dependent fate specification^94^, ectopic activation of YAP1 is not sufficient to induce amnion fate. This supports that the keratin-actomyosin toggle switch operates at the level of the cytoskeleton and is not merely a consequence of fate changes. The cytoskeleton thus implements the switch-like logic generally assigned to GRNs^99–101^, providing a design principle enabling sharp morphogenetic bifurcations.

The link between fate specification and morphogenetic programs has become increasingly appreciated over recent years, with recent studies uncovering, for instance, a role for membrane tension in gating the transition from naive-to-primed pluripotency^16,127,128^. Our study extends this notion by showing that shape changes gate amnion fate maturation. Together, our data support a model in which BMP4 stimulation initiates amnion specification and shape changes, which in turn promote YAP1 nuclear accumulation and binding to TEAD factors. YAP1-TEAD signalling further reinforces both the cytoskeletal switch and amnion specification, yielding a mature, fully flattened amniotic epithelium. Our findings provide a mechanistic explanation for the recently reported role of substrate stiffness^93^ or cell density in amnion specification^49^ and explain recent observations in the primate gastrula *in vivo*, where the columnar-to-squamous transition within the amniotic sac was shown to coincide with different states of amnion maturation. Indeed, cells at the boundary with the columnar epiblast, which displayed only intermediate flattening, were found to co-express both amnion and pluripotency markers, including SOX2, while fully squamous cells expressed solely markers for the amnion lineage^34^. Since amnion-secreted signals were shown to play a key role in mesoderm specification in primates^36,129^, coupling its sharp morphogenetic transition to fate programs likely ensures not only robust assembly of this key extraembryonic tissue but also supports gastrulation.

Finally, we show that the cytoskeletal switch was recapitulated in non-human primate stem cell models *in vitro*, with marmoset-derived amnion cells adopting the same keratin-high (actomyosin-low) state identified in human gastrula models. Consistent with the idea that this mechanical program is conserved across primates, the transcriptional signatures of the cytoskeletal toggle switch identified and validated *in vitro* were recapitulated *in vivo,* in both human and non-human primate amnion. Overall, our findings support that the cytoskeleton can act not only as an effector of tissue shape but as a decision-making module, encoding the same mutual-inhibition logic used by GRNs to drive cells into discrete architectures. Because keratins are ubiquitously expressed not only across several extraembryonic tissues^35,39,93,95^, but also throughout mammalian epithelia in development, homeostasis and disease^130–133^, such cytoskeletal toggle switches may represent a general strategy for generating divergent epithelial architectures.

## Supporting information

Roffay_et-al_Theory-Note

## Author contributions

C.R. and D.P. designed the research. C.R. and S.J. performed the experiments and analysed the experimental data. O.D. developed the theoretical models and simulations. O.D. performed the 3D reconstructions of self-organizing gastruloid discs and contributed to the analysis of the experimental data. C.H. and C.R. established fluorescent reporters. M.N. performed the bioinformatic analyses. E.J. and A.M.C. provided key expertise and supported C.R. respectively with tether pulling assays or FLIM imaging. N.R. provided key reagents and supervised C.R. with marmoset PSC experiments. D.P. and É.H. wrote the manuscript with input from all authors.

## Acknowledgements

We thank Prisca Liberali, Anna Kicheva, Yohanns Bellaïche, Andrea Pauli, Alexandra Schauer and Guilherme Ventura for feedback on the manuscript. We thank the members of the Rivron, Mendjan, Tanaka and Knoblich labs, in particular Heidar Heidari Khoei, Theresa Sommer, Giovanni Sestini, Jana Slovakova, Yvonne Scholte Op Reimer, Stefan Jahnel, Hannah Stuart, Jamie Littleboy and Sandra Schepers for technical advice and/or reagents. We thank all members of the Pinheiro and Hannezo groups for discussions and technical advice. We thank all members of the BiOptics and the Tissue Culture Support Unit at IMP/IMBA/GMI for continuous support. We thank the Bioimaging facility at ISTA, specifically Gabriel Krens, for support with tether pulling assays. This project has received funding from the European Union’s Framework Programme for Research and Innovation Horizon 2020 (2014-2020) under the Marie Skłodowska-Curie Grant Agreement Nr. 847548 to C.R. and the European Union (European Research Council starting grant 101116586 to DP). The IMP receives institutional funding from Boehringer Ingelheim and the Austrian Life Sciences Program 2023 (# 48924910).

## Declaration of Interests

The authors declare no competing interests.

## Extended data figure legends

**Extended data figure 1:**
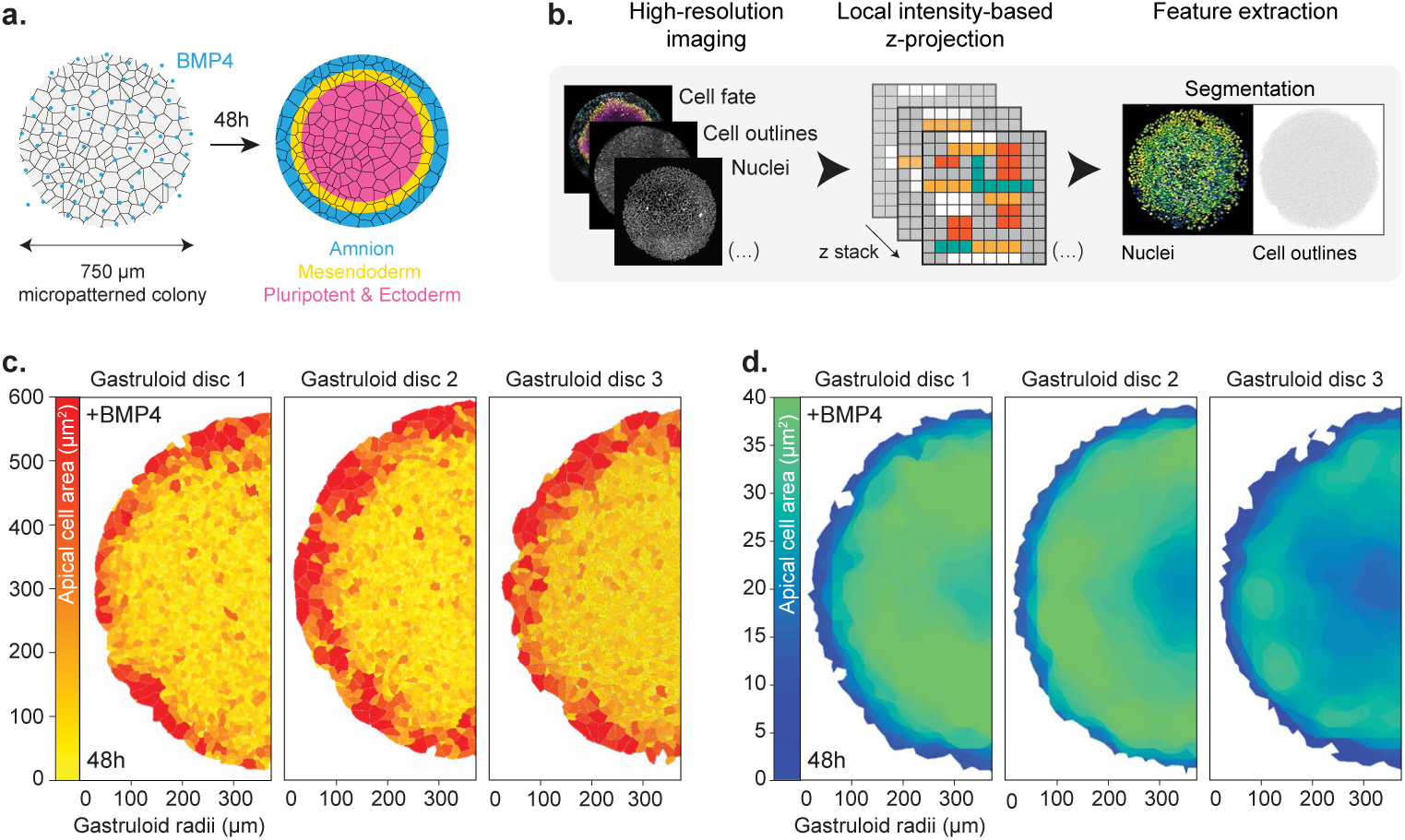
Scalable morphometric pipeline to extract cell shape descriptors during gastruloid disc self-organization. **a**. Schematic representation of gastruloid disc self-organization. **b**. Schematic representation of the custom workflow established to quantify the morphological signatures of gastruloid discs during self-organization. (**c,d**). Examples of apical cell area (c) or local apical height (d) maps for individual self-organized gastruloid discs after 48h of BMP4 stimulation.

**Extended data figure 2:**
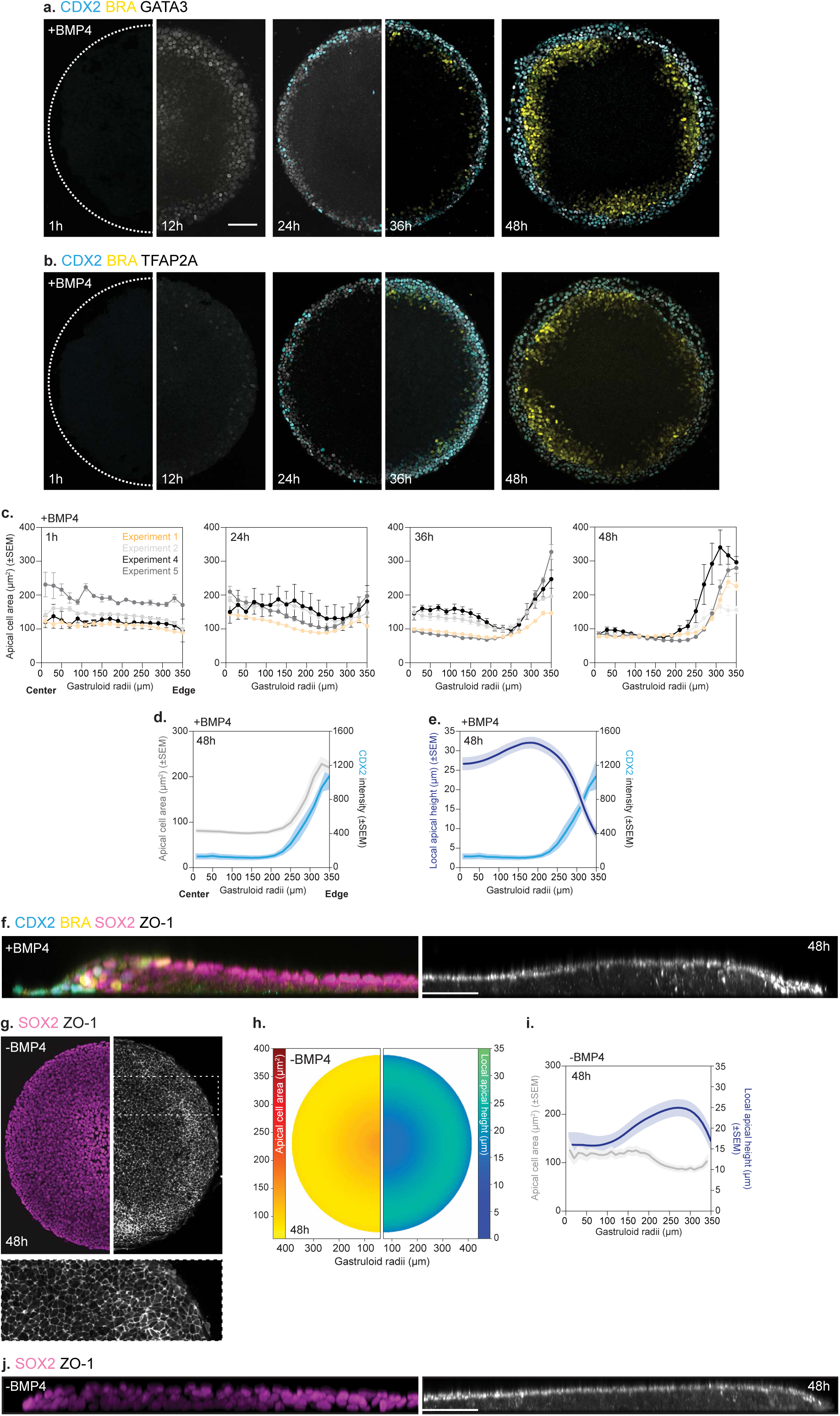
Distinct morphological signatures arise due to gastruloid disc self-organization and not geometrical confinement. (**a,b**) Immunostaining of gastruloid discs, marked for CDX2 (cyan), BRA (yellow) and GATA3 (a, amnion: grey) or TFAP2A (b, amnion: grey) after 1, 12, 24, 36 and 48h of BMP4 stimulation. **c**. Average apical cell area as a function of gastruloid radii for individual experiments after 1, 24, 36 and 48h of BMP4 stimulation (n and N as indicated in Figure 1c). (**d,e**) Average apical cell area (d, left axis) or local apical height (e, left axis) and CDX2 intensity (d,e, right axis) as a function of gastruloid radii after 48h of BMP4 stimulation (n and N as Figure 1b,c). **f**. Side view of immunostained gastruloid discs marked for CDX2 (cyan), BRA (yellow), SOX2 (magenta) and ZO-1 (grey) for BMP4-stimulated (+BMP4) gastruloid discs after 48h. **g**. Immunostaining of micropatterned colonies, marked for SOX2 (magenta) and ZO-1 (grey), that were left untreated for 48 h. Zoom-in images are shown below. **h**. Average atlases of apical cell area (left) and local apical height (right) in micropatterned colonies that were left untreated for 48 h (n=18, N=3). **i**. Average apical cell area (left axis) and local apical height (right axis) as a function of the radii in micropatterned colonies that were left untreated for 48h (n and N as in h). **j.** Side view of micropatterned colonies immunostained for SOX2 (magenta) and ZO-1 (grey) (-BMP4) that were left untreated for 48h. Maximum projection is shown for cell fate markers, and the custom workflow was used for the apical surface projection, based on ZO-1 local intensity (see Methods for details). Scale bars: 100 μm (a,b,f,g,j). Error bars are shown as mean ± SEM (c-e,i).

**Extended data figure 3:**
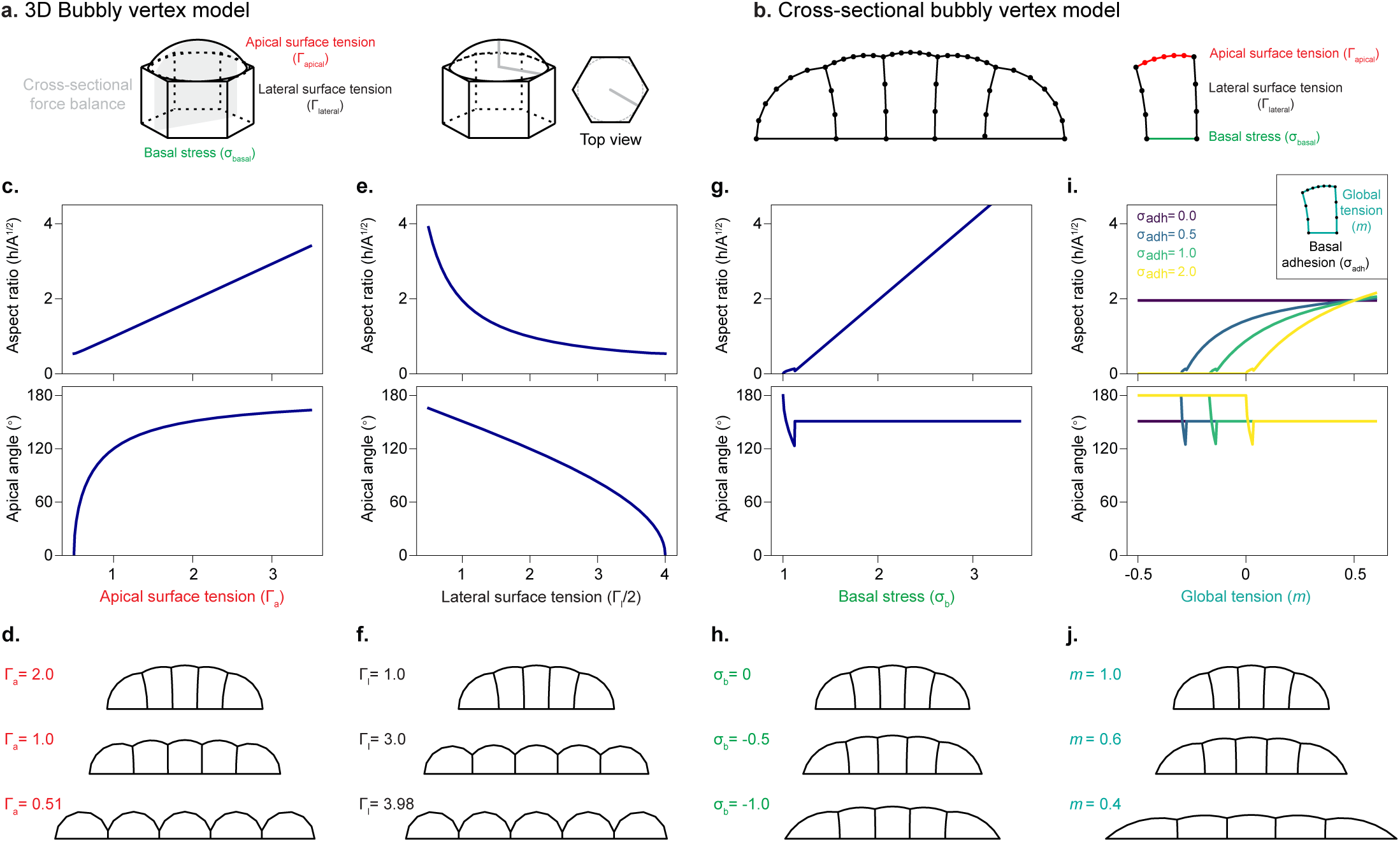
Testing distinct strategies for cell flattening in the bubbly vertex model. **a**. Schematic representation of the analytical approximation for the 3D bubbly vertex model (BVM), in which the cell is modelled as a hexagonal prism with a spherical cap. The cap radius is assumed to be the incircle radius of the regular hexagon. We assume apical, lateral and basal tensions, as well as basal adhesion (which rescales basal tension, defining effective basal stress). Lateral interfaces are shared with neighbouring cells. Apical-lateral contact angles are determined based on the ratio of apical and lateral tensions. **b.** Schematic depiction of the numerical cross-sectional vertex model. Each cell is discretized via nodes and straight edges connecting these nodes, where distinct apical, lateral and basal line tensions are implemented, with basal line tensions encoding the basal stress, as in (A). (**c-j**) Aspect ratio (top) and interfacial apical angles (middle) for different flattening strategies employed in the 3D BVM (c,e,g,i). Corresponding images of cross-sectional 2D BVM simulations of five cells undergoing flattening, via each of these strategies are shown below (bottom; d,f,h,j). The strategies used are: i) an apical tension decrease (c,d), ii) a lateral tension increase (e,f); iii) a basal stress decrease (g,h) and, finally, iv) a global tension decrease, in the presence of basal adhesion (i,j). The modified parameter values in the simulations (d,f,h,j) are indicated next to each simulation.

**Extended data figure 4:**
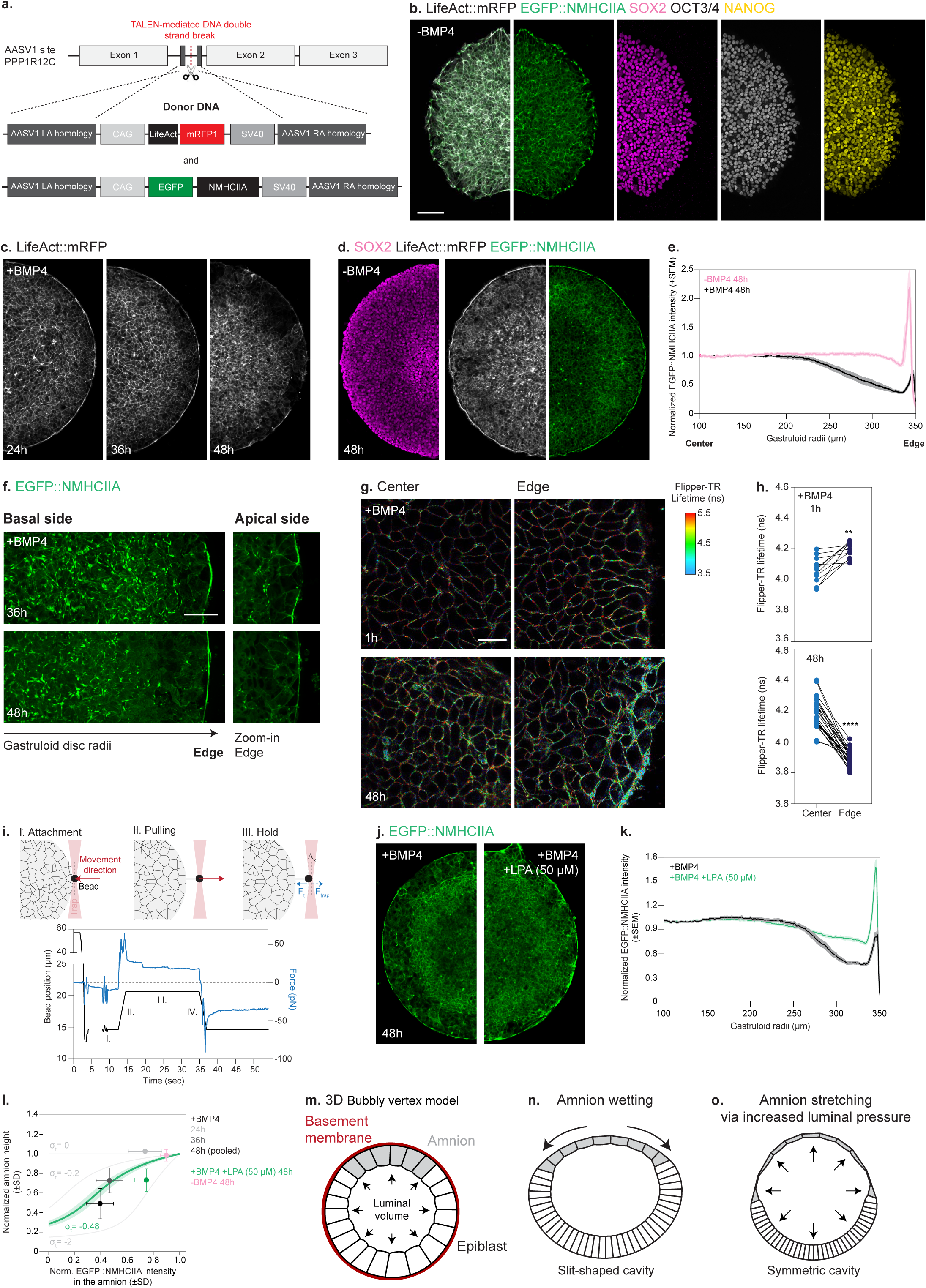
Direct tension measurements in self-organizing gastruloid discs. **a**. Schematic representation of the editing approach used to generate the dual H9 reporter line for LifeAct::mRFP1 and EGFP::NMHCIIA. **b**. Untreated micropatterned colonies, expressing LifeAct::mRFP1 (grey) and EGFP::NMHCIIA (green), immunostained for several pluripotency markers, namely SOX2 (magenta), NANOG (yellow) and OCT3/4 (grey; n=21, N=2). **c**. LifeAct::mRFP (gray)-expressing gastruloid discs after 24, 36 and 48 h of BMP4 stimulation. Corresponding EGFP::NMHCIIA expression and fate markers for these gastruloid discs are shown in Figure 2f (n and N as indicated in Figure 2G). **d**. Immunostaining of SOX2 (magenta) in micropatterned colonies expressing EGFP::NMHCIIA (green) that were left untreated for 48h. **e**. Normalized EGFP::NMHCIIA intensity as a function of radii in BMP4-stimulated gastruloid discs (+BMP4, n=24, N=2) or micropatterned colonies (-BMP4, n=7, N=1) after 48h. **f**. Snapshot of the basal (left) and apical sides (right) of amnion cells in gastruloid discs expressing EGFP::NMHCIIA (green) after 36 or 48h of BMP4 stimulation (n and N as in Figure 2G). (**g**) Fluorescence lifetime imaging of the Flipper-TR probe, a proxy for membrane tension, during gastruloid disc self-organization (1h and 48h after BMP4 stimulation). FLIM images are colour-coded by Flipper-TR lifetime (ns) and the corresponding scale is shown below. FLIM images were acquired in regions of interest located at the edge or in the centre of self-organizing gastruloid discs, as indicated in the figure. **h.** Average Flipper-TR lifetime in the edge versus centre of the gastruloid disc after 1 (n=13, N=2) and 48h (n=29, N=3) of BMP4 stimulation. **i**. Representative curve indicating both the position of the trap (black, left) and the force measured on the trap (blue, right) during a tether pulling assay. A schematic representation of the assay is shown on the top and directly indicated in the curve. I. Bead attachment. II. Tether pulling. III. Hold the bead in position. IV. Bead retraction (not shown as a schematic). **j**. EGFP::NMHCIIA (green)-expressing gastruloid discs treated for 48h with BMP4 ligands alone or together with LPA (50 µM). **k**. Normalized intensity of EGFP::NMHCIIA as a function of gastruloid radii for control (n=24, N=2) or LPA-treated discs (50 µM; n=29, N=2) for 48h. **l.** Predictions from the 3D bubbly vertex model (see Theory Note for details) for the height of amnion cells as a function of the fold-change in EGFP::NMHCIIA intensity versus the experimentally determined values for control self-organizing gastruloid discs or treated with LPA (50 µM) and unstimulated micropatterned colonies at 48h (n and N as in (k)). Green line corresponds to maximum-a-posteriori Bayesian fit of confinement stress, with the associated confidence interval. **m.** 3D cross-sectional model of the amniotic sac, composed of amnion and epiblast cells surrounded by a thin ECM layer to mimic the early human gastrula (see Theory Note for details). **n**. Modelling the squamous transition of amnion cells via active wetting. This recapitulates both the characteristic low curvature of the amnion domain and the slit-shape organization of the amniotic cavity (see Theory Note for details). **o**. Modelling the squamous transition of amnion cells via increased hydrostatic pressure within the amniotic cavity. Since amnion cells were found to adopt a low tensile state (Figure 2l, Extended data figure 4f-i), this increased hydrostatic pressure is expected to produce effective amnion stretching. However, it results in a fully spherical shape for the amniotic cavity (see Theory Note for details). Maximum projection is shown for cell fate markers, and the custom workflow was used to project the apical surface, based on EGFP::NMHCIIA intensity. Scale bars: 100 μm (b,c,j), 30 μm (f,g) and 50 μm (d). Error bars are shown as mean ± SEM (d,k) and as mean ± SD (l). Paired t test (for 1h). **P=0.0022. Wilcoxon test (for 48 h). ****P<0.0001 (h).

**Extended data figure 5:**
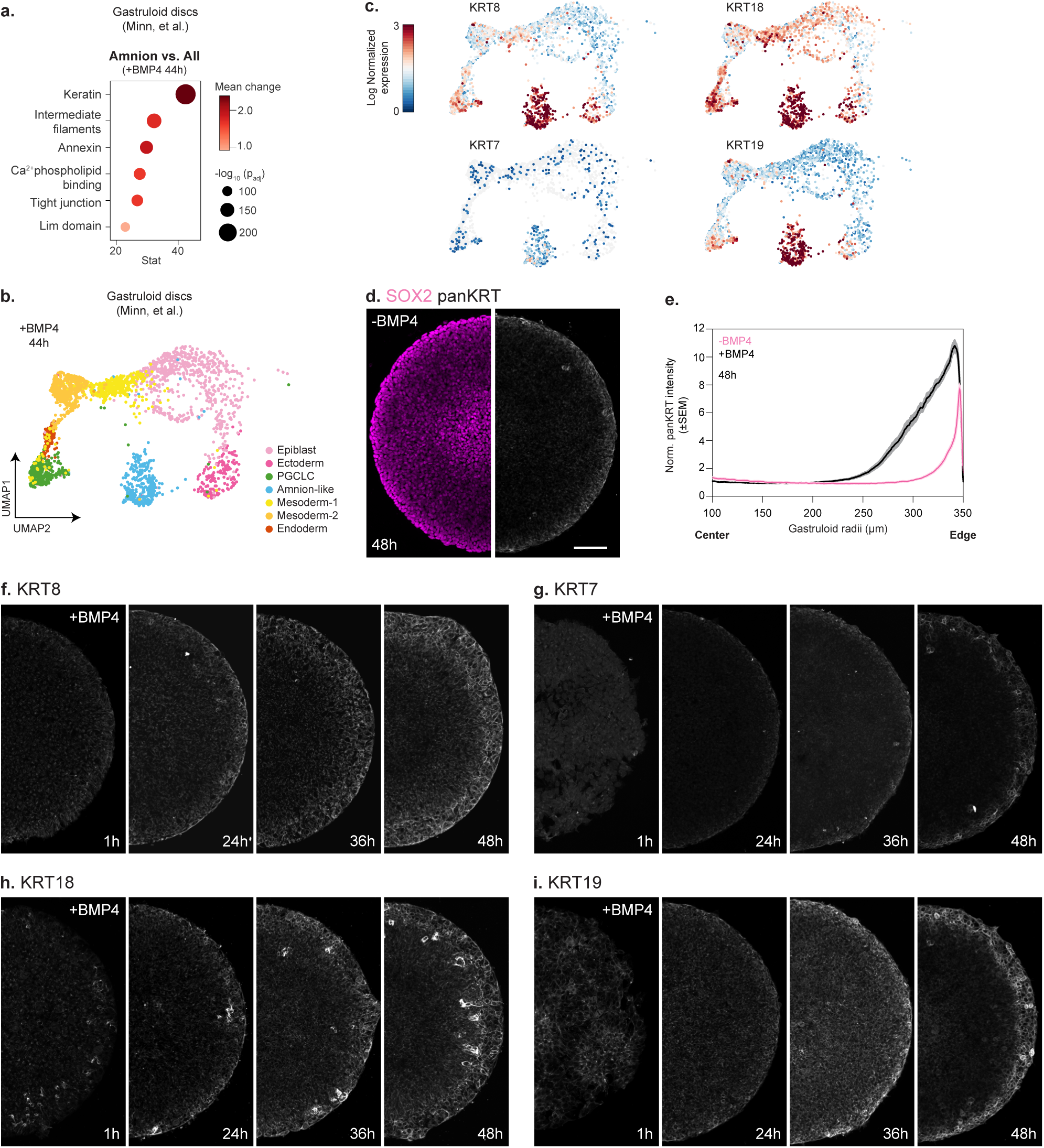
Multiple keratin isoforms are dynamically enriched in the amnion. **a.** Enrichment analysis of UNIPROT categories in amnion-like cells compared to all other annotated cell types in gastruloid discs upon 44h of BMP4 stimulation (dataset and annotations from^39,40^, see Methods for details). **b.** UMAP displays of cells after 44h of BMP4 stimulation clustered by cell type. PGCLC: primordial germ cell-like cells. **c.** Normalized expression of the indicated markers after 44h of BMP4 stimulation on the UMAP clustered by cell type. **d**. Immunostaining of micropatterned colonies, marked for SOX2 (pluripotent: magenta) and pan-Keratin (panKRT, grey), that were left untreated for 48h. **e**. Normalized intensity of pan-Keratin as a function of radii for gastruloid discs stimulated with BMP4 ligands (n=24, N=3) or left untreated (n=10, N=1) for 48h. (**f-i**) Immunostaining of BMP4-stimulated gastruloid discs, marked for Keratin 8 (f, KRT8, grey; 1h: n=4; 24h: n=4; 36h: n=5; 48h: n=5, N=1 for all timepoints), Keratin 7 (g, KRT7, grey; 1h: n=8; 24h: n=9; 36h: n=10; 48h: n=10, N=2 for all timepoints), Keratin 18 (h, KRT18, grey; 1h: n=5; 24h: n=5; 36h: n=6; 48h: n=5, N=1 for all timepoints) or Keratin 19 (i, KRT19, grey; 1h: n=9; 24h: n=10; 36h: n=13; 48h: n=11, N=2 for all timepoints) after 1, 24, 36 and 48h. Maximum projection is shown for cell fate markers and pan-Keratin. Scale bars: 100 μm (d,f-i). Error bars are shown as mean ± SEM (e).

**Extended data figure 6:**
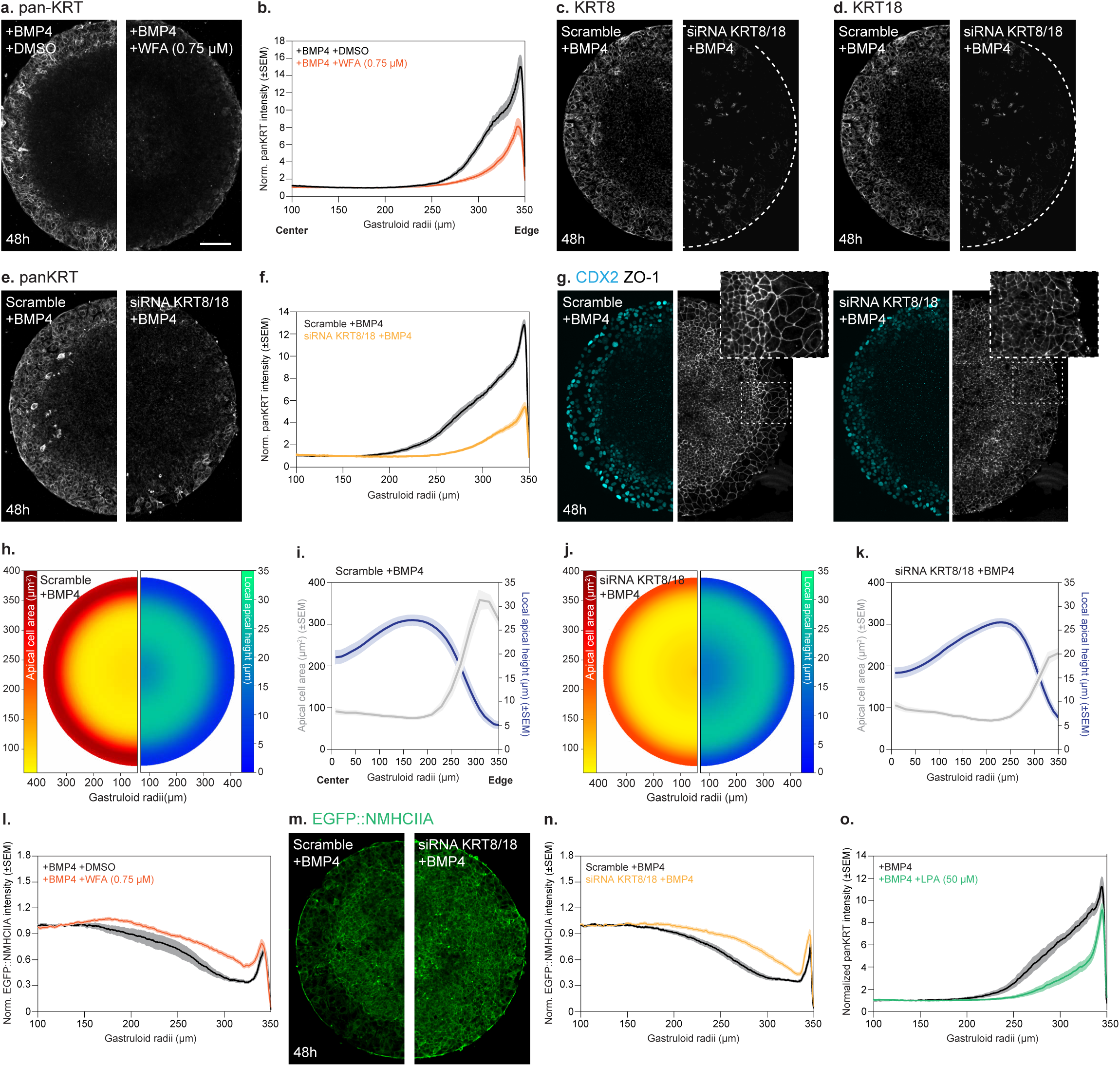
Rewiring cytoskeletal composition, via dual negative feedback between actomyosin and keratins, is necessary for amnion flattening. **a.** Immunostaining of DMSO or WFA-treated gastruloid discs, marked for pan-Keratin (grey), after 48h of BMP4 stimulation. DMSO and WFA were added between 24 and 48h of BMP4 stimulation. **b.** Normalized intensity of pan-Keratin as a function of gastruloid disc radii for DMSO (n=12, N=2) or WFA (n=17, N=2)-treated gastruloid discs after 48 h of BMP4 stimulation. (**c,d**) Immunostaining of gastruloid discs treated with a mix of non-targeting control (scramble) or KRT8/18 (right) siRNAs, marked for Keratin 8 (c, grey, n=24; N=2) or Keratin 18 (d, grey, n=24; N=2) after 48 h of BMP4 stimulation. **e**. Immunostaining of gastruloid discs treated with a mix of non-targeting control (scramble) or KRT8/18 siRNAs, marked for pan-Keratin (grey), after 48 h BMP4 stimulation. **f**. Normalized intensity of pan-Keratin as a function of radii in non-targeting control (scramble; n=31, N=2) or KRT8/18 (n=37, N=2) siRNAs treated gastruloid discs after 48 h of BMP4 stimulation. **g.** Immunostaining for CDX2 (cyan) and ZO-1 (grey) in gastruloid discs treated with non-targeting control (scramble) or KRT8/18 (right) siRNAs (see Methods for details). Corresponding zoom-ins are shown on the top right. (**h,j**) Average atlases of apical cell area (left) or local apical height (right) in gastruloid discs treated with a mix of non-targeting control (scramble, h, n=24; N=2) or KRT8/18 (j, n=25, N=2) siRNAs and stimulated with BMP4 ligands for 48h. (**i,k**) Average apical cell area (left axis) and local apical height (right axis) as a function of radii for gastruloid discs treated with a mix of non-targeting control (scramble, i) or KRT8/18 (k) siRNAs and stimulated with BMP4 for 48 h (n and N as in H,J). **l.** Normalized intensity of EGFP::NMHCIIA as a function of radii for DMSO (n=13, N=2) or WFA (n=20, N=2) treated gastruloid discs and stimulated with BMP4 for 48h. **m.** EGFP::NMHCIIA (green)-expressing gastruloid discs treated with a mix of non-targeting control (scramble) or KRT8/18 siRNAs after 48h of BMP4 stimulation. **n.** Normalized intensity of EGFP::NMHCIIA as a function of radii for gastruloids treated with a mix of non-targeting control (scramble, n=36, N=2) or KRT8/18 (n=36, N=2) siRNAs after 48h of BMP4 stimulation. **o**. Normalized intensity of pan-Keratin as a function of gastruloid radii after 48 h of incubation with BMP4 ligands alone (n=21, N=2) or together with LPA (50 µM, n=22, N=2). Maximum projection is shown for cell fate markers and pan-Keratin, while the custom workflow was used for projecting the apical surface, based on the intensity of EGFP::NMHCIIA and ZO-1. Scale bars: 100 μm (a,c-e,g,m). Error bars are shown as mean ± SEM (b,f,i,k,l,n,o).

**Extended data figure 7:**
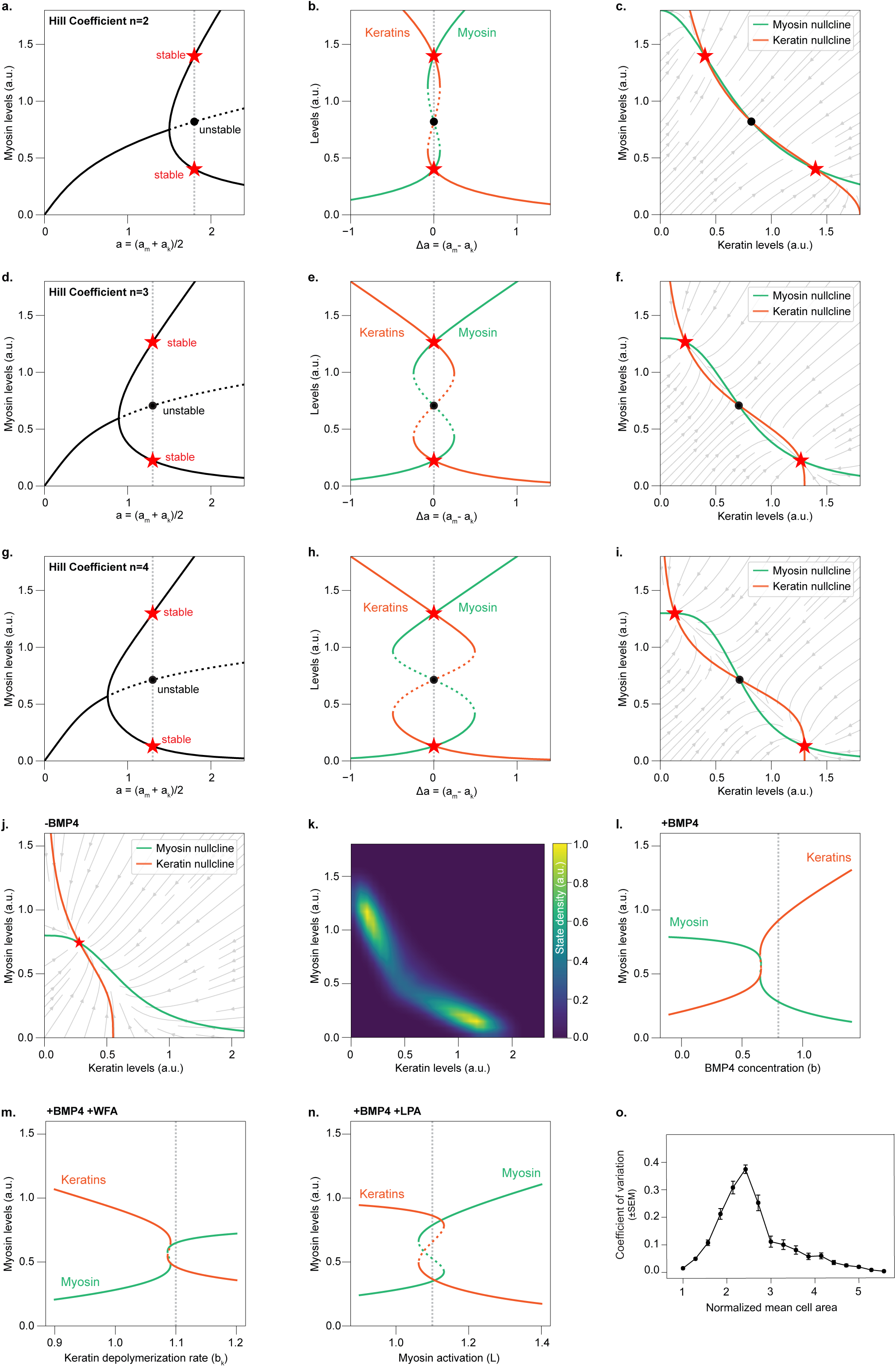
Theoretical modelling of a cytoskeletal toggle switch motif. (**a-c**) Solutions of the toggle switch model for Hill coefficient n=2. **a.** Stable (full black lines) and unstable (dotted black line) steady states of the toggle switch equations, characterized by a pitchfork bifurcation upon increase of negative feedback coupling (assuming symmetric couplings between relative myosin and keratin intensities). Stable/unstable states for chosen parameter values are indicated as respectively the red or black markers, with a vertical dotted line indicating the used parameters. **b.** Asymmetric coupling strengths lead to a loss of bistability. **c.** Phase portrait of the toggle switch for the chosen parameters (marked in a) with flow field indicating the trajectories in myosin-keratin phase space, and nullclines from the myosin and keratin conservation equation (respectively shown in green and orange). Red markers indicate stable states from (a), located at intersections of the nullclines. (**d-i**) Toggle switch model with Hill coefficients n=3 (d-f) or n=4 (g-i) showing qualitatively the same behaviour as in (a-c). **j.** Phase portrait of the best-fit parameters without BMP4 stimulation with only a high myosin and low keratin steady state (red marker). **k.** Sampled landscape, shown as state density, with the two attractors of the toggle switch motif. (**l-n**) Bifurcation diagrams showing myosin and keratin intensities of the toggle switch with best-fit parameters upon BMP4 stimulation alone (l), or together with WFA (m) or LPA (n). The pharmacological perturbations respectively disrupt keratin filament polymerization (modelled as increase in keratin degradation rate) or promote actomyosin contractility (modelled via an increase of the myosin activation rate). The vertical dotted line indicates the best-fit value (cf. Figure 3l-n). **o.** Simulated variability in amnion cell area, shown as coefficient of variation, as a function of the mean apical cell area in the amnion. Error bars are shown as mean ± SEM and were generated from 1000 simulations with varying levels of myosin activation (see Theory Note). In (a,b,d,e,g,h,j,l,m,n), the solid lines represent stable steady states, and dotted lines represent unstable states.

**Extended data figure 8:**
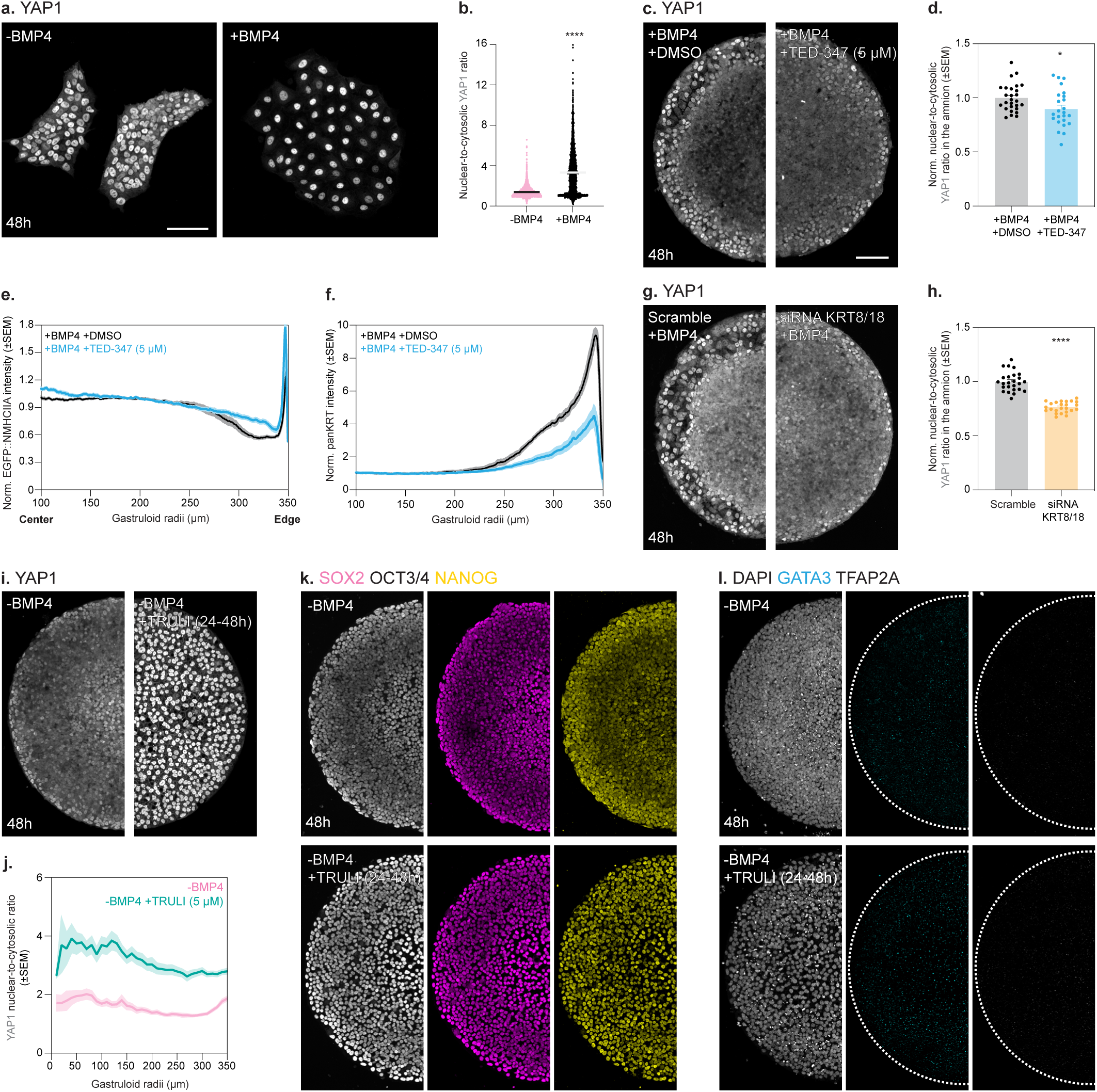
Activating YAP1 is sufficient to induce cytoskeletal rewiring in the absence of BMP4 or amnion induction. **a.** Immunostaining of YAP1 (grey) in unconfined colonies that were either left untreated (-BMP4) or stimulated with BMP4 ligands (+BMP4) for 48h. **b.** YAP1 nuclear-to-cytosolic ratio for unconfined colonies that were either left untreated (n=3166 cells, N=2) or stimulated with BMP4 ligands (n=2419 cells, N=2) for 48h. **c.** Immunostaining of YAP1 (grey) in DMSO or TED-347-treated gastruloid discs after 48h of BMP4 stimulation. DMSO or TED-347 (5 µM) were added between 24 and 48h of BMP4 stimulation. **d.** Normalized nuclear-to-cytoplasmic ratio of YAP1 (see Methods) in the amnion domain for DMSO (n=27, N=3) or TED-347 (n=24, N=3)-treated gastruloid discs after 48h of BMP4 stimulation. (**e,f**) Normalized intensity of EGFP::NMHCIIA (e) or pan-Keratin (f) as a function of radii for gastruloid discs treated with DMSO (e: n=22, N=2, f: n=14, N=3) or TED-347 (e: n=18, N=2, f: n=14, N=3) after 48h of BMP4 stimulation. **g.** Immunostaining of YAP1 (grey) in gastruloid discs treated with a mix of non-targeting control (scramble) or KRT8/18 siRNAs after 48h of BMP4 stimulation. **h.** Normalized nuclear-to-cytoplasmic ratio of YAP1 (see Methods) in the amnion domain for gastruloid discs treated with a mix of non-targeting control (scramble, n=25, N=2) or KRT8/18 (n=24, N=2) siRNAs after 48h of BMP4 stimulation. **i.** Immunostaining of YAP1 (grey) in micropatterned colonies left untreated or incubated with TRULI (20 µM) at 48h. **j.** YAP1 nuclear-to-cytoplasmic ratio in micropatterned colonies left untreated (n=23, N=4) or incubated with TRULI (20 µM; n=28, N=4) at 48h (see Methods for details). **k.** Immunostaining of SOX2 (magenta), OCT3/4 (grey) and NANOG (yellow) in micropatterned colonies left untreated (SOX2: n=22, N=4; OCT3/4: n=12, N=2; NANOG: n=12, N=2) or incubated with TRULI (SOX2: n=24, N=4; OCT3/4: n=12, N=2; NANOG: n=12, N=2) at 48h. **l.** Immunostaining of DAPI (grey), GATA3 (cyan) and TFAP2A (grey) in micropatterned colonies left untreated (GATA3: n=26, N=4; TFAP2A: n=24, N=4) or incubated with TRULI (GATA3: n=26, N=4; TFAP2A: n=26, N=4) at 48h. TRULI (20 µM) was added between 24 and 48h in (i-k). Scale bars: 100 μm (a,c,g,i-l). Maximum projections are shown for YAP1 and cell fate markers. Error bars are shown as mean ± SEM (b,d-f,h,j). t test. *P=0.0187. (d). Welch’s *t* test. ****P<0.0001 (h).

**Extended data figure 9:**
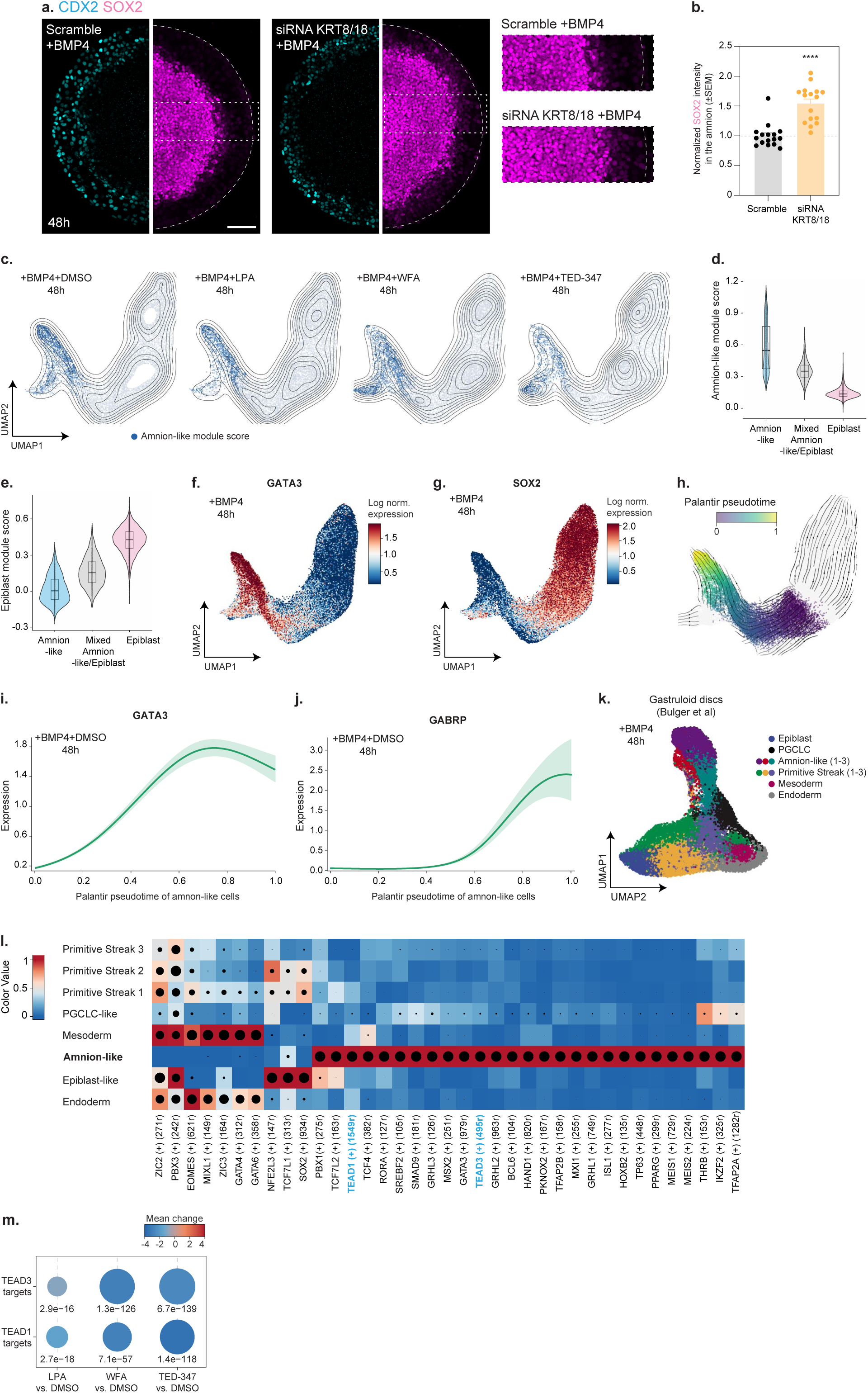
TEAD transcription factors, well-established co-transcriptional partners of YAP1, are highly enriched in amnion cells. **a.** Immunostaining of gastruloid discs treated with non-targeting control (scramble) or KRT8/18 siRNAs, marked for CDX2 (cyan) and SOX2 (magenta) after 48h of BMP4 stimulation. (**b**) Normalized intensity of SOX2 (b) for gastruloid discs treated with non-targeting control (scramble, n=16, N=2) or KRT8/18 (n=16, N=2) siRNAs after 48h of BMP4 stimulation (see Methods for details). Maximum projection is shown for cell fate markers, and the corresponding zoom-ins are shown below. (**c**) UMAP embedding of DMSO (n=11920 cells, N=2), LPA (n=12134 cells, N=2), WFA (n=11377 cells, N=2) and TED-347 (n=8946 cells, N=2)-treated gastruloid discs at 48 h of BMP4 stimulation, coloured for cells with an amnion-like signature, defined based on the gene signature enrichment scores (AUCell > 0.19, see Methods for details). (**d,e**) Amnion-like (d) or epiblast (e) module score enrichment in amnion-like, mixed amnion-like/epiblast and epiblast cells, defined based on the gene signature enrichment scores (see Figure 5e and Methods for details). (**f,g**) UMAP embedding of DMSO, LPA, WFA and TED-347-treated gastruloid discs at 48h of BMP4 stimulation (n and N as in c, see Methods for details), coloured by GATA3 (f) or SOX2 (g) expression. (**h**) Palantir pseudotime shown on the UMAP embedding for amnion-like cells, defined as a CellRank fate probability > 0.5, in DMSO, LPA, WFA or TED-treated gastruloid discs at 48h of BMP4 stimulation (n and N as in c, see Methods for details). These cells are more likely to reach the amnion-like terminal state than any alternative fate. (**i,j**) Expression of GATA3 (i) and GABRP (j) along pseudotime for cells committed to an amnion-like lineage (CellRank fate probability > 0.9) in gastruloid discs treated with DMSO at 48h of BMP4 stimulation (n and N as in e, see Methods for details). **k.** UMAP denoting ten cell clusters in gastruloid discs stimulated with BMP4 for 48h (dataset and annotations from^41,42^, see Methods for details). PGCLC: primordial germ cell-like cells. **l.** Average SCENIC+ regulon activity for extraembryonic amnion-like cells, with TEAD factors highlighted in bold (Supplementary Table 3, see Methods for details). (**m**) Dot plot showing the activity of the inferred TEAD1/3 regulons in cells displaying an amnion-like signature, defined based on the gene signature enrichment scores (AUCell > 0.19), in LPA, WFA or TED-347-treated gastruloid discs vs. DMSO at 48h of BMP4 stimulation (n and N as in (c), see Methods for details). Dot colour represents the mean change in expression, with dark red indicating positive and blue showing negative correlation. The -log10(adjusted p-value) is shown below the dots. LPA (50 µM) was added for 48h together with BMP4, while WFA (1 µM) and TED-347 (5 µM) were added only between 24 and 48h of BMP4 stimulation (c-l). Scale bars: 100 μm (a). Error bars are shown as mean ± SEM (b). Mann-Whitney test. ****P<0.0001 (b).

**Extended data figure 10:**
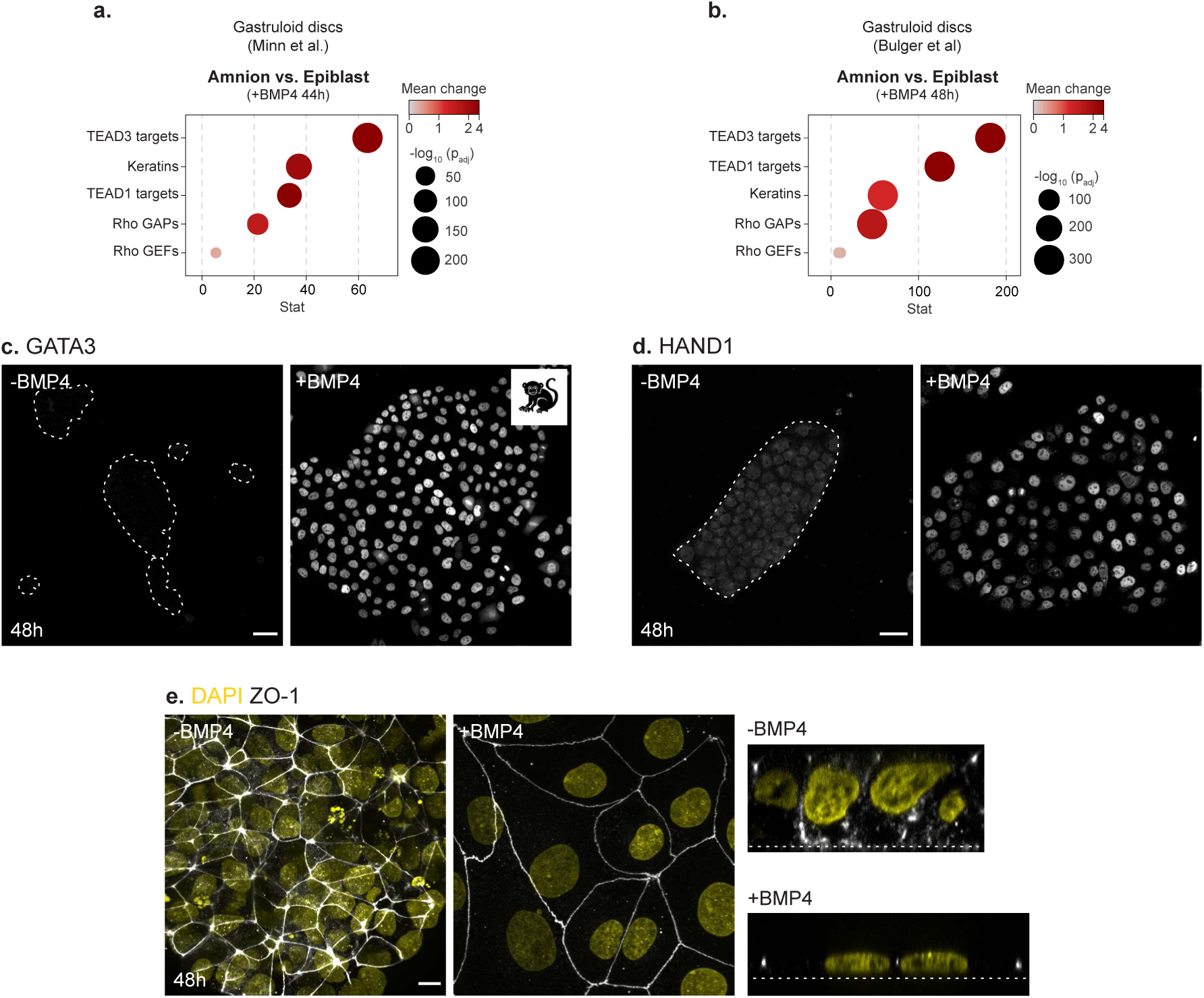
BMP4-induced marmoset PSCs upregulate well-established amnion markers by 48h. (**a,b**) Difference in transcriptional activity for the cytoskeletal switch in amnion-like vs. epiblast cells in across all available scRNA-seq datasets of gastruloid discs after 44h^39,40^ (a) or 48h of BMP4 stimulation (b)^41,42^. Dot colour represents the mean change in expression, with dark red indicating higher activity and grey lower. Dot size indicates the significance of the difference, shown as - log10(adjusted p-value), with larger dots indicating greater significance. (**c,d**) Immunostaining of GATA3 (grey, c), or HAND1 (grey, d) in unconfined marmoset colonies that were either left untreated (-BMP4; GATA3: n=3758 cells, N=1; HAND1: n=1926 cells, N=1) or stimulated with BMP4 ligands (+BMP4; GATA3: n=4377, N=1; HAND1: n=3640, N=1) for 48 h (see Methods for details). Dashed lines outline the unstimulated colonies. **e.** Immunostaining of ZO-1 (grey) and DAPI (yellow) in unconfined marmoset colonies that were either left untreated (-BMP4) or stimulated with BMP4 ligands (+BMP4) for 48 h. Side view is shown on the right. Maximum projections are shown for cell fate markers and nuclei. Scale bars: 50 μm (c,d), 10 μm (e).

## Methods

### Human embryonic stem cell (hESC) line and cell culture

All experiments were conducted using the H9 hESC line, purchased from WiCell (WA09; RRID: CVCL_9773, female). The karyotype and genomic integrity checks were performed by the manufacturer. Pluripotency monitoring (SOX2+, OCT4+ and NANOG+) and routine mycoplasma screening were performed during this study. All the reporter cell lines were generated in the H9 background (see below for details). Cells were cultured in feeder-free conditions, seeded in six-well plates coated with (growth-factor reduced) Matrigel (Corning, 1:100 in DMEMF12) and maintained in chemically defined medium (Essential 8, Thermofischer, A1517001) at 37°C with 5% CO_2_ under humidified conditions. Cells were fed daily with E8 media and passaged in clumps every 3-4 days by incubation with PBS-EDTA for 3 min, at room temperature.

### Stable cell line generation

To generate the live-cell reporters expressing LifeAct::mRFP1 and EGFP::NMHCIIA (Extended data figure 4a) lines, feeder-free H9 cells were engineered with TALEN technology to insert the reporter constructs into the AAVS1 safe-harbour locus, as described previously^134^. The CAG promoter was chosen to drive constitutive expression of the reporter constructs^135^ and donor plasmids were designed as follows: (i) CAG > LifeAct::mRFP1-SV40, and (ii) CAG > EGFP::NMHCIIA-SV40 (Extended data figure 4a). The coding sequence for EGFP::NMHCIIA was amplified from the Addgene plasmid 11680. All donor plasmids were verified by whole-plasmid sequencing prior to electroporation.

For stable cell line generation, hESCs were cultured with a modified in-house media based on the E8 culture system, supplemented with 0.5% BSA (Bovogen, BSAS-AU)^136^. Prior to electroporation, hESCs cultured in in-house E8 media were incubated with 10 μM of Y-27632 (R&D Systems, 1254/10 or Sigma-Aldrich, Y0503) for 2 h. DNA nucleofection was performed with the 4D-Nucleofector (Lonza) system according to the manufacturer’s protocol using the P3 Primary Cell 4D-Nucleofector Kit (Lonza). 24 h after electroporation, media was replaced by fresh media, and the transfection efficiency was first assessed by fluorescence microscopy 2-3 days later. To avoid false positives, 7 days after electroporation, edited cells were sorted for EGFP and/or dual expression of RFP and EGFP by Fluorescence-Activated Cell Sorting (FACS). After sorting, cells were incubated with in-house produced medium containing 0.5% BSA, 10 μM of Y-27632, Penicillin-Streptomycin (PS, 1:100) and Normocin (1:500) for the next 4-5 days to prevent contamination. Next, a SH800S Cell Sorter (Sony) equipped with Cell Sorter software (version 2.2.4.5150) was used to sort single cells into individual wells of 96-well plates, using low-pressure conditions to minimize mechanical stress. Sorted single cells were kept in-house produced media supplemented with 0.5% BSA, 1x CloneR2 (STEMCELL Technologies, 100-0691), PS (1:100) and Normocin (1:500) to support viability and genomic integrity, as well as prevent contamination.

4-6 clones were then selected by high-resolution imaging for further expansion and validation. 2 clones were selected for STR testing, whole genome SNP genotyping and directed differentiation assays (StemMACS Trilineage Differentiation Kit, Cat. No 130-115-660, Miltenyi Biotec) and compared with the parental H9 line. DNA was extracted using the QuickExtract DNA Extraction Solution (Biozym, 101094) for genotyping, STR testing and whole genome SNP genotyping.

### Generating micropatterned gastruloid discs and small, unconfined hESC colonies

In line with previous work^137^, hESCs were maintained in E8 medium and passaged 4 days prior to seeding onto circular micropatterned substrates with a 350 μm radius, when a confluency of 70-80% was reached. Circular micropatterns were generated using either ultraviolet lithography on 35 mm Ibidi dishes (Ibidi, 81156) according to the manufacturer’s instructions (Primo, Alvéole) or directly purchased from CYTOO (96 DC-S-A, glass-bottom 96-well microplates). Micropatterned dishes were then coated with Laminin-521 at 5 µg/ml (Biolamina, LN521) diluted in DPBS+/+ (with Calcium and Magnesium, ThermoFisher Scientific, 14040133). For 35 mm dishes, we used 2 ml of coating solution, while for 96-well plates, only 200 µl were used per well. Unattached laminin was then washed out with DPBS+/+ 4 times.

Cells were dissociated with Accutase (ThermoFischer Scientific, 00-4555-56), pelleted by centrifugation (120 rcf for 5 min, at 21°C) and resuspended in E8 with 10 μM of Y-27632. Using an automated counter Countess 3FL (Invitrogen) for cell counting, a solution with a total of 1.000.000 cells was prepared and seeded onto micropatterned 35 mm dishes. In CYTOO plates, only 100.000 cells were plated per well. After 1h, unattached cells were washed away with E8 media supplemented with 10 μM Y-27632 and 1:1000 PS. Cells were kept in this media for another hour, after which the media was once more exchanged by fresh E8 containing only PS. 12-14 h later, BMP4 ligands were added at a concentration of 50 ng/ml (R&D, 314-BP-050/CF) to the micropatterned colonies in fresh E8 media with PS. For unstimulated colonies, the same protocol was followed with the exception that the media did not contain BMP4 ligands.

To generate small, unconfined colonies, cells were dissociated with Accutase, pelleted by centrifugation (120 rcf for 5 min, at 21°C) and resuspended in E8 with 10 μM of Y-27632. A solution with a total of 50.000 cells was prepared in 500 µl of E8 and seeded onto 35 mm dishes (Ibidi, 81156) previously coated with Laminin-521, as described for gastruloid discs above. Unstimulated colonies were kept in E8 media, while amnion-induced colonies were supplemented with BMP4 ligands at a concentration of 50 ng/ml.

### Immunofluorescence staining

Cells were fixed in 4% paraformaldehyde (Ted Pella, 18505) for 20 min at room temperature and then washed with PBS twice. Permeabilization and blocking were performed with 3% Donkey serum (Jackson ImmunoResearch, 017-000-121) in PBS supplemented with 0.1% Triton X-100 (Sigma, X100) for 60 min, at room temperature. The primary antibodies were diluted in blocking solution as indicated in Supplementary Table 1 and incubated either for 2 h at room temperature or overnight at 4 °C. After primary antibody incubation, cells were washed 3 times with PBS supplemented with Tween-20 (1:1000) and then incubated with secondary antibodies diluted in blocking solution at 1:500 (Supplementary Table 1) with DAPI (1:3000) and incubated at room temperature for 30 min. Finally, the samples were washed once with PBS supplemented with Tween-20 (1:1000) and then 3 times only in PBS. Cells were kept and imaged in PBS.

### Iterative immunofluorescence staining

Iterative immunofluorescence staining was performed following the procedure described in^104^. Briefly, after the initial round of immunostaining (as detailed above), samples were incubated in PBST (PBS + 0.1% Triton X-100) for, at least 2h at room temperature, or overnight at 4 °C. Antibodies were then eluted using an elution buffer containing 0.5 M L-glycine, 3 M Urea, 3M Guanidinium chloride (GC), and 70 mM TCEP-HCl, adjusted to pH 2.5 with HCl or NaOH. Samples were then incubated in the elution solution overnight at 4 °C to ensure maximal removal of the previously bound antibodies. Following elution, samples were washed extensively with PBS Tween-20 (1:1000) to remove all residual elution buffer and re-blocked in blocking buffer before proceeding with the subsequent staining round as detailed above.

### High-resolution imaging of gastruloid discs and small, unconfined hESC colonies

Imaging of immunostained or live gastruloid discs was performed using a Spinning Disk Confocal Olympus IX3 Series (IX83) inverted microscope, equipped with a Hamamatsu Orca Fusion CMOS camera. The objectives used for gastruloid disc imaging were the 10x/0.4 (Air) WD 3.1 mm objective, 20x/0.8 (Air) WD 0.6 mm or the 100x/1.45 (Oil) WD 0.13 mm. The excitation (ex) and emission (em) filters used were as follows: i) blue 405 ex: ex 365/50; Em LP 397; BSP 395; green 488 ex: Ex 480/20; Em 530/40; BSP 495, red Ex 540/20; Em LP575; BSP 570 and far-red Ex 631/23; Em 700/50; BSP 660. All components are controlled via the CellSense software.

### Live imaging of gastruloid discs

For 3D reconstruction of cell outlines during self-organization, gastruloid discs were generated as previously described and incubated with 1:1000 CellMask (Invitrogen, C10046) as in^72^and imaged using the 100X objective with a z-step of 0.2 μm or 0.34 μm. After high-resolution imaging, the gastruloids were fixed and stained for established cell fate markers (CDX2, BRA, SOX2) to ensure that proper patterning was achieved despite incubation with CellMask. An Okolab incubation system was used to maintain temperature at 37°C and a 5% CO_2_ atmosphere.

### FLIM imaging of Flipper-TR

Flipper-TR (Spirochrome, SC020) was dissolved in DMSO at 1 mM, as indicated by the manufacturer. Gastruloid discs were labelled with a 1:1000 dilution of the Flipper-TR stock solution in E8 media supplemented with BMP4 and PS (see above for details). Cells were incubated with the probe for 1h at 37 °C and imaged without washout, as described in ^72,73^.

FLIM imaging was performed using an Olympus IX83 inverted microscope, equipped with a UplanSApo 100x, NA = 1.4 (Oil). The excitation was performed using Picosecond LDH pulsed lasers (Sepia PDL 828, PicoQuant) at 485 nm with 475/435nm filters for the FLIM detector. The photon detector used was a hybrid photomultiplier detection assembly (with <50ps time resolution), controlled via SymPhoTime64 software (PicoQuant). The fluorescence lifetimes of Flipper-TR were extracted from the FLIM images using the SymPhoTime64 software (PicoQuant), as described in^73^ using full images (rather than a region of interest).

### Tether pulling assays using optical tweezers

Glass coverslips were micropatterned using ultraviolet lithography according to the manufacturer’s instructions (PRIMO, Alvéole). Cells were seeded onto the microprinted coverslips, coated with Laminin 521 as described above, and then placed in 35 mm petri dishes. 2 μm-sized carboxyl latex beads, 4% w/v (ThermoFisher Scientific, C37278) were diluted as a drop in 1 mL of E8 media containing PS with or without BMP4 ligands, as indicated in the main text and corresponding figures. The coverslips were mounted on strapped microscopy slides in the bead medium diluted in the induction medium at 1:36. Water and oil for the corresponding objectives were kept at 37°C to avoid degassing.

The optical tweezers measurements were performed on a Nikon X-light V3-01 Spinning disk with an Impetux Sensocell Optical Trap, using a trapping-infrared laser (5W, 1064nm). The trap stiffness was measured as 92 pN/µm. Prior to image acquisition, the traps were used to move away floating dust particles between the plasma membrane and the bead. The bead was then trapped and pushed gently against the plasma membrane until a detectable drop of tension was observed. To ensure the bead was well-attached, the bead was kept in place for 7 secs. Then, membrane tubes of 5 µm were automatically pulled at a speed of 3 µm/sec, followed by a 20 sec hold. In this manner, a position-force curve is recorded (see a representative example in Extended data figure 4i). The static force was measured after tension stabilization as in^78–80^. The tubes were then released from the optical trap to verify that a membrane tether was pulled. The process was repeated manually 1-2 times to verify that the bead retracts back onto the cells. This verification was necessary especially when pulling after 48h of BMP4 induction to confirm the low static force measured.

### Pharmacological treatments

Gastruloid discs were incubated with 20 or 50 µM of LPA (Lysophosphatidic acid, Tocris 3854), 0.75 or 1 µM of Withaferin A (EMD Millipore, 681535), 5 µM TED-347 (MedChem Express, HY-125269) and 20 µM TRULI (Selleckchem, E1061), as indicated in the main text and corresponding figures. The inhibitors were prepared and stored according to the manufacturer’s instructions. The treatments were applied either for the full duration of the experiments or partially, as indicated in the main text and corresponding figures.

### Keratin knockdown with siRNA

For siRNA knockdowns, cells were dissociated to single cells, counted and seeded as for the gastruloid disc protocol described above. Reverse transfection was carried out using Lipofectamine RNAiMAX (Invitrogen, 13778030) and ON-TARGET-Plus pool siRNAs (Dharmacon, Supplementary Table 2) at a concentration of 25nM per siRNA. After overnight incubation media with 10 μM of Y-27632, gastruloid discs were induced with BMP4 following the protocol described above.

### 10x Genomics scRNA-seq

Micropatterned gastruloid discs were generated as described above and incubated with DMSO (1:2000), LPA (50 µM), WFA (0.75 µM) or TED-347 (5 µM). As indicated in the figures and main text, LPA was added together with BMP4 for 48h while DMSO, WFA and TED-347 were added only from 24 to 48h of BMP4 stimulation. For single-cell preparation, gastruloid discs at 48h were washed with PBS and incubated with PBS-EDTA for 3 min, followed by Accutase treatment for 3 min (for discs treated with WFA), 4 min (for discs treated with TED-347), 6 min (for discs treated with LPA), or 7 min (for discs treated with DMSO). For each condition, 10 wells from a 96-well plate were individually dissociated and the resulting cell suspensions were pooled together. Cells were collected by centrifugation (120 rcf for 5 min), mechanical dissociation was performed in PBS by pipetting up and down with a p1000 pipette and the cell suspension was kept on ice. Cell number and cell viability were determined using Solution 18 AO-DAPI staining reagent (ChemoMetec, 910-3018) using a NucleoCounter NC-250. Cells were then fixed using the Chromium Next GEM Single Cell Fixed RNA Sample Preparation Kit (10x Genomics, PN-1000414) according to the manufacturer’s protocol (10x Genomics, CG000478). Single cell RNA profiling was performed using Chromium Next GEM RNA profiling for multiplexed samples V1 workflow according to the manufacturer’s instructions (10x Genomics, CG000527). Libraries were sequenced on NovaSeqX platform (Illumina).

### Marmoset embryonic stem cell and cell culture

The common marmoset (*C. jacchus*) embryonic stem cell line cj367 (female), established at the Wisconsin National Primate Research Centre, was cultured in E8 medium supplemented with E8 supplement (ThermoFisher Scientific, A15171-01), GlutaMAX (Gibco, 35050-061), Lipid mixture 1 (1:100; Sigma Aldrich, L0288), Activin-A (20 ng/mL; R&D System, 338-AC-050) and Glutathione (1.94 μg/mL; Sigma Aldrich, G4251). Cells were grown on laminin-coated plates (5 μg/ml, Biolamina, LN521) and passaged using Accutase, when reaching a confluency of 80%. Cells were kept in 10 μM Y-27632 for the first 24 h post passaging to improve cell survival and maintain pluripotency.

To generate small, unconfined colonies of marmoset PSCs, cells were sparsely seeded on laminin-coated dishes following the same protocol as for hPSCs. Unstimulated colonies were kept in supplemented E8 media as described above for maintenance, while BMP4-stimulated colonies were cultured in E8 medium with E8 supplements only. BMP4 ligands were added at a concentration of 50 ng/ml.

### Quantification and statistical analysis

All image data analysis was performed using Fiji (National Institutes of Health)^138^, MatLab R2023b (license 41134468) and Python.

### Quantification of cell fate markers as a function of gastruloid disc radii

High-resolution images of immunostained gastruloid discs were max projected, and the nuclei, marked by DAPI, were segmented using Stardist^139^. The resulting masks were used to extract the average intensity within each object across all other channels. The centre of mass of all objects was detected to extract their respective spatial coordinates. These were then used to: i) identify the centre of the gastruloid disc (R=0), and ii) systematically exclude all detected objects located beyond 380 µm from the gastruloid disc centre of mass, which is well beyond the radii of the micropatterned circular patterns used in this study. A radial intensity profile was then generated by binning nuclei intensity every 10 µm, calculated from the centre of the gastruloid disc. The background intensity, determined as the median pixel intensity outside of the micropattern, was measured for each channel to check for consistency across experiments, but no additional post-processing steps were performed.

For the analysis of CDX2 and SOX2 intensity across control, LPA, WFA or siRNA-treated gastruloid discs, the average intensity for each marker was averaged in the control conditions of each experiment and used to renormalize the corresponding experimental perturbation. This additional step was necessary in these experiments, because the immunostainings were performed in different channels and/or in a second round of staining. The codes used for these analyses are provided as supplementary files, respectively Stardist_plugin.ijm and MASTER_PLOTTER.m.

### Intensity-based apical projection of gastruloid discs

High-resolution stacks (2304×2304 pixel images with z-step of 1µm) of immunostained gastruloid discs marked for the tight junction component ZO-1 (see details in the “Immunofluorescence staining" section) were pre-processed using the “Bleach Correction” plugin^140^, with the “simple ratio” correction method and the background subtraction function (rolling=2) in Fiji^138^.The pre-processed and raw stacks were then loaded in MATLAB, using the MIJ plugin^141^. Using a custom-made code, pixel intensity and standard deviation was measured in 96 volumes within the image and across all z-planes to identify the local intensity maxima and corresponding z-plane. These were then selected in the original (raw) stacks per volume, projected across 60 µm and averaged across neighbouring volumes to obtain a projection of the local apical surface across the gastruloid disc radii. The code used for this analysis (Intensity-based-z-projection.m) is provided as a supplementary file.

High-resolution stacks (2304×2304 pixel images with z-step of 1µm) of gastruloid discs expressing EGFP::NMHCIIA and immunostained for the tight junction component ZO-1 were used to locally extract the apical plane, based on ZO1 intensity, and the corresponding planes were then retrieved in the EGFP::NMHCIIA channel and projected. These apical projections were used solely for visualization purposes and the details regarding the quantification of EGFP::NMHCIIA intensity are provided below.

### Quantification of local apical height in gastruloid discs or in small, unconfined colonies

Using the z-plane corresponding to the local intensity maxima of ZO-1 per volume, as determined in intensity-based apical projections (see above), we generated local apical height maps per gastruloid disc. The gastruloid edge was detected based on the bounding area of the nuclei contained in the disc. Average maps were then obtained by averaging local height per pixel across micropatterned colonies with controlled geometry. The code used for this analysis (Plot_average_height_map_correct_edge.m) is provided as a supplementary file.

To determine the apical cell height in small, unconfined hESC colonies, max projections of the ZO1 channel were generated from high-resolution imaging stacks (2304 × 2304 pixels with a z-step of 0.34 µm), using Fiji. 2D ROIs were defined per cell based on the maximum intensity projection of ZO-1 and used to generate corresponding 3D masks across the image stack. Within each masked region, mean fluorescence intensity was measured for the DAPI channel in every z-slice. The intensity profile along the z-axis was used to determine the peak signal position and the basal-most plane per cell, using an intensity-based threshold. The difference between these planes was used to calculate the apical-basal height. This analysis was performed using CellPose^142^ segmentation on the ZO1 channel followed by custom Python scripts. Similar analysis approach was used for small, unconfined colonies of marmoset stem cells.

### Quantification of apical area in gastruloid discs or in small, unconfined colonies

We used CellPose^142^ to segment cell outlines within gastruloid discs, based on ZO-1 projections. This workflow enables manual correction, but this was kept to a minimum given the high quality of the automatic segmentation and the possibility of generating average maps across hundreds of gastruloid discs. The centre of mass of all the segmented objects was detected and used to define the centre of the gastruloid disc. Objects located 380 µm beyond the gastruloid disc centre of mass, which is well beyond the radii of the micropatterned circular patterns used in this study, were systematically excluded from further analysis. To automatically exclude segmentation errors, cells smaller than 30 µm^2^ and larger than 2000 µm^2^ were systematically excluded from further analysis. The resulting masks were individually examined and cases representing obvious errors of under- or over segmentation were excluded. Average apical area maps were obtained by binning cell area per 20 µm from the centre of each gastruloid disc and then averaged across gastruloid discs to plot the resulting measurements onto a circular map. The code used for this was provided as a supplementary file, termed Plot_area_distance_to_center_cellpose.m.

For small, unconfined colonies, Cellpose^142^ was used to segment cell outlines based on the ZO-1 projection and apical cell areas were extracted from the obtained ROIs. This analysis was performed using custom Python scripts. Similar analysis approach was used for small, unconfined colonies of marmoset stem cells.

### 3D reconstruction of cell outlines and apical interfacial angle determination

Gastruloid discs imaged with CellMask were reconstructed using CellPose-SAM^143^. First, the imaging datasets (z-step of 200 or 320 nm) were rescaled and down sampled from the original xy-voxel size of 130 nm to 400 or 320 nm to obtain cubic voxels. To achieve a high-quality segmentation of cell outlines, particularly at basal cell outlines, we further trained the CellPose-SAM model by performing manual annotations. Interfacial apical angles were determined from the interfacial normals, which were fitted in a point-cloud fashion to obtain a voxel representation of the apical and cell-cell interfaces, using ForceIn3D^62^. The apical angles were then spatially averaged by binning the radial positions across the gastruloid section in bins with 20 µm bin size, using Python. The codes used for these analyses are provided as supplementary files, termed Interfacial_apical_angles.py.

### Quantitative analysis of actomyosin and keratin network intensity

High-resolution stacks (2304×2304 pixel images with z-step of 1µm) of gastruloid discs expressing EGFP::NMHCIIA were max projected using Fiji. These images were used for the polar transformation of the EGFP::NMHCIIA signal along the gastruloid radii, using the Polar Transformer plugin from Fiji (at 360 degrees). The polar transform was then aligned on the gastruloid edge, rather than the centre. For this, the polar-transformed image was copied, blurred and thresholded, based on pixel intensity, to create a binary mask. For each angular row, the rightmost non-zero pixel, corresponding to the maximal radial position of the signal, was identified. The average alignment position was defined at the reference radius of 350 µm and the original image was aligned according to the thresholded mask. The mean intensity of EGFP::NMHCIIA as a function of gastruloid disc radii was measured on this aligned kymograph. The intensity of EGFP::NMHCIIA is not shown between 0-100 µm, because the edge alignment often results in black pixels being added at the centre of the gastruloid. Finally, to account for brightness differences across dish types and experiments, intensity of EGFP::NMHCIIA as a function of gastruloid disc radii was normalized by the average signal measured between 100-150 µm in each gastruloid. The code used for this analysis was provided as a supplementary file, termed Polar_kymograph_aligned.ijm.

High-resolution stacks (2304×2304 pixel images with z-step of 1µm) of gastruloid discs immunostained for pan-Keratin were projected using a custom MatLab script to automatically detect the basal-most plane, based on the DAPI channel. Briefly, the stack was subdivided into 96 smaller windows, as previously described for the apical projections based on ZO-1 intensity, and for each window the bottom plane, defined as the lowest brightest plane, was determined locally. This is necessary to correct for small z differences in the gastruloid bottom across the disc. The lower plane was then used as reference to generate a local mean projection of 18, 13 or 4 planes for gastruloid discs respectively at 24, 36 or 48h after BMP4 stimulation. Here a mean, rather than maximum, projection was necessary to account for the striking differences in pan-Keratin intensity across the apical-basal axis, particularly at earlier time points of gastruloid disc self-organization where only an apical network of keratins is detected. The adjustment of the number of planes projected is necessary to project the full apical-basal length of cells at the gastruloid edge over time without averaging black planes. These images were used for the polar transformation of the pan-Keratin signal along the gastruloid radii, using the Polar Transformer plugin from Fiji and aligned at the gastruloid edge as described above. Finally, to account for brightness differences across dish types and experiments, intensity of pan-Keratin as a function of gastruloid disc radii was normalized by the minimum average signal in 50 µm spatial bins in each gastruloid and experimental condition.

### Quantitative analysis of the nuclear-to-cytosolic levels of YAP1

High-resolution stacks (2304×2304 pixel images with z-step of 1µm) of gastruloid discs or small, unconfined colonies (2304×2304 pixel images with z-step of 1µm) immunostained for total YAP1 were maximum projected, using Fiji. The nuclei were segmented, based on DAPI signal, using Stardist^139^. The nuclei masks were expanded by 1.7 pixels to obtain a cytosolic band, which was defined as 1.5 pixels around the segmented and expanded nuclei mask. The code used for this analysis was based on a previous script of Nicholas Condon^144^. The nuclei (defined as objects with an area larger than 30 µm^2^), cytosolic, and resulting nuclei-to-cytosolic ratio of YAP1 intensity was then automatically extracted per ROI and stored for manual inspection. The average nuclei-to-cytosolic ratio of YAP1 intensity was averaged in the control conditions of each experiment and used to renormalize the corresponding experimental perturbation (LPA, WFA or siRNA). This additional step was necessary, because the immunostainings were performed in different channels and/or in a second round of immunostaining. The modified code used for this analysis was provided as a supplementary file, termed YAP_quantification_Stardist.ijm and plot_YAP1_radii.m.

### GRN inference and *in vivo* transcriptional signatures for the cytoskeletal switch

Published raw single-cell transcriptomic (scRNAseq) data of self-organizing gastruloid discs^39,40^ (GSE169074) were reprocessed using Cell Ranger (count v9.0.0), using the prebuild 10X GRCh38 reference (refdata-gex-GRCh38-2024-A). Published raw single-cell multiome dataset of gastruloid discs after 48 h of BMP4 stimulation^41,42^ (GSE245998) were reprocessed using Cell Ranger ARC (v2.0.0) with GRCh38 reference (refdata-cellranger-arc-GRCh38-2020-A-2.0.0) from 10X Genomics as a reference. For both datasets, the cell filtering and annotations provided by the authors were applied to the processed data. Based on the multiome dataset, enhancer-driven gene regulatory networks were inferred via SCENIC+^113^ for control gastruloid discs (nr_cells_per_metacells=15, rho=0.5, Supplementary Table 3).

Preprocessed and annotated scRNAseq data of gastrulating human embryos were downloaded from the Petropoulos & Lanner Labs human embryonic reference tool (https://petropoulos-lanner-labs.clintec.ki.se/dataset.download.html), focusing on the dataset for CS7 gastrula^115^ (Tyser_2021_PMID_34789876.meta.tsv and Tyser_2021_PMID_34789876.counts.gz). Preprocessed and annotated spatial embryo profiling data for CS6 marmoset embryos^31^, which contained more amnion cells, were obtained from https://github.com/Boroviak-Lab/SpatialModelling (*In vivo* dataset: BaseDimRed.Rds).

To analyse genes overrepresented in the Amnion versus Epiblast, we first subsetted all aforementioned datasets for the indicated cell types. Feature set enrichment scores were inferred using decoupleR v2.1.1, via the univariate linear model (ULM) method. Differences in scores between groups were evaluated using the scanpy t test adaptation (t-test_overestim_var). The evaluated feature sets were derived from OmniPath UniProt keywords, extended by four custom sets - the TEAD1 and TEAD3 SCENIC+-derived regulons, based on the multiome dataset of gastruloid discs, keratins, GAPs and GEFs for the GTPase Rho, a well-known regulator of myosin contractility. This list was compiled based on the literature, focusing on GAPs and GEFs with specific activity towards the GTPase Rho (gene list is provided in Supplementary Table 4).

### scRNA-seq analysis of gastruloid discs across cytoskeletal perturbations

Four experimental conditions (DMSO, LPA, WFA and TED-347) were investigated by scRNA-seq using the 10x Genomics Chromium Flex (Fixed RNA Profiling) assay, with two biological replicates per condition (N=2). Samples were processed across two independent Flex experiments, with each experiment including a replicate of all conditions. Sequencing reads were analysed using cellranger multi v9.0.0 and the build in Chromium Human Transcriptome Probe_Set v1.1.0 GRCh38-2024-A.csv. Downstream analyses, including data integration, dimensionality reduction, clustering, and visualization, were performed in R using Seurat v5.1.0. Low-quality cells and doublets were removed on a per-sample basis by excluding: 1) cells with a gene count more than 2 MADs below or above the sample median, and 2) doublets identified with scDblFinder v1.20.2, using default settings. Expression data were log-normalized, scaled regressing out the number of detected genes and mitochondrial transcript percentage and dimensionality reduction was performed using PCA on the top 3000 most variable genes. Batch correction across libraries was performed with Harmony v1.2.3. Neighbour graph construction, clustering, and UMAP visualization were performed on the Harmony-corrected embeddings.

Cells with an amnion-like or epiblast signature were quantified using AUCell v1.28.0 and lineage markers derived from^39,40^, namely using the table elife-59445-data2.xlsx. The AUC was calculated based on the top 5% of ranked genes, and cells with an AUCell score exceeding the threshold calculated from a single Gaussian fit to the global AUC distribution were classified to the respective lineage, respectively 0.19 for amnion-like and 0.33 for epiblast. Differential feature set enrichment analysis was performed on cells with an amnion-like or epiblast signature. Briefly, feature set activity scores were estimated using the univariate linear model (ULM) method implemented in decoupleR v2.1.1, and differential activity between groups (LPA, WFA or TED-347 vs. DMSO in cells with an amnion-like signature; or amnion-like vs. epiblast for the DMSO control) were assessed by t test on ULM scores.

Cell types were annotated based on clusters identified using the FindClusters function with the Leiden algorithm (algorithm = 4) at a resolution of 0.2. The mesoderm and ectoderm populations were defined using clusters obtained at a higher clustering resolution to achieve finer separation of these two populations.

Trajectory inference was performed in Python using CellRank 2.3.4.dev3^112^ and Palantir v1.4.5^111^. The Harmony-corrected embedding was used to build a k-nearest neighbour graph (k=8), from which diffusion maps and a multiscale diffusion space were computed. Pseudotime was then estimated with Palantir^111^, using as a biologically informed starting point the cell nearest the centroid of the population with the highest smoothed Epi-like score, defined as described above. Palantir pseudotime was supplied to the CellRank PseudotimeKernel to compute a directed cell-cell transition matrix. Transitions were visualized as stream plots on the UMAP embedding. A Schur decomposition was computed with the GPCCA estimator, from which five macrostates were identified. The initial state was set to the macrostate containing the Palantir start cell, and all remaining macrostates were set as terminal states, and fate probabilities toward the terminal states were computed. Cells were considered biased toward the amnion-like lineage if their fate probability for the amnion-like terminal state was > 0.5, and strongly committed to it if the probability was > 0.9. To resolve the amnion-specific trajectory, cells biased towards the amnion terminal state were reprocessed using the same settings as for the full dataset (normalization, highly variable gene selection, batch harmonization, dimensionality reduction, neighbor graph construction and pseudotime inference).

### Statistics and reproducibility

The number of biological (n) and experimental replicates (N) is indicated in the corresponding figure legends. The statistical analyses and plots were performed using GraphPad Prism and the error bars are shown as SEM or SD as indicated in the corresponding figures and figure legends. The statistical tests used to assess significance are indicated in the figure legends, along with the corresponding P values. To choose an appropriate statistical test, the distribution of each group was first analysed using the D’Agostino-Pearson normality test. To compare two groups, a two-sided t test or a Mann–Whitney test was used, depending on whether either data set shows (or not) normal distribution. In cases where a t-test was used, the variances were subsequently tested using the F-test. If significantly different, a Welch t test was used instead. To compare multiple conditions, an ANOVA or Kruskal-Wallis test was used, depending on whether either data set shows (or not) normal distribution.

---

**Supplementary Table 1:**
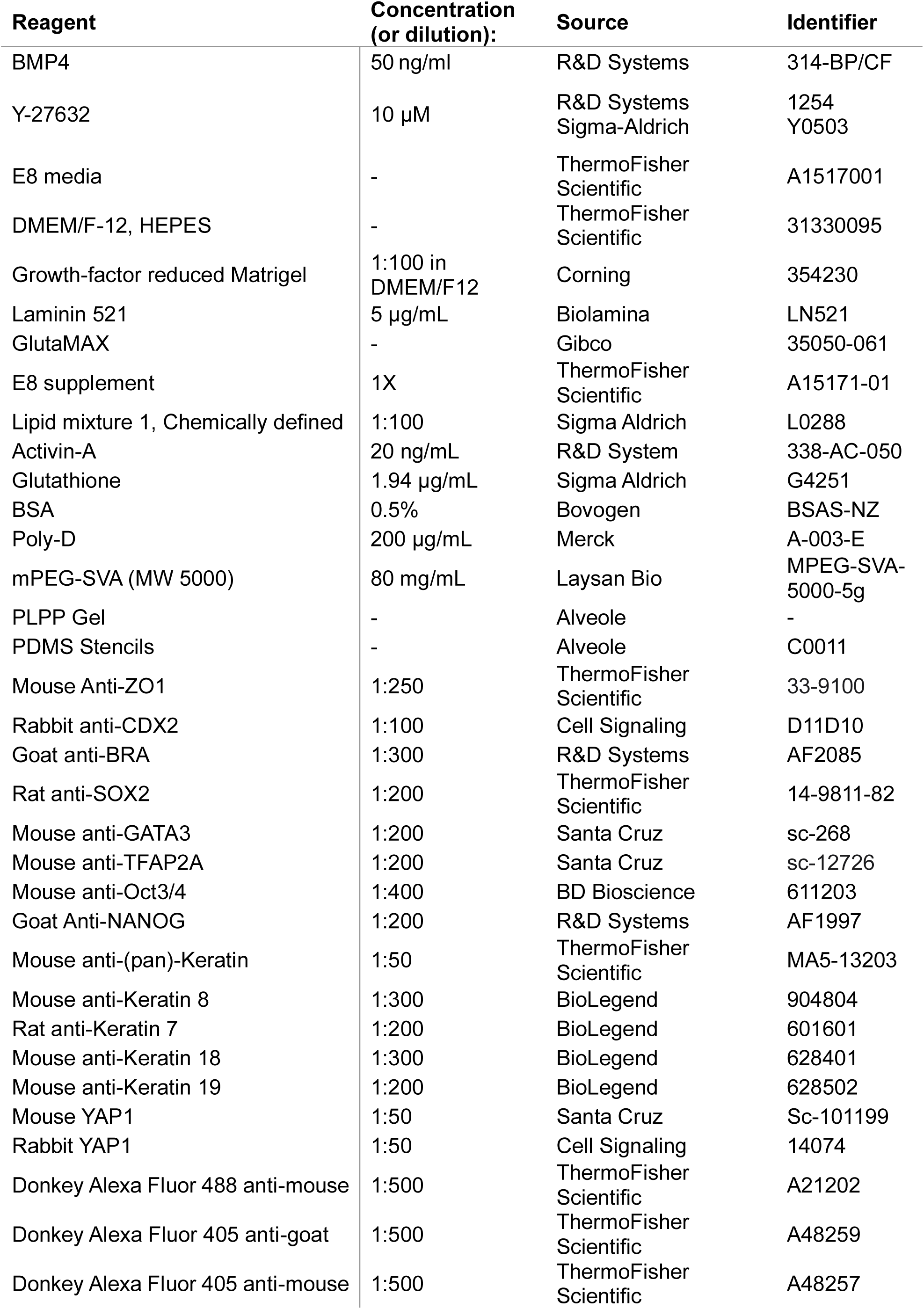
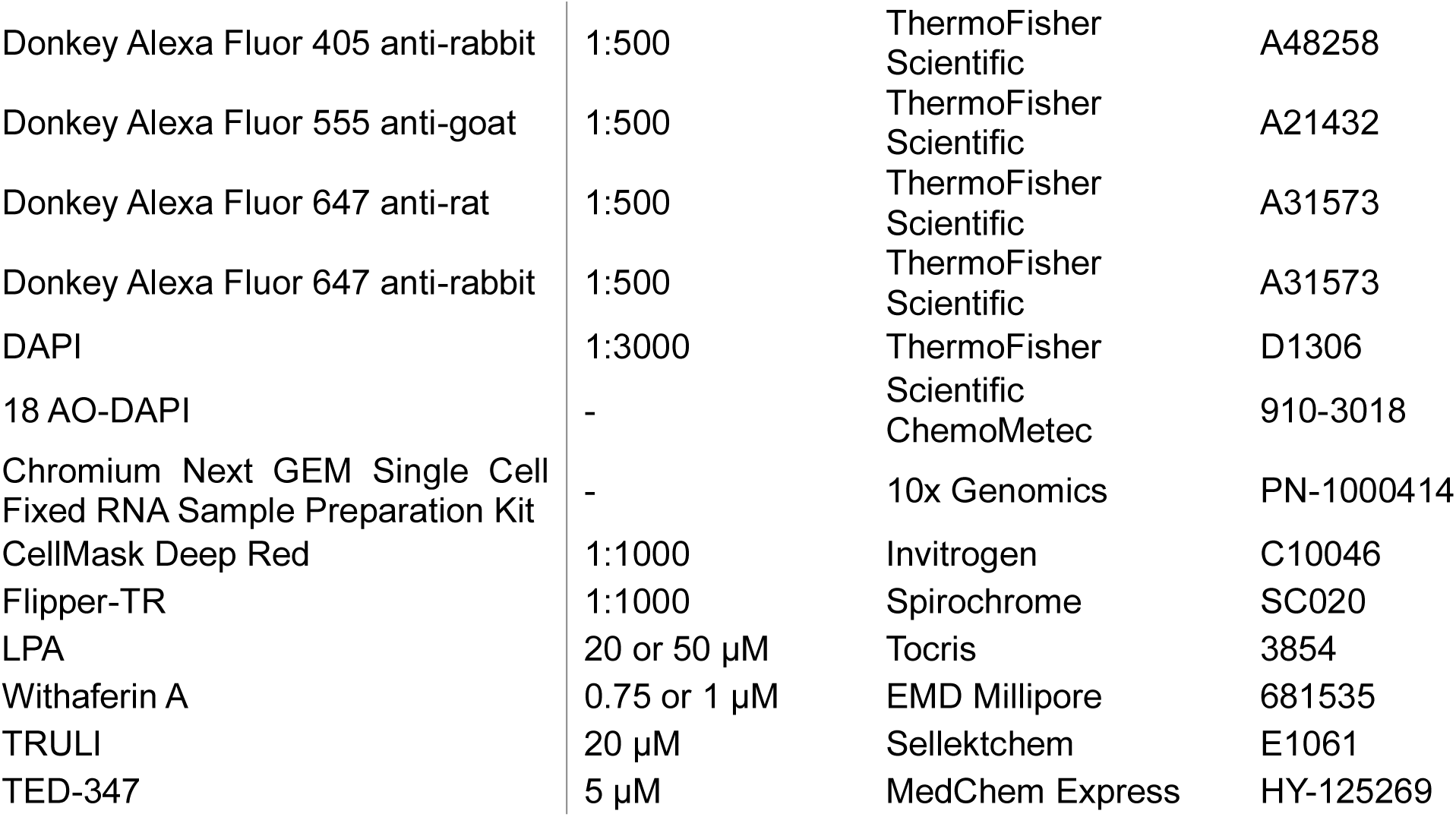
List of key reagents or resources.

| Reagent | Concentration (or dilution): | Source | Identifier |
|---|---|---|---|
| BMP4 | 50 ng/ml | R&D Systems | 314-BP/CF |
| Y-27632 | 10 µM | R&D Systems<br>Sigma-Aldrich | 1254<br>Y0503 |
| E8 media | - | ThermoFisher Scientific | A1517001 |
| DMEM/F-12, HEPES | - | ThermoFisher Scientific | 31330095 |
| Growth-factor reduced Matrigel | 1:100 in DMEM/F12 | Corning | 354230 |
| Laminin 521 | 5 µg/mL | Biolamina | LN521 |
| GlutaMAX | - | Gibco | 35050-061 |
| E8 supplement | 1X | ThermoFisher Scientific | A15171-01 |
| Lipid mixture 1, Chemically defined | 1:100 | Sigma Aldrich | L0288 |
| Activin-A | 20 ng/mL | R&D System | 338-AC-050 |
| Glutathione | 1.94 µg/mL | Sigma Aldrich | G4251 |
| BSA | 0.5% | Bovogen | BSAS-NZ |
| Poly-D | 200 µg/mL | Merck | A-003-E |
| mPEG-SVA (MW 5000) | 80 mg/mL | Laysan Bio | MPEG-SVA-5000-5g |
| PLPP Gel | - | Alveole | - |
| PDMS Stencils | - | Alveole | C0011 |
| Mouse Anti-ZO1 | 1:250 | ThermoFisher Scientific | 33-9100 |
| Rabbit anti-CDX2 | 1:100 | Cell Signaling | D11D10 |
| Goat anti-BRA | 1:300 | R&D Systems | AF2085 |
| Rat anti-SOX2 | 1:200 | ThermoFisher Scientific | 14-9811-82 |
| Mouse anti-GATA3 | 1:200 | Santa Cruz | sc-268 |
| Mouse anti-TFAP2A | 1:200 | Santa Cruz | sc-12726 |
| Mouse anti-Oct3/4 | 1:400 | BD Bioscience | 611203 |
| Goat Anti-NANOG | 1:200 | R&D Systems | AF1997 |
| Mouse anti-(pan)-Keratin | 1:50 | ThermoFisher Scientific | MA5-13203 |
| Mouse anti-Keratin 8 | 1:300 | BioLegend | 904804 |
| Rat anti-Keratin 7 | 1:200 | BioLegend | 601601 |
| Mouse anti-Keratin 18 | 1:300 | BioLegend | 628401 |
| Mouse anti-Keratin 19 | 1:200 | BioLegend | 628502 |
| Mouse YAP1 | 1:50 | Santa Cruz | Sc-101199 |
| Rabbit YAP1 | 1:50 | Cell Signaling | 14074 |
| Donkey Alexa Fluor 488 anti-mouse | 1:500 | ThermoFisher Scientific | A21202 |
| Donkey Alexa Fluor 405 anti-goat | 1:500 | ThermoFisher Scientific | A48259 |
| Donkey Alexa Fluor 405 anti-mouse | 1:500 | ThermoFisher Scientific | A48257 |
| Donkey Alexa Fluor 405 anti-rabbit | 1:500 | ThermoFisher Scientific | A48258 |
| Donkey Alexa Fluor 555 anti-goat | 1:500 | ThermoFisher Scientific | A21432 |
| Donkey Alexa Fluor 647 anti-rat | 1:500 | ThermoFisher Scientific | A31573 |
| Donkey Alexa Fluor 647 anti-rabbit | 1:500 | ThermoFisher Scientific | A31573 |
| DAPI | 1:3000 | ThermoFisher | D1306 |
| 18 AO-DAPI | - | Scientific ChemoMetec | 910-3018 |
| Chromium Next GEM Single Cell Fixed RNA Sample Preparation Kit | - | 10x Genomics | PN-1000414 |
| CellMask Deep Red | 1:1000 | Invitrogen | C10046 |
| Flipper-TR | 1:1000 | Spirochrome | SC020 |
| LPA | 20 or 50 µM | Tocris | 3854 |
| Withaferin A | 0.75 or 1 µM | EMD Millipore | 681535 |
| TRULI | 20 µM | Sellektchem | E1061 |
| TED-347 | 5 µM | MedChem Express | HY-125269 |

**Supplementary Table 2:**
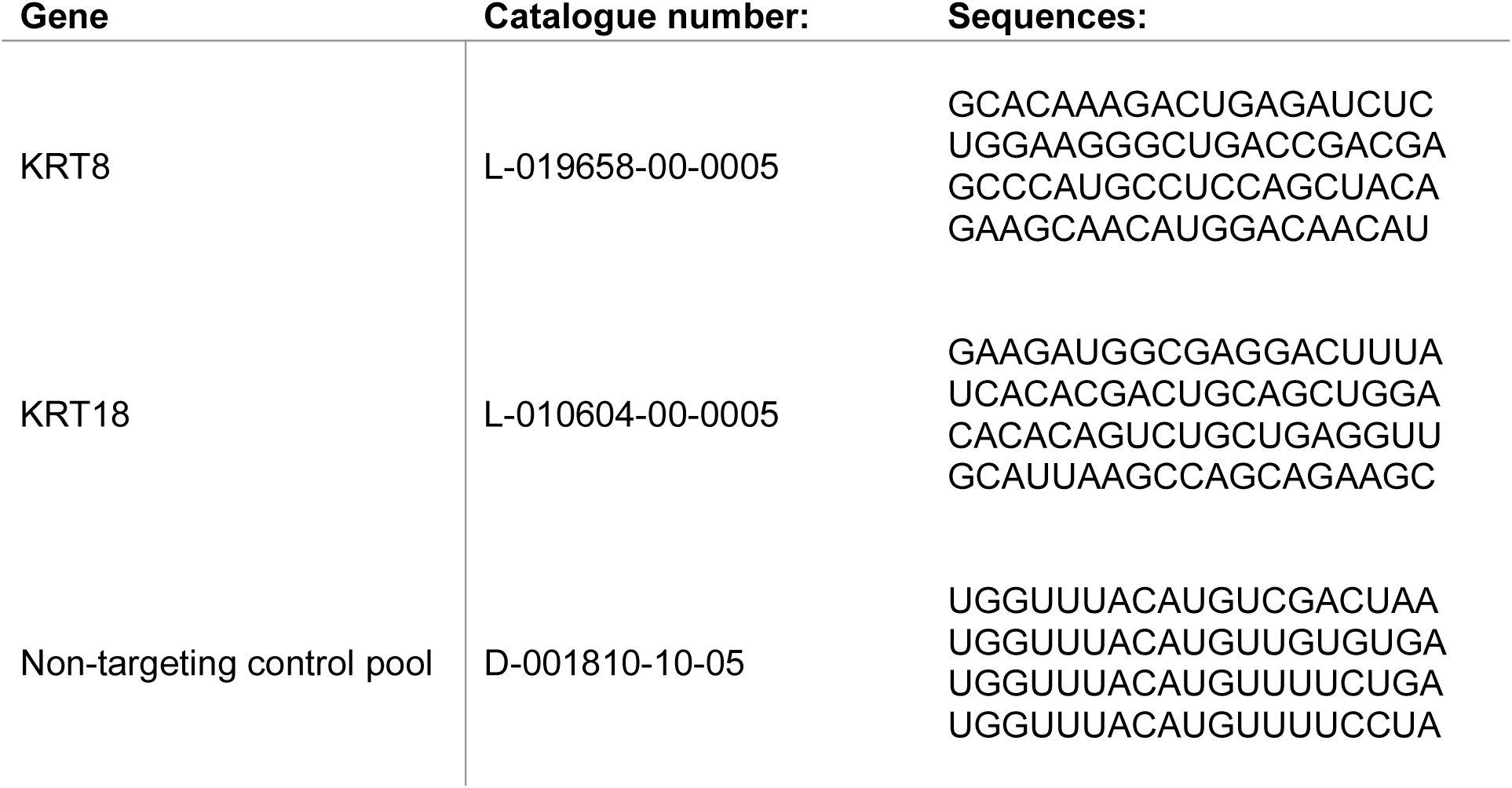
List of siRNAs used in this study.

**Supplementary Table 3: List of genes in the inferred TEAD regulons.**

Provided as an excel file.

**Supplementary Table 4: List of GEFs and GAPs for the GTPase RhoA.**

Provided as an excel file.

## Notes

### Competing Interest Statement

The authors have declared no competing interest.

