## Supplementary material for "A cytoskeletal toggle switch controls amnion architecture and fate": Roffay_et-al_Theory-Note

### Theory Note for:
### A cytoskeletal toggle switch controls amnion architecture and fate

In this Theory Note, we give additional details on the modelling approach used in the main text, as well as on parameter estimation and fitting. The two sections correspond to the two length scales of the modelling, i.e. the vertex model of the amnion domain, and the keratin-myosin switch within amnion cells.

#### 1. Three-dimensional bubbly vertex model of an amnion cell layer

##### Model definition

To theoretically model the columnar-to-squamous transition in the amnion, we consider a three-dimensional vertex model approach, where the individual interfaces between neighbouring cells (lateral) and the cells and the medium (apical) are subject to surface tensions [1]. On the basal side (interface between cell and extracellular matrix) we assume an active stress, which comes from basal contractile tension and a basal adhesion energy [2]. The total energy of a representative cell in the monolayer thus reads:

$$E = \sum_{\text{cells}} \left( \Gamma_a A_a - \sigma_{\text{act}} A_b + \frac{\Gamma_l}{2} A_l \right), \tag{T1}$$

with apical and lateral surface tensions, $\Gamma_a$ and $\Gamma_l$, respectively; basal active stress $\sigma_{\text{act}}$; and corresponding areas $A_a$, $A_b$, and $A_l$. We assume a set cellular volume of $V_{\text{cell}}$, as the osmotic forces that regulate volume are much larger than actomyosin tensions [3].

Going beyond classical vertex model approaches [4], we also allow in this section for interfacial curvature (the so-called bubbly vertex model) [1], as this is both a more realistic assumption, but also provides, via the interfacial angles it introduces, additional degrees of freedom with which inference of relative tensions can be performed [5].

Assuming a hexagonal lattice structure, the shape of a representative cell in the monolayer can be approximated as a hexagonal prism with a spherical cap as apical side, with the hexagon's in-circle as base, see Fig. 2c and Extended Data Fig. 3a. With hexagonal edge length $s$, prismatic height $h'$, hexagonal in-radius $r$, spherical cap height $h''$, and geometric constant $\zeta_{\text{geom}} = 3^{3/2}/2$ the areas are:

$$A_a = \pi \left( r^2 + h''^2 \right), \tag{T2}$$

$$A_b = \zeta_{\text{geom}} s^2, \tag{T3}$$

$$A_l = 6h's. \tag{T4}$$

The spherical cap has a radius of $R$, which is implicitly defined by the tensions via force balance at the apico-lateral tri-interfacial intersection lines (Fig. 2b, Extended Data Fig. 3a) and the volume constraint. The force balance implies:

$$\frac{\Gamma_l}{2\Gamma_a} = \sin(\alpha) = \frac{r}{R} \tag{T5}$$

with the angle between the apical interface and the horizontal reference $\alpha$. It is related to the apical angle $\varphi$ (between the apical interfaces) via $\varphi/2 = \pi/2 - \alpha$. Non-dimensionalizing the volume by $V_{\text{cell}}$, i.e. setting

$V_{\text{cell}} \equiv 1$, we obtain for the volume:

$$V_{\text{prism}} = A_b h', \tag{T6}$$
$$V_{\text{cap}} = \frac{\pi}{3} R^3 \left(2 + \cos\alpha\right)\left(1 - \cos\alpha\right)^2, \tag{T7}$$
$$1 = V_{\text{prism}} + V_{\text{cap}}. \tag{T8}$$

This yields for the cap radius:

$$R^3 = \frac{3\left(1 - \zeta_{\text{geom}} s(R)^2 h'\right)}{\pi\left(2 + \cos\alpha\right)\left(1 - \cos\alpha\right)^2}, \tag{T9}$$

where only the prism height $h'$ is a free parameter because all other geometric quantities are defined through the angle, which we obtain from the tensions, and this radius, i.e.

$$s = \eta_{\text{geom}} r = \eta_{\text{geom}} R \sin\alpha, \tag{T10}$$
$$h'' = R\left(1 - \cos\alpha\right), \tag{T11}$$

where the geometric constant $\eta_{\text{geom}} = 2/\sqrt{3}$ relates the hexagonal edge to in-radius length.

Now inserting the areas into the energy allows us to numerically determine the minimum-energy configuration. For this, we numerically solve for the cap radius $R$ as a function of $h'$ and then minimize the total energy with respect to $h'$ for given surface tensions.

#### Shape changes via differential actomyosin localization

At each apical edge shared by two neighboring cells in the 3D bubbly vertex model, force balance implies that the sum of the vertical components of the two apical tensions has to be equal to the lateral tension (which always points downward). In order to achieve this, apical tensions have to be large enough, i.e.

$$2\Gamma_a \geq \Gamma_l. \tag{T12}$$

Otherwise, even an apical inclination angle of $\alpha = \pi/2$ (apical angle $\varphi = 0$, apical tension being purely vertical at the edge) would not be sufficient to balance lateral stresses, meaning that cells would de-adhere from each other. Note that these effects are neglected in classical, non-bubbly, vertex models, meaning that squamous transitions can be achieved by increasing $\Gamma_{\text{l}}/\Gamma_{\text{a}}$ to arbitrarily large values [2, 6, 7]. Yet, we show here that this is incorrect from a physical perspective, as it is inconsistent to neglect curvature if $\Gamma_{\text{a}}$ is not much larger than $\Gamma_{\text{l}}$. Note that past examples where lateral tension changes were proposed to contribute to shape changes involved highly columnar cells [2, 4, 7], in which case this assumption is warranted, although it fundamentally fails in our case of a squamous transition.

In the extreme case where the apical tensions exactly balance lateral tension $2\Gamma_a = \Gamma_l$, the interfacial angle becomes $\alpha = \pi/2$, which corresponds to the case that the apical spherical cap is a half sphere with a vertical tangent at the contact points. In this limit, the cap volumes and areas simplify and we can solve for the cap radius and height directly,

$$R = h'' = \left(\frac{3V_{\text{cap}}}{2\pi}\right)^{1/3} = r = \frac{\sqrt{3}}{2} s, \tag{T13}$$

which yields for the equilibrium height:

$$h'_{\text{eq}} = \frac{2}{3^{3/2}} s_{\text{eq}}^{-2} - \frac{\pi}{3} s_{\text{eq}}, \tag{T14}$$

with equilibrium edge length:

$$s_{\mathrm{eq}} = \left( -\frac{9}{2\pi} \frac{\sigma_{\mathrm{act}}}{\Gamma_{\mathrm{l}}} + \frac{3^{3/2}\pi}{2} \frac{\Gamma_{\mathrm{a}}}{\Gamma_{\mathrm{l}}} - \sqrt{3}\pi \right)^{-1/3} . \tag{T15}$$

We also find that this height approaches zero for decreasing (potentially negative) basal stress, when the total height reaches $R$. Decreasing $\sigma_{\mathrm{act}}$ further is incompatible with the apico-lateral force balance condition, while increasing this basal stress leads to more columnar taller cells.

In case of a vanishing prism, i.e. $h' = 0$, if we drop this apico-lateral force balance by considering a single cell which does not interact mechanically with its neighbours, we obtain a spherical cap with force balance between the apical interface, which pulls inward and the basal side, which pushes outward (for negative basal stress resulting from adhesion). This is a classical wetting problem, where we obtain full wetting for $\sigma_{\mathrm{act}} > \Gamma_{\mathrm{a}}$. This effect yields a discontinuity upon cellular flattening, as it is necessary to mechanically split cell-cell contacts (Extended Data Fig. 3e,f) to achieve full wetting ($h \to 0$).

The opposite case of a flat apical interface and a prismatic cell is achieved for $\Gamma_{\mathrm{a}} \gg \Gamma_{\mathrm{l}}$, where the angle becomes $\alpha \approx 0$ and the cap radius diverges $R \gg 1$. This is approximately the geometry of a classical three-dimensional vertex model with completely flat interfaces, allowing us to directly solve for the equilibrium height,

$$h_{\mathrm{eq}} = \frac{2^{1/3}}{3^{1/6}} \left( \frac{\Gamma_a - \sigma_{\mathrm{act}}}{\Gamma_l} \right)^{2/3} . \tag{T16}$$

If we allow for negative basal stress from adhesion, then in this limit the vertex model undergoes a transition to a completely wetted state with $h_{\mathrm{eq}} = 0$ at $\sigma_{\mathrm{act}} = -\Gamma_a$.

#### Shape changes via global tension drop

Our theoretical result suggests that flattening can be achieved by increasing lateral tension or decreasing apical tension but the extent of flattening is bounded by the limit of a vanishing apical angle, when cells fully round up (Extended Data Fig. 3c-f). Cell height is then determined by volume conservation as this implies upward apical tangential angles at the edge. Alternatively, cell height can also be decreased by decreasing basal stress, which effectively means that the balance of basal tension and adhesion is reduced, which enables a full squamous transition without changing the apical angle (Extended Data Fig. 3g,h).

To capture realistic shapes for the observed experimental situation, where we find spreading with reduced overall actomyosin levels and tensions (Fig. 2f,g,l, Extended Data Fig. 4a-i), we therefore explicitly separate the basal stress into a contractile basal tension $\Gamma_{\mathrm{b}}$ and an adhesion energy density $\sigma_{\mathrm{adh}}$ on the basal side, i.e. $-\sigma_{\mathrm{act}} = \Gamma_{\mathrm{b}} - \sigma_{\mathrm{adh}}$, so that the Hamiltonian becomes:

$$E = \sum_{\text{cells}} \left( \Gamma_a A_a + \Gamma_b A_b - \sigma_{\mathrm{adh}} A_b + \frac{\Gamma_l}{2} A_l \right) . \tag{T17}$$

With this formulation, we can model the decrease in overall tension by simply rescaling all tensions in the system by a factor $m$, leaving constant the contribution from basal adhesion:

$$\Gamma_a \to m\Gamma_a, \tag{T18}$$

$$\Gamma_l \to m\Gamma_l, \tag{T19}$$

$$\Gamma_b - \sigma_{\mathrm{adh}} \to m\Gamma_b - \sigma_{\mathrm{adh}}. \tag{T20}$$

As we show in Extended Data Fig. 3i,j, a tension reduction in a free cell, i.e. decreasing $m$, yields cell flattening. This is consistent with unconfined colonies undergoing such flattening, without the mechanical influence of other tissue (Fig. 2i,j, Extended Data Fig. 3i,j).

#### Finite-size corrections for gastruloid discs

The above description, which considers only the force balance of amnion cells, should directly hold when examining unconfined colonies (Fig. 1e-i, Fig. 2j). When cells flatten, their height decreases and their covered surface area increases, as we assume volume conservation, which fits well the experimental findings from small unconfined colonies upon BMP4 stimulation. However, on a micropattern, such as in the gastruloid discs, flattening of one part of the monolayer and a concomitant surface area increase must be accompanied by the compression of another part of the monolayer, which in turn means the surface area decreases in the latter. When fitting the amnion shape changes within gastruloid discs, we therefore need to consider the compression of the embryonic core domain, which will mechanically resist the amnion flattening. This can be described via an additional pressure against which the amnion has to flatten. Further, cell division will also contribute to this pressure, as more cells in a monolayer will be accommodated by an increased height, driving the cell shape out of its equilibrium shape determined purely from tension energy minimization.

In order to calculate the corresponding pressure, we consider the mechanical work which is done on a single hexagonal cell in the limit of flat interfaces ($h'' = 0$) upon in-plane area changes [8], i.e. we consider:

$$\mathrm{d}e_{\mathrm{cell}} = P\,\mathrm{d}A_a. \tag{T21}$$

where we can insert the single cell energy,

$$e_{\mathrm{cell}} = \Gamma_{\mathrm{a}} A_{\mathrm{a}} + (\Gamma_{\mathrm{b}} - \sigma_{\mathrm{adh}}) A_{\mathrm{b}} + \frac{6}{2} \Gamma_{\mathrm{l}} h s, \tag{T22}$$

with edge length $s$ and height $h = h'$. Non-dimensionalizing length by cellular volume, i.e. setting $V_{\mathrm{cell}} = 1$, we have for a hexagonal cell $A_{\mathrm{b}} = A_{\mathrm{a}} = \frac{3^{3/2}}{2} s^2$ and $h = 1/A_{\mathrm{a}}$, which yields:

$$\mathrm{d}e_{\mathrm{cell}} = \left( \Gamma_{\mathrm{a}} + \Gamma_{\mathrm{b}} - \sigma_{\mathrm{adh}} - \frac{3}{2} \Gamma_{\mathrm{l}} \frac{2^{1/2}}{3^{3/4}} A_{\mathrm{a}}^{-3/2} \right) \mathrm{d}A_{\mathrm{a}}. \tag{T23}$$

We thus have a pressure which depends on the local tissue height $h$, i.e.

$$P = \Gamma_{\mathrm{a}} + \Gamma_{\mathrm{b}} - \sigma_{\mathrm{adh}} - \Gamma_{\mathrm{l}} \frac{3^{1/4}}{2^{1/2}} h^{3/2}. \tag{T24}$$

Note that the sign of the pressure denotes the energetic change of work performed by the cell, i.e. cells which are compressed compared to their equilibrium shapes yield a negative pressure.

We may now approximate the total pressure by assuming uniform, but not necessarily equal heights in the two subdomains: the amnion ring in $R > R_{\mathrm{amnion}}$ and the compressed inner part with $R < R_{\mathrm{amnion}}$. We denote the total gastruloid radius as $R_{\mathrm{tot}} = 350\,\mu$m. Assuming equal pressure in the subdomains, we have in the ring of amnion cells, which reduce their tension by a factor of $m$,

$$P = (m\Gamma_{\mathrm{a}} + m\Gamma_{\mathrm{b}} - \sigma_{\mathrm{adh}}) - m\Gamma_{\mathrm{l}} \frac{3^{1/4}}{2^{1/2}} (h)^{3/2}, \tag{T25}$$

while in the rest, which is the center part with height $h_{\mathrm{center}}$ in which we assume no changes in tensions and adhesion,

$$P = (\Gamma_{\mathrm{a}} + \Gamma_{\mathrm{b}} - \sigma_{\mathrm{adh}}) - \Gamma_{\mathrm{l}} \frac{3^{1/4}}{2^{1/2}} (h_{\mathrm{center}})^{3/2}. \tag{T26}$$

We assume total volume conservation $V_{tot} = h\pi(R_{\mathrm{tot}}^2 - R_{\mathrm{amnion}}^2) + h_{\mathrm{center}}\pi R_{\mathrm{amnion}}^2$. We can derive the amnion height introducing a relative amnion domain volume $v = V_{\mathrm{amnion}}/V_{\mathrm{center}}$, which implies:

$$h_{\mathrm{center}}(h, R_{\mathrm{amnion}}) = \frac{1}{v} \frac{R_{\mathrm{tot}}^2 - R_{\mathrm{amnion}}^2}{R_{\mathrm{amnion}}^2}\, h. \tag{T27}$$

We solve for the amnion size, i.e.

$$R_{\mathrm{amnion}}(h) = \sqrt{R_{\mathrm{tot}}^2 - \frac{V_{\mathrm{tot}}}{\pi h}\left(\frac{v}{1+v}\right)}. \tag{T28}$$

For a fixed total volume and relative amnion volume, we obtain an implicit equation for $h$, which can be solved numerically,

$$\frac{2^{1/2}}{3^{1/4}}(1-m)\left(\frac{\Gamma_{\mathrm{a}}+\Gamma_{\mathrm{b}}}{\Gamma_{\mathrm{l}}}\right) + m\,h^{3/2} = h_{\mathrm{center}}^{3/2} \tag{T29}$$

Note that the uniform basal adhesion term $\sigma_{\mathrm{adh}}$ drops out. It only influences the total pressure offset and thus the tendency to spread (or shrink) upon removal of the boundary condition. This explains why the exact value of basal adhesion does not impact our prediction for amnion shape in gastruloid discs, as the constraint of constant micropatterned area means that the total basal adhesion energy across the entire gastruloid disc is constant irrespective of shape changes. However, this description still implicitly requires a minimal adhesion strength, otherwise gastruloid discs would not spread over the entire radius of the micropatterns (see below for detailed parameter estimations). The assumption of a constant basal adhesion energy density will be relaxed below.

To compare the model predictions to experiments, we normalize both the effective amnion and the effective embryonic core height by the total average height $\bar{h} = V_{\mathrm{tot}}/(\pi R_{\mathrm{tot}}^2)$, i.e. $h_{\mathrm{center}}/\bar{h} \to h_{\mathrm{center}}$ and $h/\bar{h} \to h$. This is equivalent to normalizing the domain heights by volume, factoring out the variability between gastruloid discs from their total number of cells. We obtain:

$$\begin{aligned} h_{\mathrm{center}} &= \frac{h}{h+(h-1)v}, \\ h_{\mathrm{center}}^{3/2} &= (1-m)(1-\sigma_{\mathrm{t}}) + m h^{3/2}, \end{aligned} \tag{T30}$$

where the normalized background confinement stress,

$$\sigma_{\mathrm{t}} = 1 - \left[\frac{2^{1/2}}{3^{1/4}}\left(\frac{\Gamma_{\mathrm{a}}+\Gamma_{\mathrm{b}}}{\Gamma_{\mathrm{l}}}\right)\frac{1}{\bar{h}^{3/2}}\right], \tag{T31}$$

denotes the tension-based contribution to the background pressure. It quantifies the tissue tensions neglecting adhesion balancing such contractile forces. If $\sigma_{\mathrm{t}}$ is negative, i.e. the tissue is under tension, then a tension decrease ($m < 1$) will yield amnion flattening, as global tension can be relaxed through amnion expansion. This height equation now allows the fitting of the (normalized) amnion height as a function of the overall tension $m$ with the confinement stress as sole fitting parameter (Fig. 2h). Here, we take myosin level as a proxy for $m$, given previous findings that it is the strongest predictor of local tension in tissues [9–11].

To show robustness against potential adhesion energy density reductions, we further consider the case of a linear decrease of the adhesion energy density concomitantly with the decrease of myosin tension, i.e.

$$\sigma_{\mathrm{adh}} \to [\beta_0 + m(1-\beta_0)]\,\sigma_{\mathrm{adh}} \tag{T32}$$

in the outer domain. This yields:

$$h_{\mathrm{center}}^{3/2} = (1-m)(1-\sigma_{\mathrm{t}} - (1-\beta_0)\hat{\sigma}_{\mathrm{adh}}) + m h^{3/2}, \tag{T33}$$

with $\hat{\sigma}_{\mathrm{adh}} = \frac{2^{1/2}}{3^{1/4}}\frac{\sigma_{\mathrm{adh}}}{\Gamma_{\mathrm{l}}}\bar{h}^{-3/2} > 0$. For $\beta_0 < 1.0$, i.e. for a reduction in the amnion adhesion we find an additional contribution to the fitted $\sigma_{\mathrm{t}}$ which has the same dependency on volume (corresponding to $\bar{h}$). It thus serves as a correction to the fitted background stress $-\sigma_{\mathrm{t}}$ from the fitted value $\sigma_{\mathrm{t}} - (1-\beta_0)\hat{\sigma}_{\mathrm{adh}}$, but it does not impede the quality of the fit.

To summarize, we found that in unconfined colonies undergoing the global tension reduction, full wetting can be achieved, which matches the small amnion-induced colonies (Fig. 2i-k). In the full gastruloid discs, such flattening additionally leads to a compression of the embryonic core, opposing the full wetting of the amnion domain. This entails a reduced flattening behavior for similar tension reductions, which is in line with our experimental observation, as will be outlined below (Fig. 2h,k).

#### Parameter estimation and fitting

As discussed in the main text, the ratio of apical to lateral tension $\Gamma_a/\Gamma_l$ can be extracted in a straightforward manner from data via our 3D angle reconstruction [5], and is found to be approximately constant in the amnion domain throughout the time course of shape changes (Fig. 2b). From this, we take $\Gamma_a/\Gamma_l = 2$ in our model. The background stress $\sigma_t$ is obtained from the gastruloid discs as a fitting parameter, and we show in Fig. 2h via Bayesian fitting (Maximum-a-posteriori estimator with flat prior on $\sigma_t \in [-5, 0.5]$) that $\sigma_t = -0.48$ is a good predictor for self-organized gastruloid discs with a 95% confidence interval of $\mathrm{CI} = [-0.55, -0.41]$. This also holds to explain the height change upon the increase of actomyosin tension via lysophosphatidic acid, LPA (Extended Data Fig. 4l).

As the adhesion energy density drops out in the gastruloid discs, we can directly calculate the contractile basal tension from the background tension, Eq. (T31), to be $\Gamma_b/\Gamma_l = 0.61$. From this, we can also extract an implicit minimum bound for basal adhesion by assuming a non-contracting pressure, i.e. $P < 0$ in Eq. (T26), which is $\sigma_{\mathrm{adh}}/\Gamma_l \gtrapprox 0.12$ for $h_{\mathrm{center}} \approx 25\,\mu\mathrm{m}$. As discussed above, its exact value is not required for the prediction as long as we assume that the gastruloid disc covers the entire micropattern radius of $350\,\mu\mathrm{m}$, something which is observed experimentally at the relevant timepoints of self-organization.

For unconfined colonies, we take the same values as above, although basal adhesion now plays a stronger role in dictating the exact magnitude of spreading, beyond the global decrease of tension. To estimate it, we can use the morphology of unstimulated free colonies, where we find an aspect ratio of $h/\sqrt{A} \approx 1$, which means from Eq. (T16) $\frac{\sigma_{\mathrm{act}}}{\Gamma_l} = \frac{\Gamma_b}{\Gamma_l} - \frac{\sigma_{\mathrm{adh}}}{\Gamma_l} \approx -1.07$. This predicts an adhesion strength of $\sigma_{\mathrm{adh}} \approx 1.68$, which is sufficient to localize unconfined amnion cells (but not confined colonies) in the fully wetted regime, assuming the tension reduction to be similar as in confined colonies, i.e. $m \approx 0.4$ (Fig. 2h,k).

#### Numerical cross-sectional bubbly vertex model

Key conclusions from these analytical calculations from the three-dimensional bubbly vertex model can also be verified numerically in a cross-sectional bubbly vertex simulation (Extended Data Fig. 3b). We consider a lateral cross-section through the edges, thus replacing surface tensions by line tensions, while area conservation is implemented through an elastic area contribution in the cell energy, so that the energy of a single cell reads:

$$e_{\mathrm{cell}} = \gamma_a \ell_a - \sigma_{\mathrm{act}} \ell_b + \gamma_l \ell_l + K_A (A - A_0)^2, \tag{T34}$$

with apical and lateral line tensions $\gamma_{a,l}$, basal stress $\sigma_{\mathrm{act}}$, apical, basal and lateral edge lengths $\ell_{a,b,l}$, respectively. The last term corresponds to area conservation $A$ to a reference value $A_0 = 1$ (i.e. the equivalent of volume conservation in 3D), as we take a large value of $K_A = 20$.

While force balance is identical to the three-dimensional bubbly vertex model case, the scaling of tensions and height is slightly different. The equilibrium height in the flat cross-sectional vertex model with $A = 1$, where we neglect the lines' curvatures ($h'' = 0$) reads with $\ell_l = h' = h$:

$$h_{\mathrm{eq}} = \left( \frac{\gamma_a - \sigma_{\mathrm{act}}}{\gamma_l} \right)^{1/2}. \tag{T35}$$

For a confined cross-sectional gastruloid disc, we consider a line of cells, where we fix one lateral interface to a vertical line (the center) and fix the edge cell's outer basal node position. The edge cell is assumed to not have a lateral line at the boundary but rather a contact of the apical and basal line. For simulations of the gastruloid cross-section, we assumed $\sigma_\mathrm{t} = -0.8$, $\gamma_\mathrm{a} = 2$, $\gamma_\mathrm{l} = 1$, $\sigma_\mathrm{adh} = 1.68$, and $\gamma_\mathrm{b} = 0.61$. The myosin-dependent tension reduction with $m = 1.0,\ 0.5,\ 0.25$ is assumed to occur in the outer 3 cells and the tensions are interpolated over 5 cells (Fig. 2d). Note that height changes are less pronounced in the cross-sectional model due to the difference in the scaling between height and tensions, Eq. (T35).

To consider the three-dimensional cross-sectional geometry of the *in vivo* amniotic sac, we consider a circular arrangement of cells (Extended Data Fig. 4m), which enclose a lumen with prescribed volume (with a similar quadratic area elasticity as the cells). Contrary to the previous geometry, which was constrained to a flat basal surface, basal vertices can move freely in a circular geometry, so that we additionally assume a stabilizing contribution of the extracellular matrix, which prevents unphysical cellular deformation due to basal adhesion. This is a reasonable assumption, since previous work showed that ECM surrounds the epiblast at peri-gastrulation [12]. We introduce this ECM constraint by considering only two basal nodes per cell and adding a bending energy contribution,

$$E_\mathrm{bending} = \kappa_\mathrm{B}\left(1 + \cos(\vartheta)\right), \tag{T36}$$

with bending rigidity $\kappa_\mathrm{B}$ and (interior) angle between neighboring basal edges $\vartheta$. We find that $\kappa_\mathrm{B} = 3$ is sufficient to fully stabilize the system and give rise to realistic smooth shapes. Note that we also prevent unphysical interpenetration of vertices by an additional hard-core repulsion between apical or basal nodes, via an intracellular potential of

$$U_\mathrm{repulsion}(d) = \begin{cases} \mathcal{E}\left[\left(\frac{d_0}{d}\right)^4 - 2\left(\frac{d_0}{d}\right)^2 + 1\right] & \text{if } d < d_0 \\ 0 & \text{else} \end{cases}, \tag{T37}$$

with apico-basal distance $d$, and parameters $d_0 = 0.1$ and $\mathcal{E} = 10$, is used, which contributes as an additional node-specific force.

As detailed above, we found experimentally that tension in the amnion is reduced (Fig. 2f,g,l, Extended Data Fig. 4i). We thus hypothesized that the amnion flattening observed both *in vivo* and in amniotic sac embryoids [13] could in principle be achieved either through increased luminal pressure (which preferentially stretches the soft amnion cells), or by the same wetting mechanism that we found in 2D gastruloid discs. Both potential models for amnion flattening were implemented with an inflated volume of $V_\mathrm{inflated}/V_\mathrm{initial} = 6$ for the pressure-driven case and $V_\mathrm{inflated}/V_\mathrm{initial} = 2$ for the wetting case (the latter to accommodate additional surface area). In both models the relative contractile tension in the amnion is lowered homogeneously across apical, basal and lateral interfaces, i.e.

$$\gamma_\mathrm{a,b} \to m \cdot 2.0, \qquad \gamma_\mathrm{l} \to m \cdot 1.0, \tag{T38}$$

with relative myosin reduction $m$ and $\sigma_\mathrm{act} = \sigma_\mathrm{adh} - \gamma_\mathrm{b}$. For wetting we additionally consider a constant basal adhesion energy density of $\sigma_\mathrm{adh} = 0.7$, which vanishes in the other case.

In the first scenario, we find that pressure-driven stretching of a softer amnion leads to larger curvature in the amnion-domain (Extended Data Fig. 4o, with $m = 0.15$). This is consistent with the Young-Laplace law which predicts a larger curvature from pressure for smaller tension in the amnion domain. Strikingly, in the second scenario of amnion wetting, we observe the opposite behavior, where curvature is localized in the epiblast (Extended Data Fig. 4n, with $m = 0.15$). Effectively, wetting leads to a negative spontaneous curvature, which resists the bending of the amnion and thus leads to a less-curved amnion domain [14]. Interestingly, and as discussed in the main text, the latter model thus gives a good agreement with the *in vivo* shape of the amniotic sac with its slit shape, flattened amnion, and decreased curvature of the amnion. In contrast, pressure-mediated flattening yields a symmetric cavity shape, which does not agree with the shape observed *in vivo* or in 3D *in vitro* models of the amniotic sac.

### 2. Mechanochemical model of the keratin-myosin toggle switch motif

#### Double-negative direct feedback model

We assume a minimal feedback model between keratin levels $k$ and myosin levels $m$, which are subject to a double negative feedback loop.

Conservation equations for the system read:

$$\begin{aligned} \partial_t m &= a_m \frac{\tilde{k}^n}{\tilde{k}^n + k^n} - b_m m, \\ \partial_t k &= a_k \frac{\tilde{m}^n}{\tilde{m}^n + m^n} - b_k k, \end{aligned} \tag{T39}$$

where $a_{m,k}$ quantify activation strengths of each species (which is modulated by the mutual feedback), $b_{m,k}$ are degradation rates of each species (effectively setting the timescales of the system), $\tilde{m}$ and $\tilde{k}$ are relative inflection points of the inhibition and $n$ is the Hill coefficient of the negative feedback interactions.

As we mainly study stable final states, we can set the rates $b_{m,k}$ to 1 without loss of generality. Given that we normalize maximal concentrations so that all concentrations are bounded between 0 and 1, we can generically assume that the inflection point of the feedback is at an intermediate value, and took $\tilde{k} = \tilde{m} = 0.75$; although, as discussed below, varying parameters around this value does not change qualitatively the landscape. This leaves the strength of feedback on activation $a_{m,k}$ as the main adjustable parameter.

We first study for simplicity fully symmetric negative feedbacks, $a_{m,k} = a$. As long as $n >= 2$, the system generically displays bistability and sharp transition in states above a critical value $a > a_c$, i.e. strong enough mutual inhibitory feedback (Extended Data Fig. 7a-i), characteristic of a toggle switch [15, 16]. Below this critical value of $a$, this system of equations is monostable, leading to intermediary concentrations of both myosin and keratin across conditions, which is inconsistent with our data (Extended Data Fig. 7a,d,g). Thus, in the following, we take $n = 3$ (at which $a_c \approx 0.9$) but stress that the key results are robust to different parameter choices. The bifurcation diagram showing this bistability and the corresponding phase portrait around this choice of parameter are depicted in Extended Data Fig. 7d-f. Varying the asymmetric coupling strength $\Delta a = a_m - a_k$ while keeping the average negative feedback coupling $a = (a_m + a_k)/2$ fixed leads to loss of bistability through a saddle node bifurcation of one of the solutions. Changing these parameters through perturbations thus allows for rapid switching between different states, when perturbing into the regime outside the stability region of a steady state.

To confirm the two-attractor nature of the landscape, we consider final solutions starting from different initial states in the expected myosin and keratin range $m, k \in [0.05, 1.7]$ to probe the existence of fixed points. In the range of $a_m, a_k \in [0.5, 1.3]$, $n \in [2, 4]$, and $b_m, b_k \in [0.9, 1.1]$, we integrated the system of equations over time and obtained the density of final states as a representation of the statistical landscape of reachable states by the system (Extended Data Fig. 7k). As expected from a toggle switch system, this landscape displays two distinct attractors (i.e. low myosin and high keratin, or high myosin and low keratin), showing the robustness of the phenomenology. Note that more complex network topologies, for instance where keratin and myosin do not interact directly with each other, but indirectly via other cues (e.g. cell shape, YAP1) are possible but yield qualitatively similar results in terms of landscapes, as we discuss in the final section of this Theory Note.

Simple analytical arguments could further guide our parameter search. For instance, the equation for equilibrium myosin reads: $a_m \frac{\tilde{k}^n}{\tilde{k}^n + k^n} = m$, which in the low keratin state and for sharp Hill coefficient simplifies to $m \approx a_m$. Similarly, the equation for equilibrium keratin $a_k \frac{\tilde{m}^n}{\tilde{m}^n + m^n} = k$ simplifies in the low myosin state to $k \approx a_k$. Given that we have normalized the data so that concentrations are bounded between 0 and 1, this means that the strength of the couplings must be of order unity but lower than 1. Similarly, we expect Hill thresholds to be intermediary between 0 and 1. Qualitatively, a manual parameter search suggested that $\tilde{m} = \tilde{k} = 0.75$ and $a_m = a_k = 0.7$ reproduced key features of the data. However, we see for instance that the range of myosin levels seen in the data is narrower (0.4-0.9) than the range of keratin (0.1-1), which cannot occur with purely symmetric and equal coefficients.

We therefore took a more unbiased approach, performing least-square fitting of the entire dataset across conditions shown in Fig. 3h-n, and Extended Data Fig. 6a-f,l-o, keeping $n = 3$ and optimizing for remaining parameters. We found the best fit parameters as $\tilde{m} = 0.75$, $\tilde{k} = 0.66$, $a_m = 0.8$ and $a_k = 0.55$, close to the manual inspection of parameter space (see below for other parameters for specific conditions). For these parameters without additional perturbations we only have one stable fixed point with high myosin and low keratin (Extended Data Fig. 7j,k,l), representing the unstimulated state.

#### Unperturbed dynamics upon BMP4 stimulation

In gastruloid discs, the addition of exogenous BMP4 leads to activation of BMP signaling at the edge, where the amnion cells will form and undergo a switch toward a keratin-based organization. This leads to a change in keratin production in the model, by a BMP-dependent factor $b$:

$$\partial_t k = (1 + b)a_k \frac{\tilde{m}^n}{\tilde{m}^n + m^n} - b_k k$$

While for low $b$, the model predicts that the gastruloid edge remains in a myosin-high, keratin-low state (mirroring unstimulated conditions), an increase of $b = 0.8$ leads to a switch towards a myosin-low, keratin high state (Extended Data Fig. 7l), at a timescale controlled by $b_k$ and $b_m$. Importantly, a generic feature of this type of toggle switch topology is that the path from one attractor to the other should follow a straight line—or anti-diagonal axis in (myosin,keratin) space—meaning that myosin downregulation occurs simultaneously with keratin up-regulation. This mirrors the dynamics shown in Fig. 3l, where unstimulated micropatterned colonies remain in a high myosin/low keratin state even at 48h, while stimulated colonies reach the low myosin/high keratin state.

#### Experimentally perturbing the cytoskeletal switch

Withaferin-A (WFA) was previously shown to trigger intermediate filament disassembly [17–19] and can thus induce a depolymerization of keratin. This maps in our model to an increase in $b_k$. Introducing this keratin inhibition in addition to the BMP stimulation (increasing $b_k$ for $b = 0.8$) prevented the switch toward a high keratin state, therefore stabilizing the initial high myosin state (Fig. 3n). More quantitatively, $b_k = 1.085$ was sufficient to reproduce the experimental data upon WFA treatment. Inhibiting keratin expression via small interfering RNAs (siRNA K8/K18) also agreed well with the theoretical prediction (Fig. 3n, Extended Data Fig. 7m). Note that due to the asymmetry in the model couplings ($a_m > a_k$), perturbations along the keratin axis only show weak to no bistability, as shown in the landscape of Fig. 3n.

Lysophosphatidic acid (LPA) has a well-established effect in stimulating myosin contractility [20]. This translates in our model in an increase of myosin activation by a factor $L$:

$$\partial_t m = La_m \frac{\tilde{k}^n}{\tilde{k}^n + k^n} - b_m m$$

.

Modeling this pharmacological perturbation together with the BMP treatment also abolishes the switch above a critical value of $L$, and thus that amnion cells should remain in a myosin-high, keratin-low state (Extended Data Fig. 7n). Our findings with high-doses of LPA (50$\mu$M), as seen in Fig. 3m, where we took $L = 1.08$ suggest that two attractors may co-exist in this system. This is a very generic prediction of the toggle switch logic. More specifically for our parameter regime, although parameters for high concentrations of LPA favor the myosin high attractor, the model generically predicts that lower doses of LPA (intermediate

$L$) should lead to a region where two attractors are near-equally probable, and thus where we expect the outcome to be dependent on noise and exact parameter values or initial conditions. Note that this toggle-switch behavior is much stronger when modifying myosin levels in the model (compared to modifying keratin levels), due to our current parametrization: $a_m > a_k$ effectively means that actomyosin is "stronger" than keratin in the double-negative feedback loop, and thus deeper in the bistable/toggle-switch region.

#### Cell area variability from coupling to mechanics

To quantify the variability in cellular states in the bistable regime for lower doses of LPA (Fig. 3o), we consider the mechanical effect of variability along the myosin-keratin switch axis. In experiments, intra-gastruloid variability was computed based on apical area segmentation of amnion cells (defined as elsewhere in the manuscript as all cells within $50\mu m$ of the gastruloid edge). Each gastruloid was divided into 10 angular sectors (we verified that the findings were not sensitive to this choice). We then computed the average and coefficient of variation of cell area across different sectors, which is a direct read-out of the spatial variability of cell area within a given gastruloid disc (Fig. 3p).

To match this quantification theoretically, we simulated 1000 gastruloid discs, each with 1000 amnion cells per disc according to the keratin-myosin equations described above. All cells in a given gastruloid disc share the same value of LPA activation $L$ (taken from a uniform distribution between 1 and 1.2, i.e. parameters around the best-fit values for wild-type and LPA conditions), but have also a small cell-intrinsic difference in $a_m$ (the exact value does not qualitatively change the results, we took a dispersion of 1% around the best-fit value). To translate myosin to area, we used our analytical model from section 1, where $\sigma_t$ is the key parameter, and also picked the value of $\sigma_t$ for each gastruloid disc from a random distribution between -0.5 and -2 (i.e. around the best fit value described above). This generates a range of amnion domains with different average areas and strength of myosin activation. Given the toggle switch logic of the model (Fig. 3k), this generically gives rise to a region of maximal variability for intermediate average areas (Extended Data Fig. 7o), mirroring the data.

Note that this analysis neglects spatial couplings in the system. From a mechanical perspective, the coexistence of highly columnar cells (myosin high) next to highly squamous cells (keratin high) would create an additional energy penalty. Yet, we do not expect this to change qualitatively the features discussed above, as this simply introduces a correlation length in the system, providing an explanation for why squamous cells in the 20$\mu$M LPA condition form compact domains (Fig. 3o). In the context of a mosaic switch, for instance in siRNA K8/K18 or TRULI (see below), we expect a similar structure for the keratin-myosin switch, but with dampened correlation between myosin and area due to cell-cell interactions.

#### Incorporating YAP1 in the double-negative feedback model

Our results, discussed in the main text, on the influence of YAP1 on the morphogenesis of amnion cells suggest that YAP1 is not simply a downstream sensor of cell shape, but is involved in the keratin-actomyosin switch (Fig. 4h-k). This is consistent with YAP1 inhibition leading to a perturbed actomyosin-keratin switch (Fig. 4h, Extended Data Fig. 8e,f), yielding a final state with higher myosin and lower keratin compared to stimulated gastruloid discs. Furthermore, we found that activation of YAP1 in the absence of BMP4 stimulation using TRULI is sufficient to induce the cytoskeletal switch (Fig. 4k).

In this section, we therefore consider all the possible minimal extensions of our model incorporating YAP1, which can still reproduce all of our experimental conditions. As shown in a number of other contexts [21], and by our observations that YAP1 activity is tightly linked to cell shape throughout perturbations (i.e. when the squamous morphogenesis is perturbed, YAP is less enriched in the nucleus), we consider in all of these models that YAP1 activity $y$ is positively regulated by cell spreading, and thus from our bubbly vertex model results, negatively regulated by global myosin levels. Note that this assumes full mechanical relaxation of height and the concomitant increase in apical area, which will be suppressed in mosaic states with different cytoskeletal states interspersed, as discussed above.

As to the effect of YAP1 on keratin and myosin, we found that there are three possible minimal network topologies that can qualitatively reproduce the YAP1 inhibition (Fig. 4h) and activation (Fig. 4k) data: i) YAP1 activates keratins, and keratin inhibits myosin (without the need for a direct arrow from myosin to keratin), ii) YAP1 inhibits myosin, and keratin promotes spreading, which further reinforces YAP1 nuclear enrichment (without the need for a direct arrow from keratin to myosin), or iii) a fully indirect model where YAP1 both activates keratins and inhibits myosin, and keratin/myosin regulate YAP1 via cell spreading (without any direct arrows between myosin and keratin).

In model 1, the negative feedback of myosin to keratins goes through YAP1, which implies for the dynamic equations for keratin $k$, YAP1 $y$, and myosin $m$:

$$\begin{aligned} \partial_t k &= -b_k k + c_k y, \\ \partial_t y &= a_y \frac{\tilde{m}^n}{\tilde{m}^n + m^n} - b_y y, \\ \partial_t m &= a_m \frac{\tilde{k}^n}{\tilde{k}^n + k^n} - b_m m, \end{aligned} \tag{T40}$$

where we assumed a reference value of $k = 0$, neglecting production terms. Changes in YAP1 cytoplasmic to nuclear localization usually occurs on the timescales of minutes [22, 23], while the developmental processes we consider occur over hours. We can thus consider fast YAP1 dynamics, which implies $b_y, a_y \to \infty$ while $a_y/b_y = \text{const.}$ and $y = \frac{a_y}{b_y} \frac{\tilde{m}^n}{\tilde{m}^n + m^n}$. In this limit, the model exactly reduces to the equations of the minimal negative feedback loop, simply with rescaled parameters.

In model 2, the negative feedback from keratin to myosin goes through YAP1, meaning that the equations for the system read:

$$\begin{aligned} \partial_t k &= a_k \frac{\tilde{m}^n}{\tilde{m}^n + m^n} - b_k k, \\ \partial_t y &= a_y \frac{\tilde{m}^n}{\tilde{m}^n + m^n} - b_y y + c_y k, \\ \partial_t m &= a_m \frac{\tilde{y}^n}{\tilde{y}^n + y^n} - b_m m. \end{aligned} \tag{T41}$$

Again, the assumption of fast YAP1 dynamics can be made and the limit $a_y, b_y, c_y \to \infty$ with $a_y/b_y = \text{const.}$ and $c_y/b_y = \text{const.}$ yields:

$$y = \frac{c_y}{b_y} k + \frac{a_y}{b_y} \frac{\tilde{m}^n}{\tilde{m}^n + m^n}. \tag{T42}$$

The reduced full dynamical equations then read:

$$\begin{aligned} \partial_t k &= a_k \frac{\tilde{m}^n}{\tilde{m}^n + m^n} - b_k k, \\ \partial_t m &= a_m \frac{\tilde{y}^n}{\tilde{y}^n + (\frac{c_y}{b_y} k + \frac{a_y}{b_y} \frac{\tilde{m}^n}{\tilde{m}^n + m^n})^n} - b_m m. \end{aligned} \tag{T43}$$

The first equation for the keratin state is unchanged compared to the direct model, while the second equation for the myosin state has a similar negative feedback structure with feedback from the keratin state, where the effective keratin level is influenced by an additional myosin self-regulatory correction. In the case of $(a_y/b_y) \ll (c_y/b_y)$ this correction is negligible (at leading order for small $a_y/c_y$) and we can recover the direct minimal feedback model with rescaled parameters. This limit requires a strong coupling of cell spreading and concomitant YAP1 activation and keratin, consistent with our experiments where YAP1 activation, even in the absence of BMP4, is sufficient to result in keratin upregulation (Fig. 4k).

In Model 3, we consider a fully indirect model with a negative feedback loop between myosin and YAP1 and a positive feedback loop between keratin and YAP1 without any direct interaction of keratin network and myosin level. Here the equations read:

$$\begin{aligned}
\partial_t k &= a_k y - b_k k, \\
\partial_t y &= a_y \frac{\tilde{m}^n}{\tilde{m}^n + m^n} - b_y y + c_y k, \\
\partial_t m &= a_m \frac{\tilde{y}^n}{\tilde{y}^n + y^n} - b_m m.
\end{aligned} \tag{T44}$$

We can reduce it to a model with similar qualitative behavior as the direct negative feedback in the case of fast YAP1 dynamics. As in model 2, the limit $a_y, b_y, c_y \to \infty$ with $a_y/b_y =$ const. and $c_y/b_y =$ const. yields again:

$$y = \frac{c_y}{b_y} k + \frac{a_y}{b_y} \frac{\tilde{m}^n}{\tilde{m}^n + m^n}. \tag{T45}$$

The reduced full dynamical equations read:

$$\begin{aligned}
\partial_t k &= \left(\frac{a_k a_y}{b_y}\right) \frac{\tilde{m}^n}{\tilde{m}^n + m^n} - \left(b_k - \frac{a_k c_y}{b_y}\right) k, \\
\partial_t m &= a_m \frac{\tilde{y}^n}{\tilde{y}^n + \left(\frac{c_y}{b_y} k + \frac{a_y}{b_y} \frac{\tilde{m}^n}{\tilde{m}^n + m^n}\right)^n} - b_m m.
\end{aligned} \tag{T46}$$

Again, the first equation for keratin is identical to the corresponding equation in the direct feedback loop, simply with rescaled parameters, as long as $b_k$ is large enough so that the effective degradation rate of keratin is always positive (required for system stability). The second equation for myosin is identical to the one from Model 2, and the same limit can therefore recover the toggle switch architecture of the direct model, with rescaled interaction parameters.

Overall, we find that incorporating YAP1 in the model can lead to qualitatively similar outcomes as the direct keratin-myosin toggle switch phenomenology that we studied above, in the limit for fast YAP1 dynamics, simply introducing additional model parameters. By construction of how we constrained network topologies, all of these models reproduce the experimental findings that myosin activation (LPA) and perturbing the keratin network, either via WFA or KRT8/18 downregulation, resulted in lower YAP1 activity in the amnion, but also that YAP1 inhibition (resp. activation) can prevent (resp. trigger) the myosin-keratin switch (Fig. 4h,k). We therefore report in the main text and figures results from the direct toggle-switch model from a principle of parsimony.

### Theory References
